# Uniquome: Construction and Decoding of a Novel Proteomic Atlas that Contains New Peptide Entities

**DOI:** 10.1101/2024.01.30.577925

**Authors:** Evangelos Kontopodis, Vasileios Pierros, Constantinos E. Vorgias, Issidora S. Papassideri, Dimitrios J. Stravopodis, George Th. Tsangaris

**Affiliations:** Proteomics Research Unit, Biomedical Research Foundation of the Academy of Athens (BRFAA); Athens, 11527, Greece; Section of Cell Biology and Biophysics, Department of Biology, School of Science, National and Kapodistrian University of Athens (NKUA); Athens, 15701, Greece; Section of Biochemistry and Molecular Biology, Department of Biology, School of Science, National and Kapodistrian University of Athens (NKUA); Athens, 15701, Greece

**Author notes:** Corresponding author; (G.Th.T.).

## Abstract

Cellular and molecular uniqueness has recently gained eminent importance, due to the large amount of data produced by “-omics” technologies. Herein, we have constructed and decoded the “**Uniquome**”, by introduction of the new peptide entities: (a) “**Core Unique Peptide**” (CrUP), defined as the peptide whose sequence is accommodated, specifically and exclusively, only in one protein in a given proteome, and also bears the minimum length of amino acid sequence; (b) “**Composite Unique Peptide**” (CmUP), defined as the peptide composed by the linear unification of CrUPs, when two or more successive in order CrUPs overlap one another; (c) “**Family Unique Peptide**” (FUP), defined as the CrUPs that are common between all members of a given family, but unique only for the protein members of the particular family, and (d) “**Universal Unique Peptides**” (UUPs), which are the common CrUPs in a given protein across organisms, carrying the important ability to securely identify a protein independently of an organism. By these entities as tool-box, we have analyzed the human and model organisms, respective, proteomes. We demonstrate that these novel peptide entities play a crucial role for protein identification, protein-function prediction, cell physiology, tissue pathology, therapeutic oncology and translational medicine. Finally, we suggest that across species the conserved sequences are not DNA nucleotides but CrUPs entities.

**One-Sentence Summary:** We constructed and decoded the “Uniquome”, by introducing the new peptide entities Core Unique Peptide, Composite Unique Peptide, Family Unique Peptide and Universal Unique Peptide

## Introduction

Cellular and molecular uniqueness has gained eminent importance, due to the large amount of data produced by genomic and proteomic technologies [Next Generation Sequencing (NGS) and Mass Spectrometry (MS), respectively]. At the genomic level, unique nucleotide sequences, within the human genome, have widely used in the past for design of PCR primers and for construction of microarray platforms, while unique nucleotide sequences have proved to play essential roles in epigenetic regulation, chromatin status alterations, transcriptional dynamics and post-transcriptional modifications in different ways (1, 2). Recently, a new group of unique nucleotide sequences, specific and tailor-made for each person, known as singleton, have been identified (3). Likewise, Quasi-prime peptides, which are the shortest peptide sequences, unique to a species, have been also detected (4).

Large-scale collaborative efforts have caused a dramatic increase in the number of identified proteins being derived from the human proteome, thereby leading to serious and extensive implications of proteomics technology in human health, disease diagnosis and therapy, and understanding of diverse biological processes. A critical issue for proteomics-driven illumination of molecular mechanisms controlling cellular pathology relies upon each protein’s unambiguous and accurate identification. Hitherto, the prevalent proteomics technology for protein identification exclusively depends on MS, during which a given protein is cleaved to peptides that are detected by a mass spectrometer, producing a mass spectrum, which is, next, analyzed computationally to finally result in each protein’s identification profile (5,6). Hence, differential protein identification, in the context of a whole proteome, is an entirely peptide-related process. This peptide-protein association unveils the major importance of peptides’ amino acid sequence characterization to the fidelity and accuracy of cognate protein’s identification, thereby strongly suggesting that the amino acid sequence of peptides being uniquely carried by a given protein can ensure its confident identification.

Towards this direction, we have, herein, attempted to construct, for the first time, the human “**Uniquome**”, by primarily defining the term of “**Unique Peptide**” (UP) as the peptide that is being recognized specifically and solely in only one protein within a given proteome. In a previous study, we have determined the human proteome-containing UPs by their molecular weight, in order to identify each analyzed protein by a MS-based platform, known as “Peptide Finder” (PF) (7). Despite its broad use, PF patterning contains an intrinsic disadvantage of not considering the position of an amino acid in the sequence of a given peptide. Therefore, peptides with identical amino acids in their sequence, but in different positions within their cognate proteins, are detected with the same molecular weight, but they cannot be separately and distinctively considered (7, 8).

Besides UP, we, herein, also introduce two completely new peptide entities: the “**Core Unique Peptide”** (CrUP) and the “**Composite Unique Peptide**” (CmUP). We define CrUP as the peptide whose sequence is accommodated, specifically and exclusively, only in one protein - in a given proteome-, while it also bears the minimum length of amino acid sequence. To the contrary, a typical CmUP is composed by the linear unification of CrUPs, when two or more successive, in order, CrUPs overlap one another (Figure S1). It is self-evident that, independently of its length, each peptide, within a given protein, containing a CrUP is also considered to carry a UP.

Ultimately, the complete set of UPs, embracing all the identified CrUP and CmUP peptide entities, can altogether structure the Uniquome of an organism. In the context of a Uniquome, additional, new, entities are also, herein, introduced: the density of CrUPs (“d-CrUPs”) and the density of CmUPs (“d-CmUPs”). d-CrUPs is defined as the ratio of total number of CrUPs to the total amino acid length of all proteins in an organism, while d-CmUPs is described as the ratio of total number of CmUPs, in each organism, to the total amino acid length of its proteins. The ratio of total number of amino acids that reside in UPs, of a given proteome, to the total amino acid length of all its proteins is, herein, defined as “Unique Coverage” (UC).

The purpose of the present study is the construction of human Uniquome, and the investigation of CrUP- and CmUP-derived molecular roles in diverse cellular functions. Demonstration of CrUP- and CmUP-dependent applications in human proteome activities, during pathology, development, and evolution, has been, also, attempted. To, further, provide an evolutionary perspective to Uniquome, and its, new, peptide entities, CrUPs and CmUPs, we have expanded our study to major model organisms and their proteomes, across species.

## Results and Discussion

### Construction of the Human Uniquome

20,422 reviewed human proteins, all derived from the Uniprot database (version 05-2023), were analyzed for the presence of CrUP and CmUP, novel, peptide entities. Following the procedure described in Material and Methods, a total of 1,004,442,267 peptides were generated containing 50,690,065,140 amino acids (47.2 GB data). By these peptides 7,271,607, from 4 to 100 amino acids (aa) in length, were characterized as CrUPs, in 20,258 human proteins (Table 1), whereas 164 proteins (0.80%) were not found to contain any UP species of the same length (4 to 100 aa) (Table S1). Further analysis revealed that the 7,271,607 CrUPs could generate 66,447 CmUPs. The total density of human proteome in CrUPs is 64%, indicating that 64 CrUPs are being generated per 100 aa (in length), while the coverage of human proteome by UP species is 93% (Table 1). As an example, TITIN (Q8WZ42), the largest Human protein with 34,350 aa lengths, a total of 3,327,003 peptides (4 to 100 aa) were generated and by these 23,706 (0.69%) were characterized as CrUPs.

**Table 1.** The human Uniquome.

| Proteins (reviewed) | 20,422 |
| --- | --- |
| Core Unique Peptides (CrUPs) | 7,271,607 |
| Composite Unique Peptides (CmUPs) | 66,447 |
| Density of CrUPs | 64% |
| Density of CmUPs | 0.59% |
| Coverage | 93% |
| Proteins with Unique Peptides (UPs<sup>+</sup>) | 20,258 |
| Proteins without Unique Peptides (UPs<sup>-</sup>) | 164 |

164 out of the 20,422 proteins that belong to the human proteome proved to lack CrUP species (Table S1). 7 out of these 164 proteins were shown to contain -only-2 (P0DPR3) to 24 aa (in length). Although they are registered in the Uniprot database, likely due to their remarkably small sizes, their aa sequences could be surprisingly recognized in other larger (in size) proteins. The remaining 157 proteins were divided in two groups. The first one includes proteins sharing aa homologies -among them-higher than 97%, while the second group contains proteins that are considered as (proteolytic) fragments of bigger proteins (or, splice-variant products). The Q9UBX2, P0CJ85, P0CJ86, P0CJ88, P0CJ89 and P0CJ90 proteins represent characteristic examples of the first group. Notably, they exhibit a homology of 99.76% -among them- and, thus, none of their respective peptides can be unique (since they belong to all of them) (Figure S2). Another example is the group of P0C7V4, P0DV73, P0DV74, P0DV75 and P0DV76 proteins. Remarkably, they share homologies from 97.63% to 100% and, thus, they are unable to include any CrUP species (Figure S3). In the second group of proteins that lack CrUP species, we have classified the P63126 and P63127 proteins (carrying 666 and 156 aa, respectively), which are -proteolytic-fragments of the -bigger-P63128 protein (1,117 aa). Of note, P63126 and P63127 are missing CrUP species, because their respective peptides, also, belong to the full-length -bigger-P63128 protein. P63128 has proved to accommodate CrUPs only in the areas of its sequence that are not common with the (P63126 and P63127) - proteolytic-fragments (667-780 and 936-1,117 aa) (Figure S4).

### Dissection of the Human Uniquome

Analysis of CrUP structure in relation to their length (number of aa they are composed of) showed that the majority (69.03%) of human CrUPs consist of 6 aa, while 20.6 % and 8.62% of CrUPs are characterized by 5 and 7 aa, respectively (Figure 1A and Table S2). 759 CrUPs are composed of 4 aa, while the biggest (in size) CrUPs are detected to contain 100 aa, respectively belonging to the Keratin-associated protein 2-1 (Q9BYU5), Gamma-aminobutyric acid receptor subunit alpha-1 (P14867) and Putative WAS protein family homolog 4 (A8MWX3).

**Figure 1.**
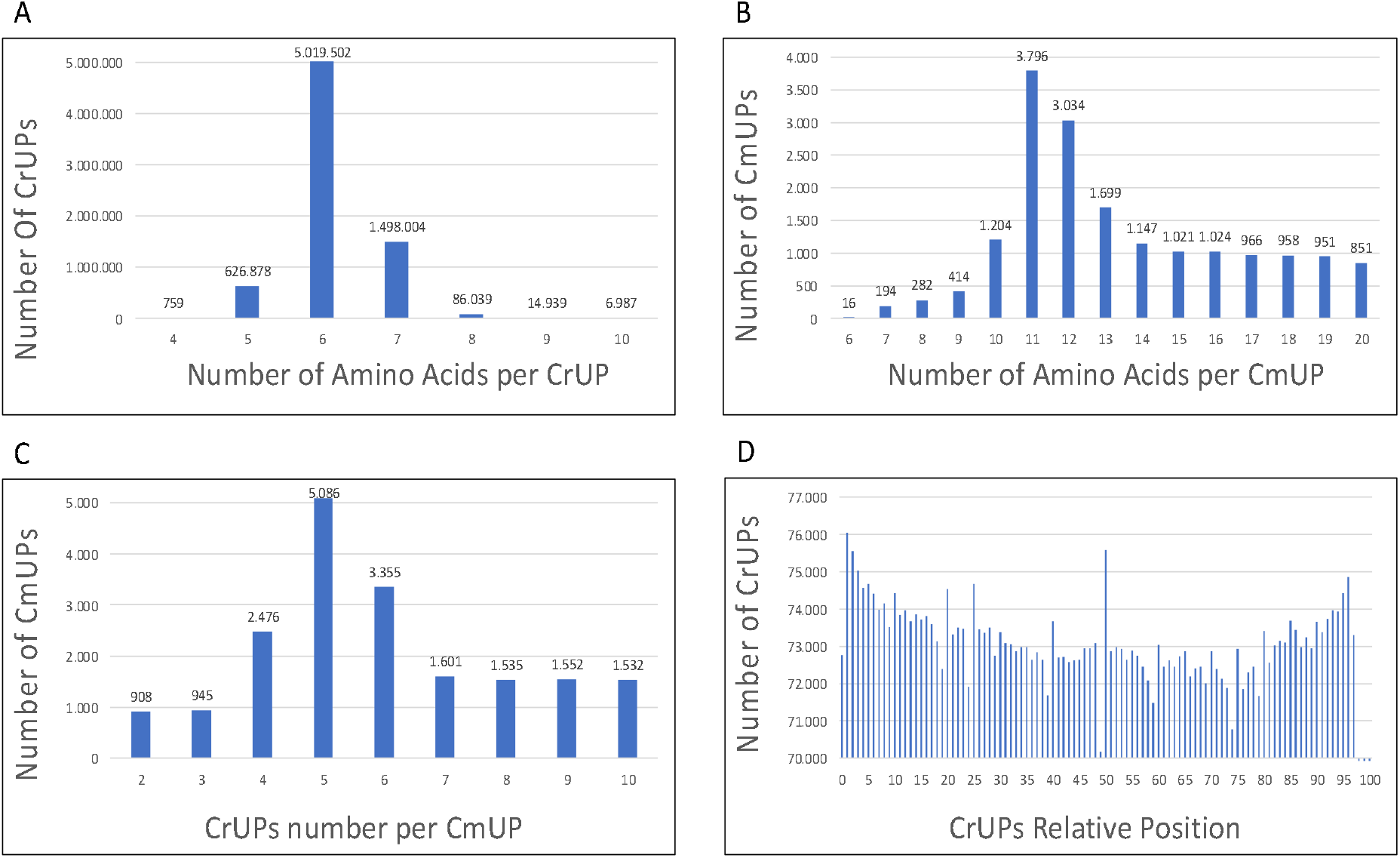
Analysis of the human Uniquome. (**A**) Number of CrUPs in relation to their number of aa they are composed of. The majority of CrUPs consist of 6 aa. (**B**) Number of CmUPs, in relation to their number of aa they are composed of. 11-peptides and 12-peptides represent the higher CmUP groups. (**C**) Number of CrUPs that construct a CmUP. The highest percentage of CmUPs is structured by 5 CrUPs. (**D**) Relative position of CrUPs in proteins. An accumulation of CrUPs are detected exactly in the molecular center of proteins, as well as in their respective carboxy- and amino-terminus.

Similar mapping of the human Uniquome for CmUP species, unveiled that 11-peptides and 12-peptides strikingly represent the higher CmUP numbers (5.71% and 4.57%, respectively), being followed by 10-peptides and 13-peptides with comparatively lower values (1.81% and 2.56%, respectively) (Figure 1B and Table S3). Of note, 908 CmUPs are composed of 2 CrUPs, while the biggest -in length-CmUP belongs to TITIN (Q8WZ42). The highest percentage of CmUPs (7.65%) is structured by 5 CrUPs, while 6 and 4 CrUPs are responsible for the 5.05% and 3.73% of CmUPs, respectively (Figure 1C and Table S4).

### Position Analysis of CrUPs and CmUPs

To evaluate the position of CrUP species within the protein sequences of human proteome, we have, herein, defined as the relative position of each CrUP the position of the first aa of a CrUP to the total aa number of its cognate protein (%). To exemplify the UP-position profiling, the relative position of a CrUP starting at the aa 30 of a protein that bears 250 aa is 12%. Remarkably, we found that the accumulation of CrUPs is detected at the relative positions 1 and 95% (carboxy- and amino-terminus, respectively), and at the position 50%, exactly in the molecular center of a protein (Figure 1D). However, CmUPs are distributed almost equally within all the relative positions of human proteins, in the context of their assembled proteome (Figure S5).

### Chromosomal Distribution of CrUPs

For the distribution profiling of CrUPs onto human chromosomes, we examined the coverage of each chromosome by CrUP species (Table S5). As shown in Figure 2A, we found that all chromosomes exhibit CrUP-coverage values from 90% to 96%, while the mitochondrial chromosome presents 100% -respective-coverage, whereas the Y sex-chromosome is characterized by the surprisingly lowest CrUP coverage of 32%. These findings indicate the importance of chromosomal conservation during molecular evolution, without loss of each chromosome’s UP-based individuality (11, 12), while, on the other hand, strongly suggest a structural divergence and functional uniqueness of the Y sex-chromosome (13, 14). In accordance, mapping of the distribution landscape containing proteins that lack CrUP species of human chromosomes revealed that the Y sex-chromosome accommodates the highest percentage of CrUP-missing proteins (15.8%), whereas all the proteins of mitochondrial chromosome are presented to contain CrUP species (Figure 2B). Intriguingly, our data show that the human chromosomes 7, 8, 13 and X are characterized by the highest percentage of proteins that do not carry CrUP species. Altogether, it seems that mitochondrial proteins possess an unprecedented uniqueness compared to the rest of human proteome contents, mechanistically coupling mitochondrial Uniquome (mtUniquome) with organelle’s evolutionary origin and functional significance for human life (15, 16).

**Figure 2.**
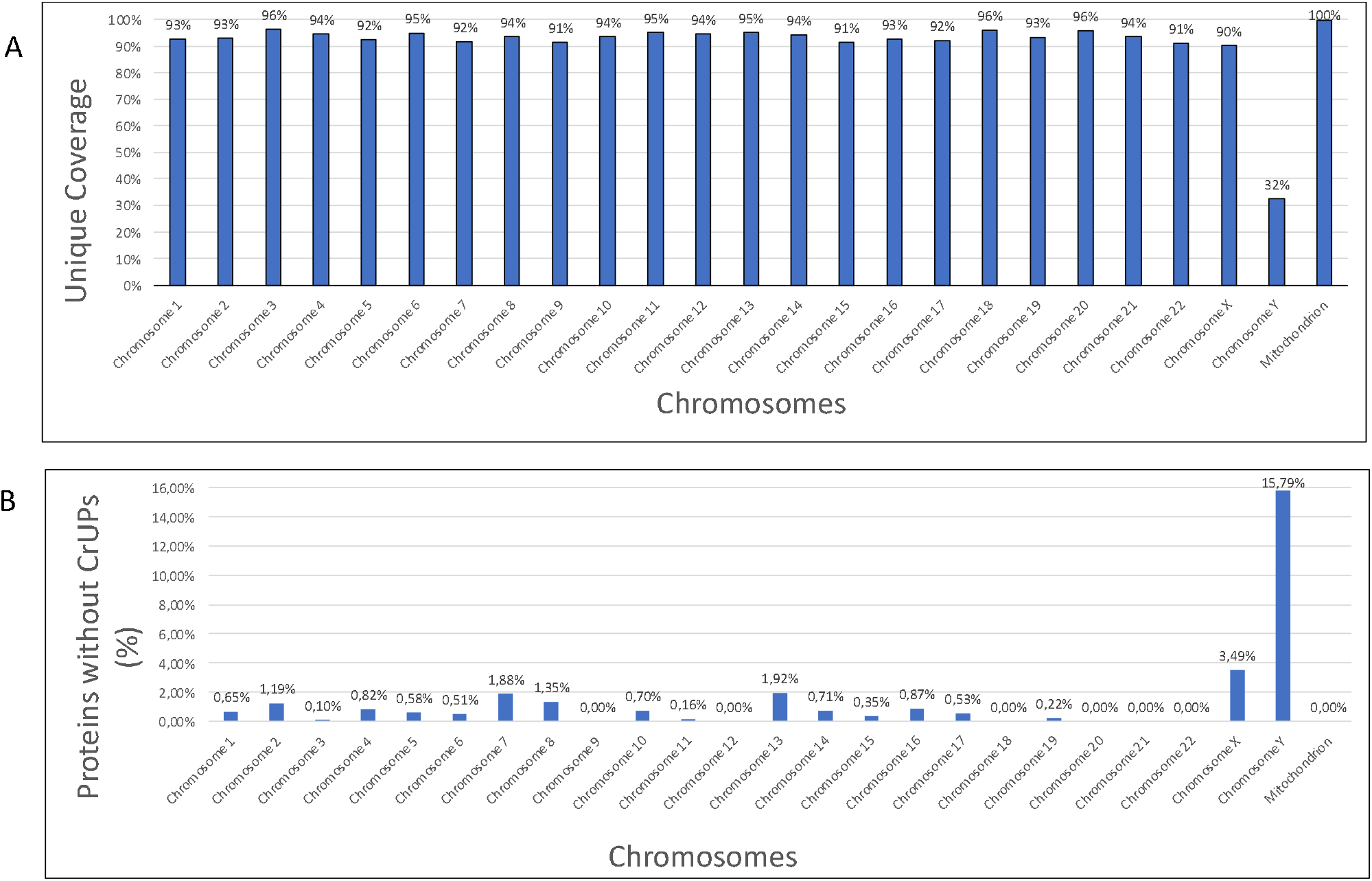
Chromosomal distribution of CrUPs. (**A**) Chromosomal coverage by CrUPs. The mitochondrial chromosome presents 100% coverage by CrUPs, while the Y chromosome appears the lowest coverage. (**B**) Chromosomal distribution of proteins that lack CrUPs. Y sex-chromosome accommodates the highest percentage of CrUP-missing proteins, whereas the proteins of mitochondrial chromosome are all presented to contain CrUPs.

### Clustering of CrUPs to Protein Families

Next, we examined the classification of CrUPs in protein families. 2,410 protein families, with more than 2 members, are being included in the SwissProt (Release 5_23) and InterPro (Release 97.0) protein databases (Table S6). To reliably evaluate the obtained results, we calculated the density and the coverage of each protein family members by their cognate CrUP species. We found that 3 protein families present the highest density (77%) and 801 protein families are characterized by 100% coverage. It seems that protein families with few members exhibit high density and coverage values. To better assess the significance of our results, we have selected, for further analysis, CrUP-containing families with more than 10 protein members each (292 families) (Table S7). The family with the lowest density (3%), also carrying very low coverage (16%), is NPIP, which contains 19 members. NPIP, together with GOLGA8, NBPF and SPATA31 families, showing very low density values (4, 5 and 19%, respectively), belong to protein families being featured by segmentally duplicated regions in the genes shared among all copies of the respective gene family (17), resulting in the same aa sequence that is being included multiple times in all family members, thus notably reducing the number of identified CrUPs. Multiple alignment of the 19 members all belonging to the NPIP protein family provides a convincing example and confirms the aforementioned finding, also introducing a new UP entity: the “**Family Unique Peptide**” (“FUP”). FUPs define peptides that are common between all members of a given family, but unique only for the protein members of the particular family (Figure S6), thereby indicating that the identification of a FUP species in a protein is capable to reliably classify this protein as member of the family. Interestingly, multiple alignment of the NPIP family members discloses the identification of 6 FUP species, residing at the aa positions 102-107, 169-179, 191-198, 239-244, 246-154 and 258-266, respectively (Figure S6).

FUP species can likely serve as a powerful tool to classify novel proteins with unknown functions as new members of a given family with defined activities. Hence, we, next, searched for proteins containing NPIP-specific FUPs in the unreviewed human proteins of Uniprot and Interpro datasets. Remarkably, we identified 113 unreviewed proteins containing NPIP-specific FUPs, with 91 being related to NPIP family and 22 lacking known functions, presumably derived from, hitherto, uncharacterized and novel genes, strongly suggesting the expansion of NPIP family with 22 additional protein members (Table 2A). Given the critical contribution of nuclear pore complex abnormalities to neurodegenerative diseases (18), the 22, putative new NPIP (Nuclear Pore Complex-Interacting Protein) family members, may prove to play essential roles in Alzheimer’s and Huntington pathologies (17, 19).

**Table 2.**
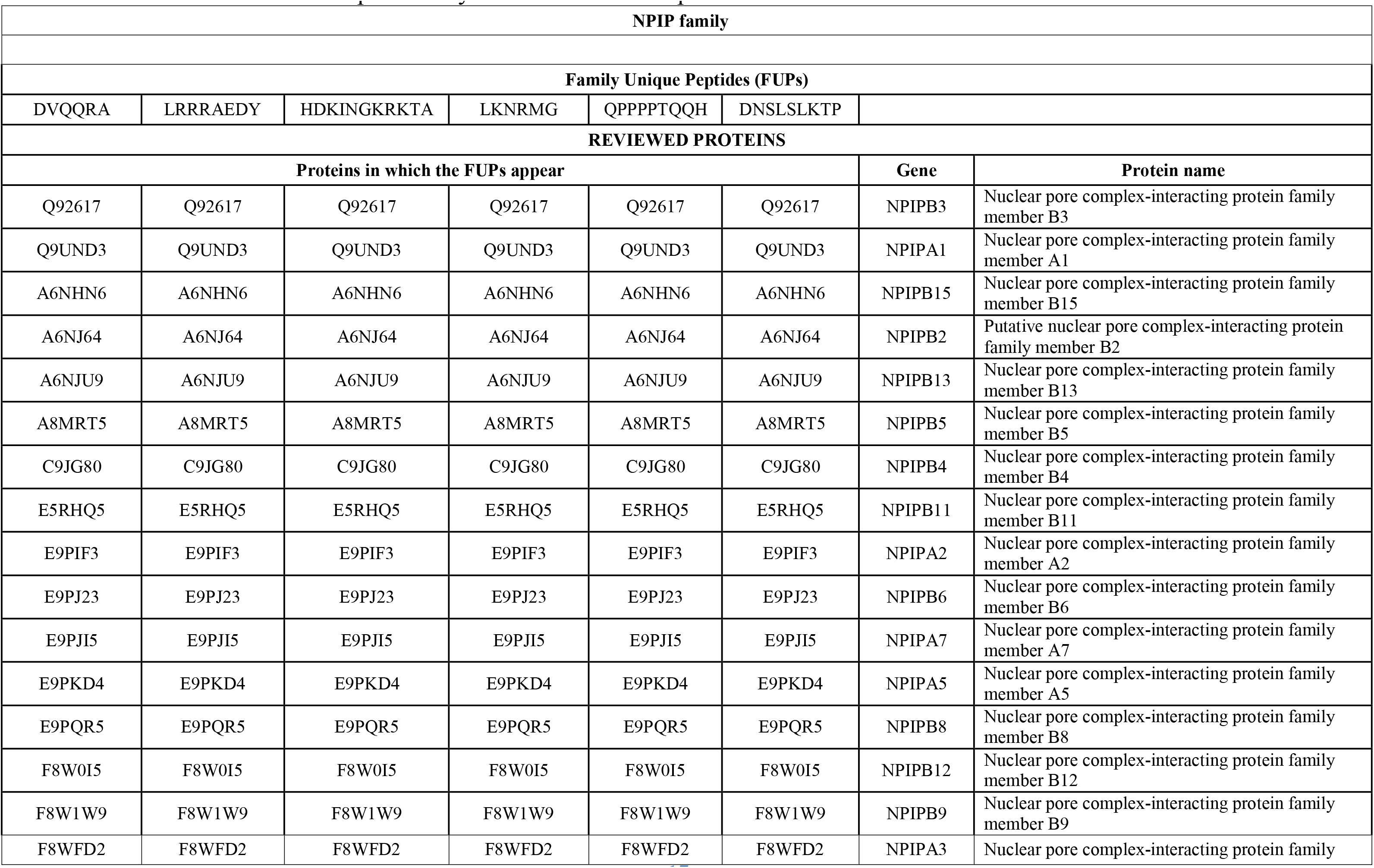
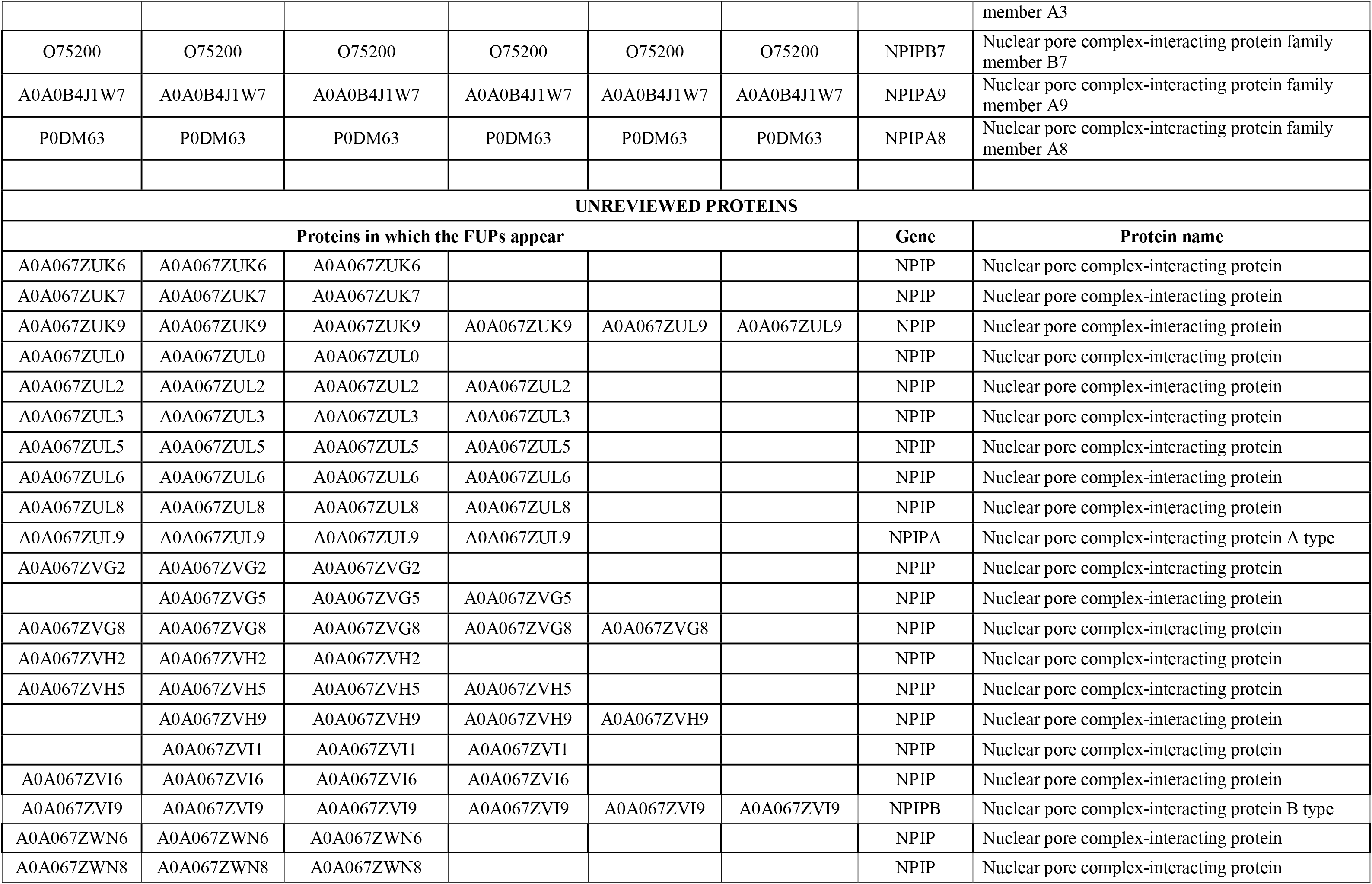
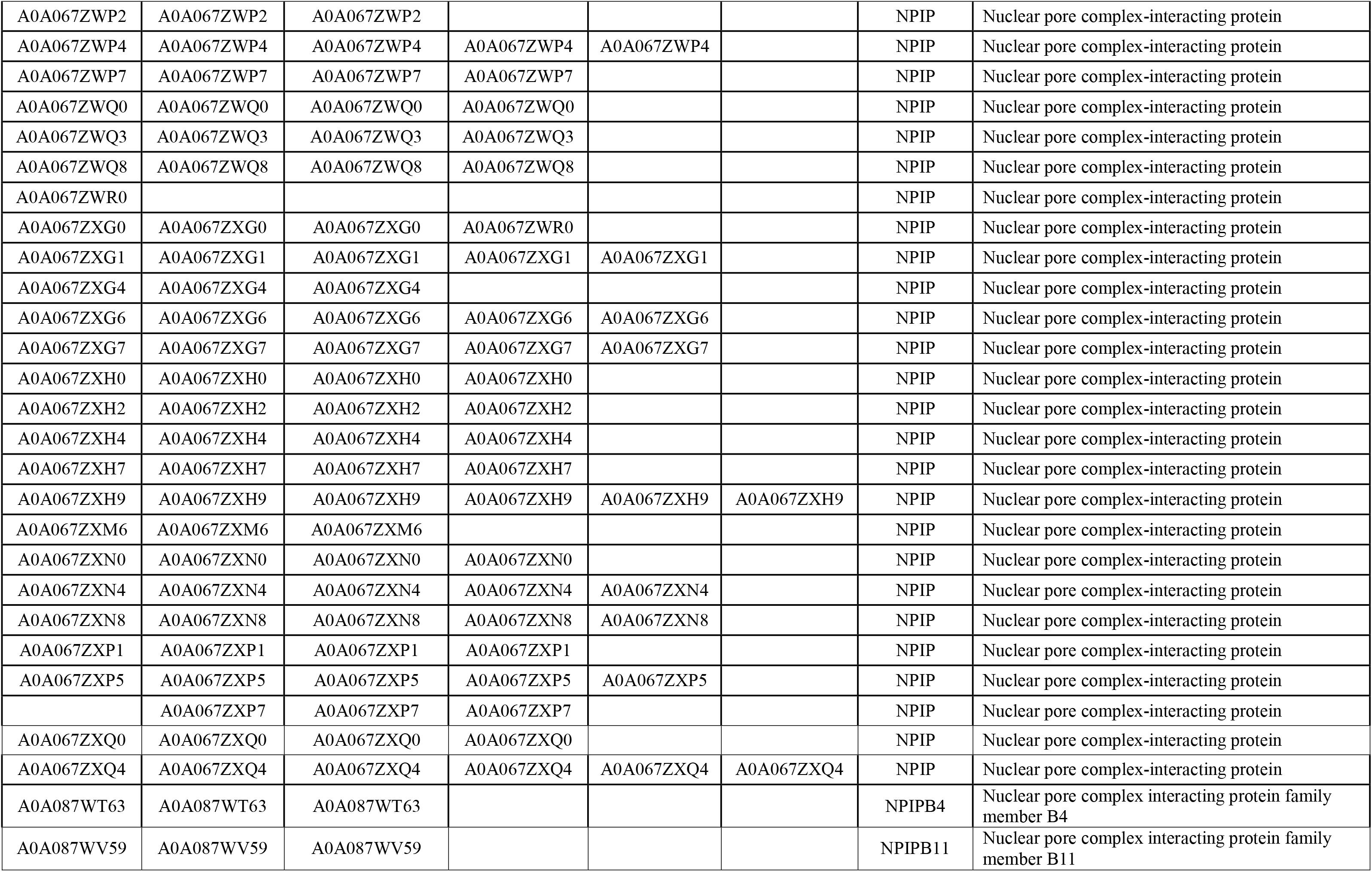
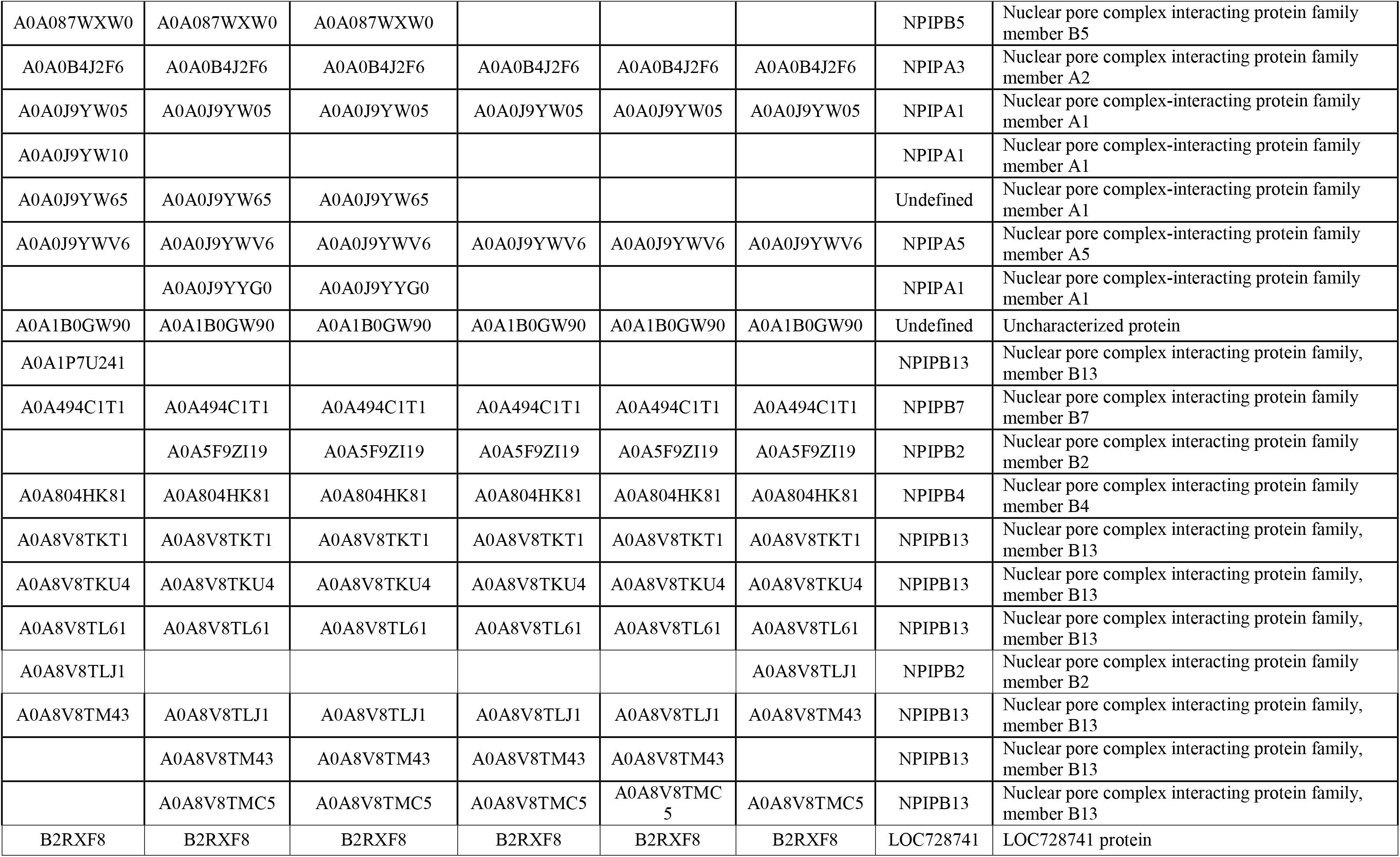
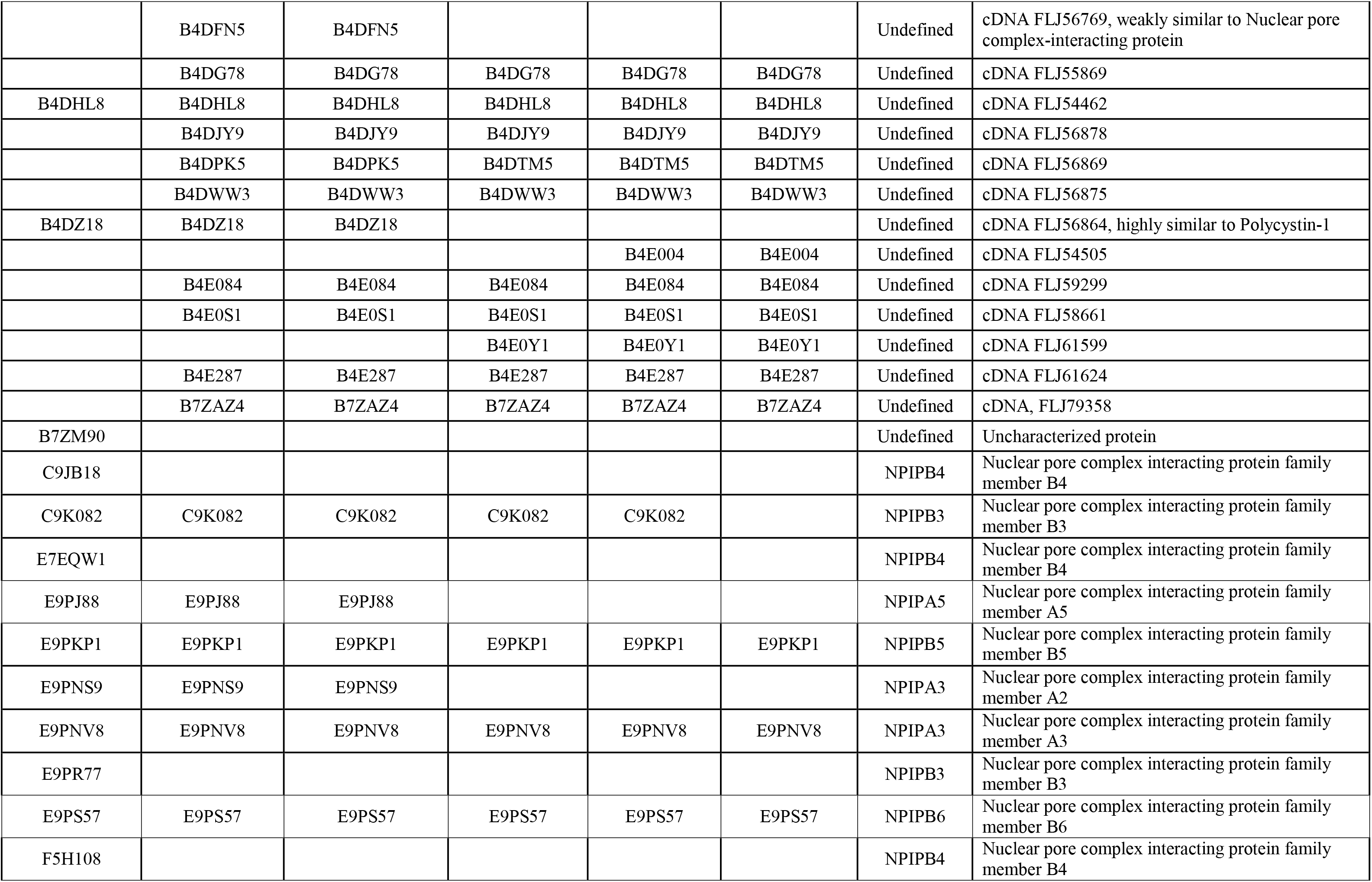
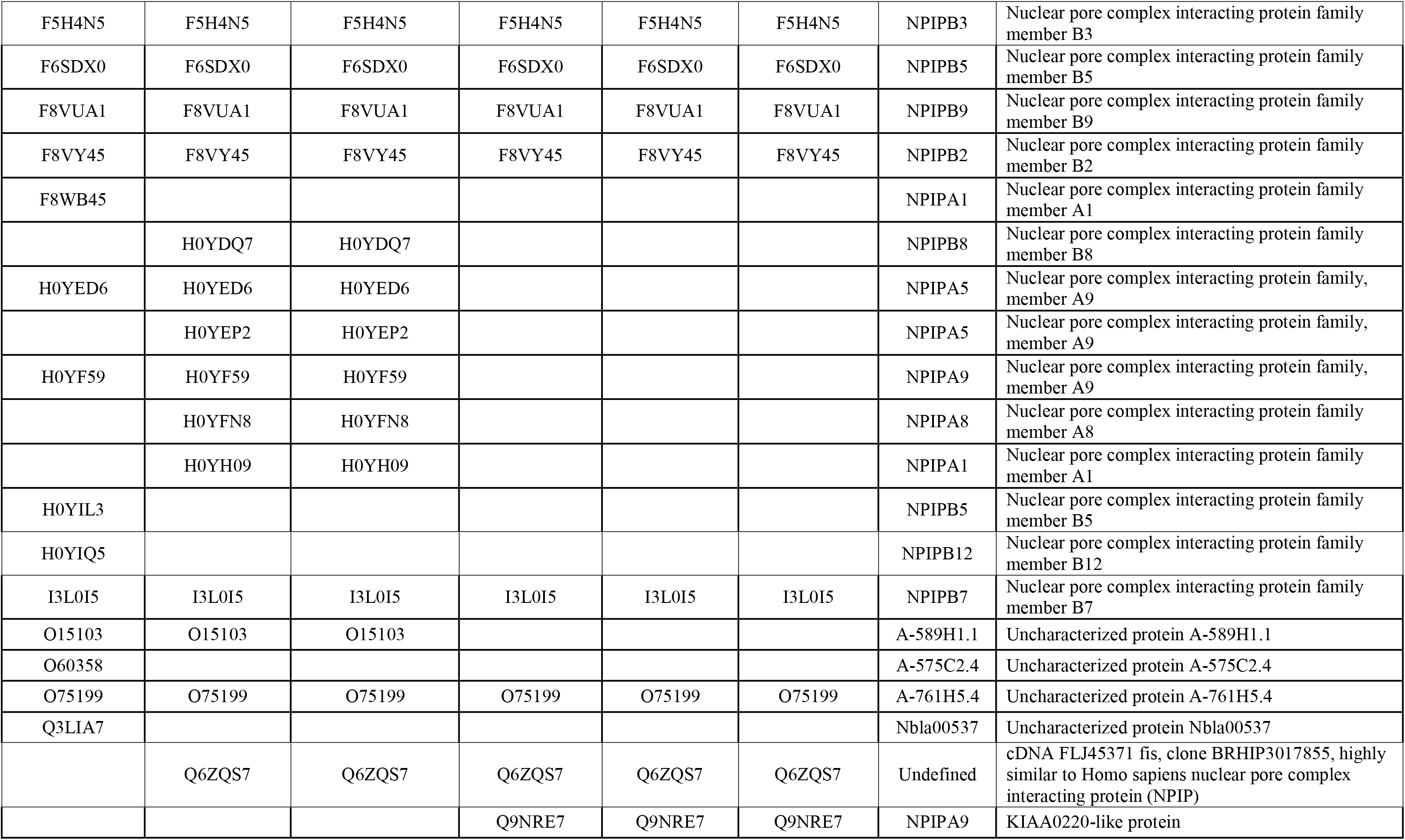
A. Identification of NPIP proteins by the FUPs in human proteome.

**Table 2B.** Probing of FUPs in the RAS oncoprotein family.

| RAS oncoprotein family |  |  |
| --- | --- | --- |
| Family Unique peptide: EEYSAM |  |  |
| REVIEWIED PROTEINS |  |  |
| Proteins in which the FUP appear | GENE | PROTEIN NAME |
| P01111 | NRAS | GTPase Nras |
| P01112 | HRAS | GTPase Hras |
| P01116 | KRAS | GTPase Kras |
| UNREVIEWED PROTEINS |  |  |
| Proteins in which the FUP appear | GENE | PROTEIN |
| A0A024RAV5 | KRAS | V-Ki-ras2 Kirsten rat sarcoma viral oncogene homolog, isoform CRA_b |
| X5D945 | HRAS | Harvey rat sarcoma viral oncoprotein isoform A |
| I1SRC5 |  | UBE2L3/KRAS fusion protein |
| A0A221ZRZ4 | KRAS | K-Ras |
| A0A3G1LBH1 | KRAS | Proto-oncogene GTPase |
| A0A804HJ06 | HRAS | HRas proto-oncogene, GTPase |
| A0A804HKM6 | HRAS | HRas proto-oncogene, GTPase |
| A0A8I5KQ21 | KRAS | KRAS proto-oncogene, GTPase |
| L7RSL8 | KRAS | V-Ki-ras2 Kirsten rat sarcoma viral oncogene homolog |
| Q14014 |  | PR310 c-K-ras protein |
| Q14015 |  | PR371 c-K-ras oncogene |
| Q5U091 | NRAS | Neuroblastoma RAS viral (V-ras) oncogene homolog |
| Q92468 | c-bas/has | p21 protein |
| Q9UM97 | N-ras | N-ras protein |

**Table 2C.** Tryptic digest unique peptides (TUPs) in the human proteome.

|  | Number | Tryptic Digest Peptides | Tryptic digest peptides containing CrUPs | Proteins with TUPs | Proteins without TUPs |
| --- | --- | --- | --- | --- | --- |
| Reviewed proteins | 20,422 | 818,082 |  |  |  |
| Reviewed proteins Include CrUPs | 20,258 | 815,026 | 564,148 (69.22%) | 20,136 (99.32%) | 122 (0.7%) |

Another characteristic example is the RAS family, which embraces the RASK, RASN and RASH proteins. They all share the same length of 189 aa, and despite their extremely high homology, 41, 45 and 41 CrUP species can be respectively detected (Table 2B). Their multiple alignment profile shows that the 1-80 aa sequences are identical and the CrUPs are being recognized within the 81-189 protein sequences (Figure S7). The “EEYSAM” peptide at 64-69 aa position is shared among all 3 RAS family members (K, N and H), and is also unique only to the 3 RAS proteins. Since it also bears the lowest aa length, “EEYSAM” can be considered as a RAS-specific core FUP (“cFUP”) entity (Figure S7). When we searched for proteins containing the “EEYSAM” cFUP species in unreviewed protein datasets (Uniprot and Interpro), 14 proteins were unveiled to accommodate the RAS-specific cFUP, thus indicating their classification, for the first time, in the RAS protein family. Given the major importance of RAS mutations in human oncogenesis (20), the 14, putative new RAS family members, may critically control tumor initiation and progression, *in vivo*, thereby opening new therapeutic windows for the disease in the clinic. Taken together, FUPs seem to emerge as new, powerful and important species for the illumination of proteome component functions, since their identification within the aa sequence of a given protein enables its classification into a particular family, thus unraveling the mystery of unknown protein activities. Of note, FUP identification will also facilitate the design of novel antigenic -peptide-sequences and the production of new antibodies specifically recognizing all protein members belonging to a family of interest.

Finally, we found that 20 families are presented with complete coverage (100%) by their respective CrUP species, because their protein members do not share identical regions in their sequences (Table S7 and Figure S8). Protein families with special interest to human biology and pathology, such TNF, IL-1, Helicase and BCL-2, feature high density and complete coverage by their respective CrUP species (100%), thus underlying the eminently significant and unique biological roles of these proteins in cellular patho-physiology. Likewise, protein families related to transmembrane receptors, such as the IL-1 receptor, and the type I or type II cytokine receptors, exhibit complete coverage (100%) values by CrUP species, thereby corroborating the high specificity and strong affinity of each receptor binding activity to its cognate ligand.

### Tryptic Digest UPs - TUPS

The detection of CrUPs in human proteome dictates our emerging ability to reliably identify a protein by use of only one peptide. This premise is crucial for proteomics, since protein identification by MS is the most popular and widely accepted technology, thus far. Briefly, a given protein mixture is digested by a proteolytic enzyme, usually trypsin, with the produced tryptic peptides being identified by MS platforms, ultimately leading to the desired recognition of proteins in the examined material (21). Towards this direction, all the reviewed proteins of human proteome containing CrUPs (20,258 proteins) were *in silico* digested by trypsin and the trypsin digest-generating peptides were analyzed for the presence of CrUPs, herein defined as “**Tryptic digest Unique Peptides**” (“TUPs”). 815,026 trypsin digest-derived peptides were produced from 20,258 proteins, with 564,148 of them containing CrUP species (69.22%) (Table 2C). The detected TUPs originated from 20,136 proteins (99.39%), whereas 122 proteins were found to lack TUP species. Therefore, it seems that the 20,136 reviewed human proteins could be reliably identified by use of only one TUP species. As shown in Table S8, the remaining (TUP-missing) 122 proteins mainly constitute isoforms with high aa sequence similarities among them, which likely necessitates the use of more than one tryptic peptide for their secure identification.

### Evaluation of CrUPs in other databases

Further searching in other databases, we founded that 89% of Cancer Antigenic Peptides (“CAPs”) and 87% of Immune Epitope Peptides (“IEPs”) contain at least one CrUP species, while it is demonstetaed that 97% of bioactives peptides included in the PeptideDB database contain at least one CrUP (Table S9) (22). The above strongly supporting the potentially beneficial applications of human CrUPs and Uniquome in tissue pathology, therapeutic oncology and translational medicine (23).

### Extension of the Human Uniquome to Other Organisms – an Evolutionary Perspective

Next, we evolutionary extended our study, by constructing the respective Uniquome maps of ten Vertebrata (vertebrates), six Magnoliopsida (flowering plants), one Nematoda (roundworm), one Arthropoda (arthropod), one Fungi (yeast) and one Bacteria (eubacteria). These organisms were selected based on their importance and use as model biological systems in a plethora of studies (Table 3). Intriguingly, we found that *Saccharomyces cerevisiae* (fungi) is presented with the highest percentage of reviewed proteins lacking CrUPs (2.99%), while all proteins that belong to zebrafish, rabbit and garden pea contain at least one CrUP species. The density of CrUPs is significantly lower in Magnoliopsida (flowering plants), in relation to the rest of the studied organisms (Table 3), indicating higher homologies among proteins of flowering plants, as compared to the respective ones in vertebrates. Further analysis of CrUP species per organism demonstrated that in organisms with the highest numbers of reviewed proteins the majority of CrUPs are 6-peptides, whereas in organisms with the lowest numbers of reviewed proteins the CrUP majority are 4-peptides (Figure 3 and Table S10). Altogether, it seems that as the number of reviewed proteins increase, the 6-peptide pattern will gain the majority of CrUP species map, strongly suggesting that in complete proteomes (in which all proteins will be reviewed) the majority of CrUPs will be the 6-peptide entity.

**Table 3.** Uniquomes of the major model organisms.

| Taxonomy | Organism | Reviewed Proteins (number) | Proteins with CrUPs (number) | Proteins without CrUPs (%) | CrUP (number) | Density of CrUP (%) |
| --- | --- | --- | --- | --- | --- | --- |
| Vertebrata (vertebrates) | Homo Sapiens (Human) | 20,422 | 20,258 | 164 (0.81%) | 7,271,607 | 64% |
|  | Mus musculus (Mouse) | 17,178 | 17,142 | 36 (0.21%) | 6,536,956 | 67% |
|  | Rattus norvegicus (Rat) | 8,182 | 8,171 | 11 (0.21%) | 2,825,568 | 67% |
|  | Bos taurus (Bovine) | 6,040 | 6,034 | 6 (0.10%) | 1,649,223 | 68% |
|  | Danio rerio (Zebrafish) | 3,306 | 3,306 | 0 | 1,093,006 | 68% |
|  | Gallus gallus (Chicken) | 2,307 | 2,305 | 2 (0.09%) | 766,884 | 68% |
|  | Sus scrofa (Pig) | 1,458 | 1,455 | 3 (0.21%) | 390,683 | 69% |
|  | Oryctolagus cuniculus (Rabbit) | 901 | 901 | 0 | 261,990 | 65% |
|  | Ovis aries (Sheep) | 466 | 463 | 3 (0.65%) | 93,208 | 65% |
|  | Equus caballus (Horse) | 292 | 291 | 1 (0.34%) | 64,920 | 66% |
| Magnoliopsida (Flower plants) | Arabidopsis thaliana (A. thaliana) | 16,351 | 16,326 | 115 (0.70%) | 4,295,898 | 58% |
|  | Zea mays (Maize) | 851 | 844 | 7 (0.83%) | 149,362 | 51% |
|  | Nicotiana tabacum (Common Tobacco) | 505 | 503 | 2 (0.40%) | 80,560 | 48% |
|  | Glycine max (Soybean) | 431 | 425 | 6 (1.41%) | 75,995 | 51% |
|  | Pisum sativum (Garden pea) | 402 | 402 | 0 | 76,518 | 59% |
|  | Triticum aestivum (Wheat) | 381 | 378 | 3 (0.79%) | 49,819 | 46% |
| Nematoda (roundworms) | Caenorhabditis elegans (C. Elegans) | 4,456 | 4,447 | 9 (0.20%) | 1,641,926 | 70% |
| Arthropoda (arthropods) | Drosophila melanogaster (fruit fly) | 3,723 | 3,694 | 29 (0.79%) | 1,623,848 | 70% |
| Fungi | Saccharomyces cerevisiae (Baker's yeast) | 6,727 | 6,532 | 195 (2.99%) | 2,002,523 | 66% |
| Bacteria (eubacteria) | Escherichia coli (E. Coli) | 4,357 | 4,287 | 70 (1.63%) | 952,021 | 70% |

**Figure 3.**
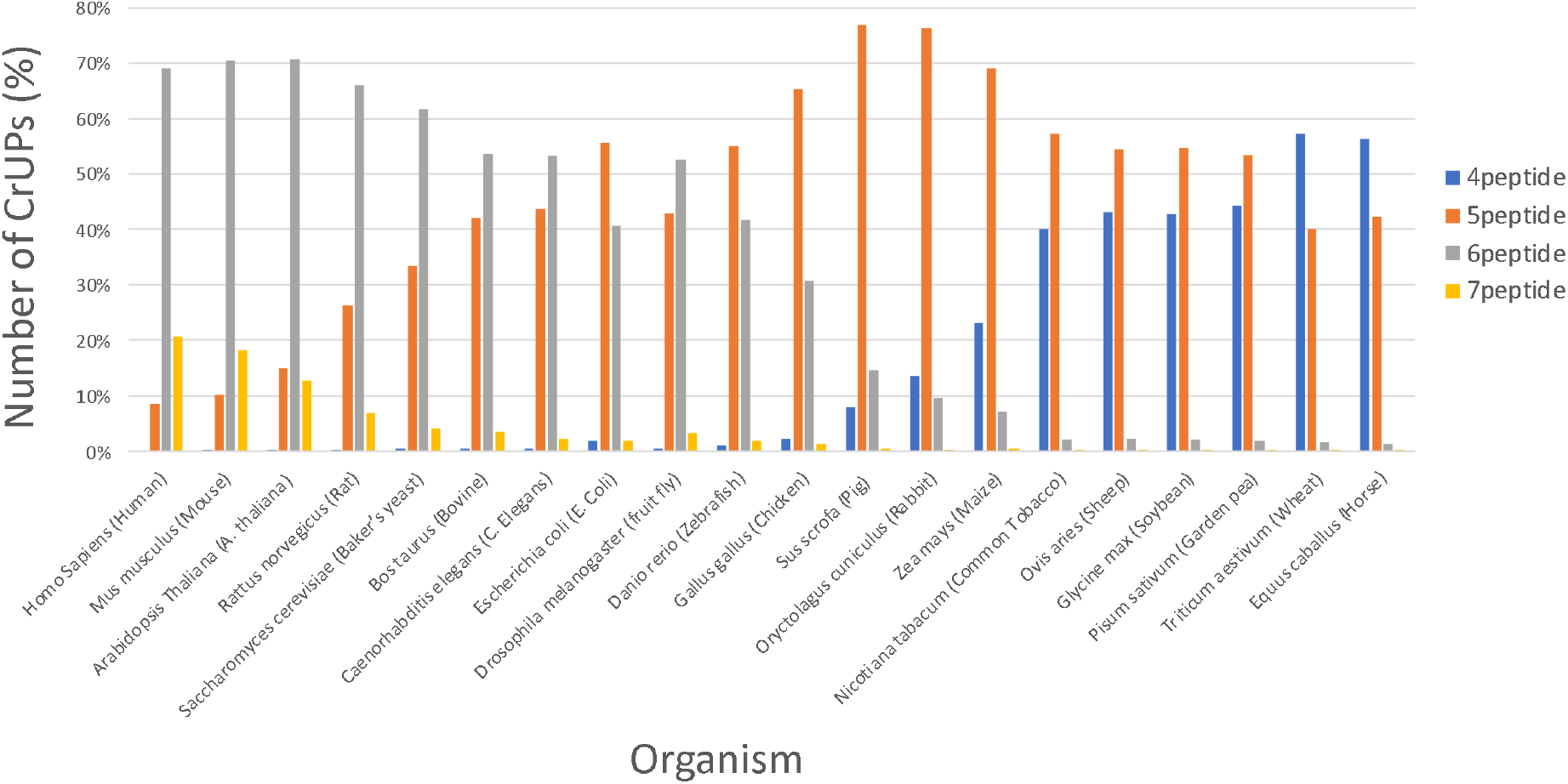
CrUP species profiling per organism. The percentage of 4-, 5-, 6-, and 7-aa length CrUPs, to the total number of CrUPs, of twenty model biological organisms are presented.

Our study, so far, has analyzed each protein of a given proteome in relation to organism’s other proteins, to investigate the presence of CrUPs. Subsequently, our study focused on detecting CrUPs of a protein across all organisms herein examined. Remarkably, we found that 20 reviewed proteins were commonly recognized in the 10 representative vertebrates included in this study and 37 proteins were identified in the six flowering plants, while between vertebrates and flowering plants there was no reviewed protein in common (Table S11). Next, for each one of the 20 common proteins among the 10 vertebrates, we identified the common CrUPs. These CrUPs, which exist in a given protein across organisms, were defined as “**Universal Unique Peptides**” (UUPs) and they have the important ability to securely identify a protein independently of an organism. As an example, the protein ATP6 (ATP synthase subunit a, MT-ATP6) was selected, because it is detected in all 10 vertebrates of our study. In Table 4, the common CrUPs of the ATP6 protein among the 10 vertebrates herein examined are presented. These CrUPs generate three UPs; the “FTPTTQLS”, the “VRLTAN” and the “IQAYVF”, which can be defined as presumable UUPs for the ATP6 protein in vertebrates. To validate our -novel-finding that these 3 CrUPs can serve as UUPs, specifically for the ATP6 protein in vertebrates, we, next, analyzed the scenario if these CrUPs could be detected exclusively and only in ATP6 of vertebrates. In Uniprot database, the ATP6 reviewed protein is referred in 68 vertebrate organisms. Hence, if these 3 CrUPs can act as UUPs, they must be identified exclusively in the ATP6 reviewed protein and only in these 68 vertebrates. To prove our argument, we recovered the totality of vertebrates’ reviewed proteins, from the Uniprot database (86,649 proteins, *in total*), and we searched the obtained dataset for proteins containing the 3 critical CrUPs. Strikingly, we revealed that among the 86,649 proteins examined, these novels, and herein described for the first time, 3 CrUPs were detected exclusively in the ATP6 protein of only the 68 vertebrates, as previously shown (100% success), thus indicating the capacity of these 3 CrUPs to operate as true UUPs for ATP6 in vertebrates, being able to safely and confidently identify only this protein (ATP6) among the plethora of all reviewed vertebrate proteins (Table S12).

**Table 4.**
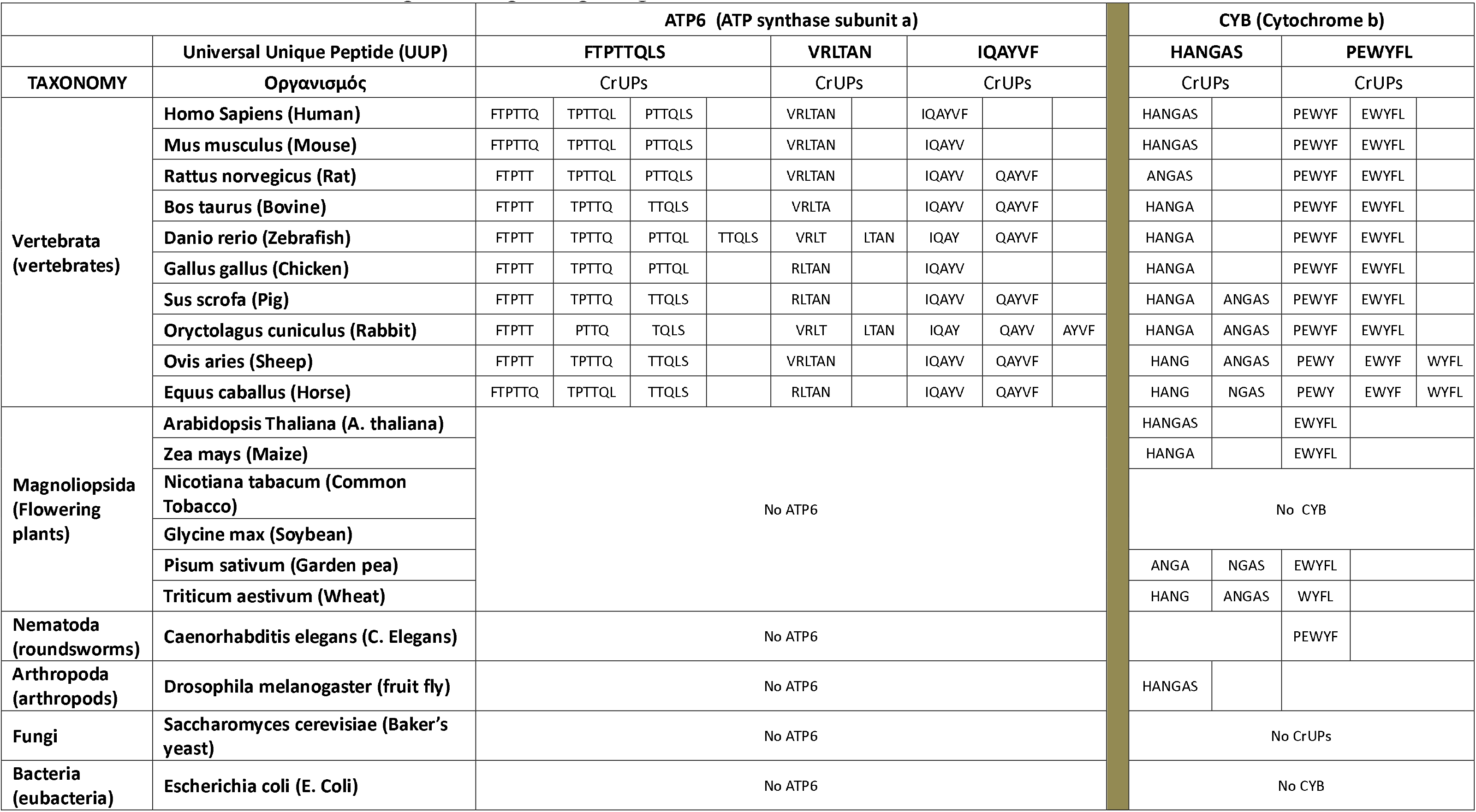
CrUPs in ATP6 and CYB proteins, participating in the formation of UUPs.

By the same strategy, we, additionally, analyzed the mitochondrial protein CYB (Cytochrome B), because it can be also recognized in a large number (1,764) of organisms. Interestingly, CYB is being presented as the protein that has been identified in the most numerous sets of organisms in the Uniprot database. As it is presented in Table 4, the reviewed CYB protein is detected in 17 out of the 20 organisms included in our study. The common CrUPs of the CYB protein among these 17 organisms dictated the ability of “HANGAS” and “PEWYFL” peptides to act as UUPs. Next, we tested the scenario if these 2 CrUPs could be identified in the CYB protein across eukaryotes. In the Uniprot database, 8,286 eukaryotes are being included, and 197,016 eukaryotic proteins have been reviewed (Table S13). In this voluminous protein dataset, the “HANGAS” 6-peptide was identified in 1,597 proteins, with 1,591 of them being characterized as CYB (99.62% success). Similarly, the “PEWYFL” 6-peptide was detected in 1,626 proteins, with 1,625 of them being classified as CYB (99.99% success). Most importantly, both 6-peptides were recognized in 1,512 reviewed proteins, with all of them being featured as CYB (100% success) (Table S13). Taken together, it seems that these 2 critical 6-peptides serve as UUPs for the CYB protein in eukaryotes. As shown in Table 4, in 2 flowering plants, the *Nicotiana tabacum* (Common Tobacco) and the *Glycine max* (Soybean), the CYB protein has not been detected into their, respective, reviewed proteomes. Therefore, we used the 2 -aforementioned-UUPs for CYB, to identify CYB, or CYB-like, proteins in the unreviewed proteomes of these 2 organisms. The results, presented in Table S14, show that 3 and 2 CYB, or CYB-like, proteins were identified in the unreviewed proteomes of Tobacco and Soybean plants, respectively. Following stringent criteria and procedures imposed by Uniprot, these newly recognized (CYB) proteins can be likely included into the, respective, -forthcoming-reviewed proteomes of these flowering plants.

Our -novel and exciting-finding that UPs / CrUPs can identify a protein of interest regardless of its cognate proteome strongly suggests that UUPs are conserved aa sequences within a protein, across species. Thereby, the final step of our study comprises the identification of DNA nucleotide sequences that, when translated, lead to UUPs production. To explore this, and with the “redundancy” limitation of Genetic Code (GC), the same UUP could be translated by either an identical or similar DNA nucleotide sequence, across species. To exemplify this issue, we analyzed the DNA nucleotide sequences of “PEWYFL” UUP for CYB protein across the *CYB* gene of the 14 organisms being characterized by the CYB protein detection. We found that in vertebrates there is a notable variation in the DNA sequence that codes for the “PEWYFL” UUP, with only human and mouse exhibiting an 100% homology identity in their respective sequences (“CCC GAA TGA TAT TTC CTA”). Likewise, all 4 flowering plants were presented with an identical among them “PEWYFL” UUP-coding DNA sequence (“CCG GAA TGG TAT TTC CTA”), which, however, significantly differ from the human/mouse respective one (Table 5). Strikingly, the respective DNA nucleotide sequences of the 9 vertebrates (including human), herein examined, proved to contain the triplet - codon “TGA”. In contrast to GC that imposes on the “TGA” codon to act as “stop - termination signal” (“stop codon”) for translation, our results clearly indicate that in the vertebrate CYB-specific context, it (“TGA”) can be translated into the aa Tryptophan (W). Such an exceptional phenomenon was previously observed in some prokaryotes [24] and eukaryotic (e.g., fungi) mitochondrial genomes and their translational machineries [25]. However, to our knowledge, it is the first time that “TGA” serves as a W-specific codon in higher vertebrates and mammals, including human, which is very important in order to understand the translation termination machinery, because a number of human heritable diseases related to stop codons (26).

**Table 5.** Conversion of the 6-aa peptide “PEWYFL” to DNA 18-nucleotides, according to the genetic code.

| TAXONOMY | Organism | Nucleotide |
| --- | --- | --- |
| Vertebrata<br>(vertebrates) | Homo Sapiens (Human) | ccc gaa tga tat ttc cta |
|  | Mus musculus (Mouse) | ccc gaa tga tat ttc cta |
|  | Rattus norvegicus (Rat) | cca gaa tga tat ttt ctc |
|  | Bos taurus (Bovine) | ccc gag tga tac ttc tta |
|  | Danio rerio (Zebrafish) | cca gaa tga tat ttc cta |
|  | Gallus gallus (Chicken) | cca gaa tga tat ttt cta |
|  | Sus scrofa (Pig) | cca gaa tga tat ttc tta |
|  | Oryctolagus cuniculus (Rabbit) | cca gaa tga tac ttt cta |
|  | Ovis aries (Sheep) | cct gaa tga tac ttc cta |
|  | Equus caballus (Horse) | cca gaa tgg tac ttc ctg |
| Magnoliopsida<br>(Flower plants) | Arabidopsis Thaliana (A. thaliana) | ccg gaa tgg tat ttc cta |
|  | Zea mays (Maize) | ccg gaa tgg tat ttc cta |
|  | Pisum sativum (Garden pea) | ccg gaa tgg tat ttc cta |
|  | Triticum aestivum (Wheat) | ccg gaa tgg tat ttc cta |

DNA-nucleotide sequence, coding for the “PEWYFL” UUP, across species. Colors indicate the codon differences in various organisms, as compared to the reference human codons. For Flower plants, the DNA-nucleotide sequence generated from the UUP peptide is identical among all family members, whereas, in vertebrates, the different DNA sequences translating the same UUP species are shown by a color code, with the TGA codon being indicated in red.

Altogether, it seems that across species the conserved sequencies are not DNA nucleotides but peptides, with mother Nature promoting and supporting the -evolutionary- conservation of aa sequences in peptides and proteins, thus creating molecular identity profiles that result in phenotypic signatures of organisms. Thus, a multiparametric cluster analysis of UUPs aa development of given proteins across species, could restructure the evolution and the taxonomy as it is possible to reveal novel phenotypically closer species.

## Author contributions

G.T.T.: conceptualization and supervision; G.T.T, and D.J.S.: scientifically responsible; C.E.V., I.P. and D.J.S.: thesis supervisory committee of E.K.; G.T.T., E.K. and V.P.: methodology development; E.K. and V.P.: software and database development, with guidance from G.T.T.; G.T.T. and D.J.S.: writing the original draft; E.K. and G.T.T.: preparation of tables and figures; G.T.T. and D.J.S.: reviewing and editing the final draft. All authors read and approved the final manuscript.

## Data and materials availability

All data are available in the main text or the supplementary materials. The results of the present study are also available in the website “Uniquome” (www.uniquome.com).

## Supplementary Materials

Materials and Methods

Figures S1 to S8

Tables S1 to S14

## References

1. J. E. Hixson, D. A. Clayton, (1985). Initiation of transcription from each of the two human mitochondrial promoters requires unique nucleotides at the transcriptional start sites. Proceedings of the National Academy of Sciences 82(9), 2660–2664 (1985). doi: 10.1073/pnas.82.9.2660

2. H. G. Almanghadim, S. Ghorbian, N.S. Khademi, M.S. Sadrabadi, E. Jarrahi, Z. Nourollahzadeh, M. Dastani, M. Shirvaliloo, R. Sheervalilou, S. Sargazi, New insights into the importance of long non-coding RNAs in lung cancer: Future clinical cpproaches. DNA and Cell Biology 40(12), 1476–1494 (2021). doi: 10.1089/dna.2021.0563.

3. D.R. Bobbili, P. Banda, R. Krüger, P. May, Excess of singleton loss-of-function variants in Parkinson’s disease contributes to genetic risk. Journal of Medical Genetics 57(9), 617–623 (2020). doi: 10.1136/jmedgenet-2019-106316.

4. Mouratidis, C.S.Y. Chan, N. Chantzi, G.C. Tsiatsianis, M. Hemberg, N. Ahituv, I. Georgakopoulos-Soares, Quasi-prime peptides: identification of the shortest peptide sequences unique to a species, NAR Genomics and Bioinformatics 5(2), lqad039 (2023). doi: 10.1093/nargab/lqad039

5. Alexandridou, G.T. Tsangaris, K. Vougas, K. Nikita, G. Spyrou, UniMaP: finding unique mass and peptide signatures in the human proteome. Bioinformatics 25(22), 3035–3037 (2009). doi: 10.1093/bioinformatics/btp516.

6. P. Sinitcyn, A.L. Richards, R.J. Weatheritt, D.R. Brademan, H. Marx, E. Shishkova, J.G. Meyer, A.S. Hebert, M.S. Westphall, B.J. Blencowe, J. Cox, J.J. Coon, Global detection of human variants and isoforms by deep proteome sequencing. Nature Biotechnology 41(12), 1776–1786 (2023). doi: 10.1038/s41587-023-01714-x.

7. Alexandridou, G.T. Tsangaris, K. Vougas, K. Nikita, G. Spyrou, Peptide Finder: mapping measured molecular masses to peptides and proteins, Bioinformatics 24(19), 2267–2269 (2008). doi:10.1093/bioinformatics/btn413

8. Alexandridou, N. Dovrolis, G.T. Tsangaris, K. Nikita, G. Spyrou, PepServe: a web server for peptide analysis, clustering and visualization. Nucleic Acids Research 39(Web Server issue), W381–384 (2011). doi: 10.1093/nar/gkr318.

9. E. Kontopodis, V. Pierros, D.J. Stravopodis, G.T. Tsangaris, Prediction of SARS-CoV-2 Omicron variant immunogenicity, immune escape and pathogenicity, through the analysis of spike protein-specific core unique peptides. Vaccines (Basel) 10(3), 357 (2022). doi: 10.3390/vaccines10030357.

10. V. Pierros, E. Kontopodis, D.J. Stravopodis, G.T. Tsangaris, Unique peptide signatures of SARS-CΟV-2 virus against human proteome reveal variants’ immune escape and infectiveness. Heliyon 8(4), e09222 (2022). doi: 10.1016/j.heliyon.2022.e09222.

11. U. C. Lavania, Chromosome diversity in population: Defining conservation units and their micro-identification through genomic in situ painting. Current Science 83(2), 124–127 (2002). http://www.jstor.org/stable/24106214.

12. T. Lappalainen, M. Sammeth, M. Friedländer et al. Transcriptome and genome sequencing uncovers functional variation in humans. Nature, 501, 506–511 (2013). doi:10.1038/nature12531

13. Rhie, S. Nurk, M. Cechova et al, The complete sequence of a human Y chromosome. Nature 621(7978), 344–354 (2023) doi: 10.1038/s41586-023-06457-y.

14. P. Hallast, P. Ebert, M. Loftus et al, Assembly of 43 human Y chromosomes reveals extensive complexity and variation. Nature 621, 355–364 (2023). doi:10.1038/s41586-023-06425-6.

15. J. Cummins, Mitochondrial DNA and the Y chromosome: parallels and paradoxes. Reproduction Fertility and Development 13(7-8), 533–542 (2001). doi: 10.1071/rd01064.

16. D. Wallace, Mitochondrial DNA in evolution and disease. Nature 535, 498–500 (2016). 10.1038/nature18902.

17. Bekpen, D. Tautz, Human core duplicon gene families: game changers or game players? Briefings in Functional Genomics 18(6), 402–411 (2019) doi: 10.1093/bfgp/elz016.

18. A.N. Coyne, J.D. Rothstein, Nuclear pore complexes - a doorway to neural injury in neurodegeneration. Nature Reviews. Neurology 18(6), 348–362 (2022). doi: 10.1038/s41582-022-00653-6.

19. N.W. Van Bibber, C. Haerle, R. Khalife, G.W. Dayhoff, V.N. Uversky, Intrinsic disorder in human oroteins encoded by core duplicon gene families. Journal of Physical Chemistry B 124(37), 8050–8070 (2020) doi: 10.1021/acs.jpcb.0c07676.

20. S. Li, A. Balmain, C.M. Counter, A model for RAS mutation patterns in cancers: finding the sweet spot. Nature Reviews. Cancer 18(12), 767–777 (2018) doi: 10.1038/s41568-018-0076-6.

21. A.K. Anagnostopoulos, D.J. Stravopodis, G.T. Tsangaris, Yield of 6,000 proteins by 1D nLC-MS/MS without pre-fractionation. Journal of Chromatography B. Analytical Technologies in Biomedical and Life Sciences 1047, 92–96 (2017). doi: 10.1016/j.jchromb.2016.08.031.

22. T. Chamoli, A. Khera, A. Sharma, A. Gupta, S. Garg, K. Mamgain, A. Bansal, S. Verma, A. Gupta, H.K. Alajangi, G. Singh, R.P. Barnwal, Peptide Utility (PU) search server: A new tool for peptide sequence search from multiple databases. Heliyon 8(12), e12283 (2022). doi: 10.1016/j.heliyon.2022.e12283.

23. R. Chen, K.M. Fulton, S.M. Twine, J. Li, Identification of MHC peptides using mass spectrometry for neoantigen discovery and cancer vaccine development. Mass Spectrometry Reviews 40(2), 110–125 (2021) doi: 10.1002/mas.21616.

24. T.Y. Wong, S. Fernandes, N. Sankhon, P.P. Leong, J. Kuo, J.K. Liu, Role of premature stop codons in bacterial evolution. Journal of Bacteriology 190(20), 6718–6725 (2008) doi: 10.1128/JB.00682-08.

25. P.R. Copeland, Regulation of gene expression by stop codon recoding: selenocysteine. Gene 312, 17–25 (2003). doi: 10.1016/s0378-1119(03)00588-2.

26. M.R. Lawson, L.N. Lessen, J. Wang, A. Prabhakar, N.C. Corsepius, R. Green, J.D. Puglisi, Mechanisms that ensure speed and fidelity in eukaryotic translation termination. Science 373(6557), 876–882 (2021) doi: 10.1126/science.abi7801.

