## SUPPLEMENTARY MATERIAL for "Uniquome: Construction and Decoding of a Novel Proteomic Atlas that Contains New Peptide Entities"

### MATERIALS AND METHODS

#### 1. Materials

##### 1.1. Uniquome

We developed a toolset named "**Uniquome Tools Suite**" based on big-data analysis approaches, in order to create the necessary datasets for this study. The tool was developed using .NET Core 6.0 and C# and the results are stored in a PostgreSQL database. The tool can generate the following:

- Save proteomes and their metadata (i.e., protein length, amino acid composition, protein families etc).
- Create the Uniquome of a given proteome.
- Extract Core Unique peptides (CrUPs) from a Uniquome
- Composite Unique Peptides (CmUPs) from a Uniquome.
- Extract Tryptic-digested Unique Peptides (TUPs) from a Uniquome.

A website entitled "Uniquome" (www.uniquome.com) was created, for the results of the present study. The user can search by CrUPs, CmUPs and TUPs on an organism.

##### 1.2. Databases

The following databases were used:

• Uniprot (www.uniprot.org) [Release 05_2023]
• Interpro (https://www.ebi.ac.uk/interpro/) [Release 97.0, 9 Nov. 23)
• Imune Epitope Database (www.iedb.org)
• Cancer Antigenic Peptide Database (https://caped.icp.ucl.ac.be/)
• Bioactive Peptide Database (http://www.peptides.be/)
• National Center for Biotechnology Information (https://www.ncbi.nlm.nih.gov)
• Kyoto Encyclopedia of Genes and Genomes (https://www.genome.jp/kegg/)

#### 2. Methods

##### 2.1.1. Core Unique Peptides (CrUPs)

The identification of CrUPs in a given proteome was performed by the custom-made tool based on big data analysis approaches (9, 10). The tool is searching for the CrUPs of a given proteome by splitting each protein of the proteome in peptides with lengths from 4 to 100 amino acids using a rolling window from the first amino acid. For a given protein with length L and window W, a total of "**C = L-W+1**" peptides will be generated. All the produced peptides are stored for further analysis. To characterize a peptide as CrUP, the given peptide is searched against all the other proteins of the proteome and if it is found in at least one protein, it is, then, discarded as non-unique. If the peptide has been characterized as unique for the given protein, all the other peptides with longer amino acids length containing that peptide are also discarded as non-core. Finally, applying this method from the shorter amino acids to the larger ones, the peptides which are selected with the above criteria are considered as CrUPs. To handle the complexity of searching for a given peptide "s" occurrence across thousands of other proteins, we split the proteome in subsets of almost equally distributed amounts of proteins based on the available cores of the machine that will make the analysis. Each subset (called bucket) assumes sequentially the role of the main bucket.

Then, it splits each protein that is contained within the bucket to peptides of 4 to 100 amino acid length. These peptides that do not contain any CrUPs are, then, searched in parallel against all the other buckets (secondary buckets). The secondary buckets are equipped with a function to examine the existence of a peptide within their proteins. If a peptide is found in any of the secondary buckets, the search is stopped across all secondary buckets, the peptide is considered to lack CrUP property and the main bucket moves to search the next peptide. To speed execution, a pre-process stage is executed, where we create an amino acid residue mapping for each protein. When a peptide starts, for example, with Arginine (R), only substrings on the locations of the protein whereat R is found need to be searched, thus significantly reducing the search space, and speeding up execution.

**2.1.2. Composite Unique Peptides (CmUPs)**

CmUPs were constructed by two or more consecutive or overlapping CrUPs. The identification of CmUPs was performed by the "Uniquome Tools Suite" (UTS) based on the sequentially peptide placement on protein sequence. Thus, peptides constructed by at least two CrUPs (overlapping or consecutive) were considered as CmUPs.

**2.2 Metadata Analysis of Uniquome**

The metadata analysis of Unique Peptides (CrUPs and CmUPs), including length distribution, position on the protein sequence, protein coverage, density, chromosomal and protein family distribution across a given proteome, were performed by the "Uniquome Tools Suite" (UTS).

**2.3 Tryptic digest Unique Peptides (TUPs)**

Trypsin cleaves proteins C-terminal to Arginine (A) and Lysine (L) (if they are not followed by Proline {P}) residues and is, by far, the most used endoproteinase in bottom-up proteomics. To search for tryptic digest peptides containing CrUPs, we created an application that can cleave the Human reviewed proteins based on the aforementioned rule. Then, peptides derived from this process are being stored and they are, next, searched if they contain CrUP(s) within their amino acid sequences, to ultimately typify them as Tryptic digest Unique Peptides (TUPs).

**2.4 Evaluation of Unique Peptides (UPs) in other databases**

For the search of CrUPs in various peptide databases, we have designed applications specialized to each database capable of searching for CrUPs. By this application, a Cancer Antigenic Peptide database (), an Immune Epitope database () and a Bioactive Peptide database () were searched for CrUPs.

**2.5 Comparison of Unique Peptides (UPs) between organisms**

The "Uniquome Tools Suite" can compare Uniquomes between two or more organisms, to identify similarities between the UP collections of the examined organisms, allowing up to one amino acid residue mismatch.

**2.6 Family Unique Peptides (FUPs)**

Firstly, the proteins of each family under examination were selected from the reviewed Human proteins of Uniprot database. Secondly, the selected dataset of proteins was aligned using the special tool of Uniprot, and the common amino acid sequences among protein members of the family were marked. Next, in these sequences, using our "Uniquome Tool", the common peptides, which appeared in the proteins of a given family, were identified. To characterize a common peptide as FUP, it was checked if this peptide appeared and in other reviewed proteins besides the family of study. If the peptide appeared only in protein members of the family of study, then the common peptide is characterized as FUP.

**2.7 Universal Unique Peptides (UUPs)**

To characterize a CrUP as UUP of a given protein across species, the proteins that appeared most often in the study organisms were identified. Then, the common CrUPs in each dataset of the same protein, across the organisms of study, were recognized. For the characterization of each of the identified common CrUP as UUP, for the given protein across species, the reviewed proteome of all organisms included in Uniprot was retrieved, and by using a searching algorithm each one of the common CrUPs was searched in the above proteome dataset. The proteins, across species, that contained the CrUP were restored and the CrUP was characterized as UUP if it was identified only in the given protein across species. The "Identification Rate" for each UUP is the percentage (%) of the number of the given proteins across species that contain the UUP, to the total number of the given proteins across species.

**2.8 Core Unique Peptide (CrUP) and DNA nucleotide sequence correlation**

It was checked whether the DNA nucleotide sequence from which the cognate CrUP/UUP is produced, across species, is the same for all organisms containing that CrUP/UUP and protein. The NCBI and KEGG databases were used to identify the DNA nucleotide sequence(s) from which the UUP is translated. Each DNA nucleotide sequence was searched manually, by matching the peptide-derived DNA nucleotide sequence(s) to the above databases.

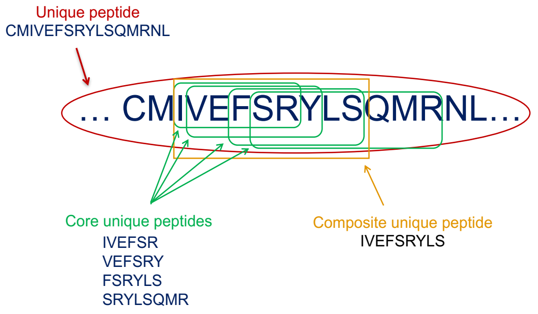

**Figure S1. Construction of CrUPs and CmUPs.**
To exemplify our new peptide entity-based rationale, we herein present a protein-derived "…CMIVEFSRYLSQMRNL…" Unique Peptide (UP), which, according to the definition of Core Unique Peptide (CrUP), contains the following CrUPs: "IVEFSR", "VEFSRY", "FSRYLS" and "SRYLSQMR". By the linear assembly of these CrUPs, the "IVEFSRYLS" Composite Unique Peptide (CmUP) is constructed (carries the "IVEFSR", "VEFSRY" and "FSRYLS" CrUPs).

A

```
sp|Q9UBX2|DUX4_HUMAN      MALPTPSDSTLPAEARGRGRRRRLVWTPSQSEALRACFERNPYPGIATRERLAQAIGIPE 60
sp|P0CJ86|DU4L3_HUMAN     MALPTPSDSTLPAEARGRGRRRRLVWTPSQSEALRACFERNPYPGIATRERLAQAIGIPE 60
sp|P0CJ85|DU4L2_HUMAN     MALPTPSDSTLPAEARGRGRRRRLVWTPSQSEALRACFERNPYPGIATRERLAQAIGIPE 60
sp|P0CJ89|DU4L6_HUMAN     MALPTPSDSTLPAEARGRGRRRRLVWTPSQSEALRACFERNPYPGIATRERLAQAIGIPE 60
sp|P0CJ90|DU4L7_HUMAN     MALPTPSDSTLPAEARGRGRRRRLVWTPSQSEALRACFERNPYPGIATRERLAQAIGIPE 60
sp|P0CJ88|DU4L5_HUMAN     MALPTPSDSTLPAEARGRGRRRRLVWTPSQSEALRACFERNPYPGIATRERLAQAIGIPE 60
                          ************************************************************

sp|Q9UBX2|DUX4_HUMAN      PRVQIWFQNERSRQLRQHRRESRPWPGRRGPPEGRRKRTAVTGSQTALLLRAFEKDRFPG 120
sp|P0CJ86|DU4L3_HUMAN     PRVQIWFQNERSRQLRQHRRESRPWPGRRGPPEGRRKRTAVTGSQTALLLRAFEKDRFPG 120
sp|P0CJ85|DU4L2_HUMAN     PRVQIWFQNERSRQLRQHRRESRPWPGRRGPPEGRRKRTAVTGSQTALLLRAFEKDRFPG 120
sp|P0CJ89|DU4L6_HUMAN     PRVQIWFQNERSRQLRQHRRESRPWPGRRGPPEGRRKRTAVTGSQTALLLRAFEKDRFPG 120
sp|P0CJ90|DU4L7_HUMAN     PRVQIWFQNERSRQLRQHRRESRPWPGRRGPPEGRRKRTAVTGSQTALLLRAFEKDRFPG 120
sp|P0CJ88|DU4L5_HUMAN     PRVQIWFQNERSRQLRQHRRESRPWPGRRGPPEGRRKRTAVTGSQTALLLRAFEKDRFPG 120
                          ************************************************************

sp|Q9UBX2|DUX4_HUMAN      IAAREELARETGLPESRIQIWFQNRRARHPGQGGRAPAQAGGLCSAAPGGGHPAPSWVAF 180
sp|P0CJ86|DU4L3_HUMAN     IAAREELARETGLPESRIQIWFQNRRARHPGQGGRAPAQAGGLCSAAPGGGHPAPSWVAF 180
sp|P0CJ85|DU4L2_HUMAN     IAAREELARETGLPESRIQIWFQNRRARHPGQGGRAPAQAGGLCSAAPGGGHPAPSWVAF 180
sp|P0CJ89|DU4L6_HUMAN     IAAREELARETGLPESRIQIWFQNRRARHPGQGGRAPAQAGGLCSAAPGGGHPAPSWVAF 180
sp|P0CJ90|DU4L7_HUMAN     IAAREELARETGLPESRIQIWFQNRRARHPGQGGRAPAQAGGLCSAAPGGGHPAPSWVAF 180
sp|P0CJ88|DU4L5_HUMAN     IAAREELARETGLPESRIQIWFQNRRARHPGQGGRAPAQAGGLCSAAPGGGHPAPSWVAF 180
                          ************************************************************

sp|Q9UBX2|DUX4_HUMAN      AHTGAWGTGLPAPHVPCAPGALPQGAFVSQAARAAPALQPSQAAPAEGISQPAPARGDFA 240
sp|P0CJ86|DU4L3_HUMAN     AHTGAWGTGLPAPHVPCAPGALPQGAFVSQAARAAPALQPSQAAPAEGVSQPAPARGDFA 240
sp|P0CJ85|DU4L2_HUMAN     AHTGAWGTGLPAPHVPCAPGALPQGAFVSQAARAAPALQPSQAAPAEGVSQPAPARGDFA 240
sp|P0CJ89|DU4L6_HUMAN     AHTGAWGTGLPAPHVPCAPGALPQGAFVSQAARAAPALQPSQAAPAEGVSQPAPARGDFA 240
sp|P0CJ90|DU4L7_HUMAN     AHTGAWGTGLPAPHVPCAPGALPQGAFVSQAARAAPALQPSQAAPAEGVSQPAPARGDFA 240
sp|P0CJ88|DU4L5_HUMAN     AHTGAWGTGLPAPHVPCAPGALPQGAFVSQAARAAPALQPSQAAPAEGVSQPAPARGDFA 240
                          ************************************************:***********

sp|Q9UBX2|DUX4_HUMAN      YAAPAPPDGALSHPQAPRWPPHPGKSREDRDPQRDGLPGPCAVAQPGPAQAGPQGQGVLA 300
sp|P0CJ86|DU4L3_HUMAN     YAAPAPPDGALSHPQAPRWPPHPGKSREDRDPQRDGLPGPCAVAQPGPAQAGPQGQGVLA 300
sp|P0CJ85|DU4L2_HUMAN     YAAPAPPDGALSHPQAPRWPPHPGKSREDRDPQRDGLPGPCAVAQPGPAQAGPQGQGVLA 300
sp|P0CJ89|DU4L6_HUMAN     YAAPAPPDGALSHPQAPRWPPHPGKSREDRDPQRDGLPGPCAVAQPGPAQAGPQGQGVLA 300
sp|P0CJ90|DU4L7_HUMAN     YAAPAPPDGALSHPQAPRWPPHPGKSREDRDPQRDGLPGPCAVAQPGPAQAGPQGQGVLA 300
sp|P0CJ88|DU4L5_HUMAN     YAAPAPPDGALSHPQAPRWPPHPGKSREDRDPQRDGLPGPCAVAQPGPAQAGPQGQGVLA 300
                          ************************************************************

sp|Q9UBX2|DUX4_HUMAN      PPTSQGSPWWGWGRGPQVAGAAWEPQAGAAPPPQPAPPDASASARQGQMQGIPAPSQALQ 360
sp|P0CJ86|DU4L3_HUMAN     PPTSQGSPWWGWGRGPQVAGAAWEPQAGAAPPPQPAPPDASASARQGQMQGIPAPSQALQ 360
sp|P0CJ85|DU4L2_HUMAN     PPTSQGSPWWGWGRGPQVAGAAWEPQAGAAPPPQPAPPDASASARQGQMQGIPAPSQALQ 360
sp|P0CJ89|DU4L6_HUMAN     PPTSQGSPWWGWGRGPQVAGAAWEPQAGAAPPPQPAPPDASASARQGQMQGIPAPSQALQ 360
sp|P0CJ90|DU4L7_HUMAN     PPTSQGSPWWGWGRGPQVAGAAWEPQAGAAPPPQPAPPDASASARQGQMQGIPAPSQALQ 360
sp|P0CJ88|DU4L5_HUMAN     PPTSQGSPWWGWGRGPQVAGAAWEPQAGAAPPPQPAPPDASASARQGQMQGIPAPSQALQ 360
                          ************************************************************

sp|Q9UBX2|DUX4_HUMAN      EPAPWSALPCGLLLDELLASPEFLQQAQPLLETEAPGELEASEEAASLEAPLSEEEYRAL 420
sp|P0CJ86|DU4L3_HUMAN     EPAPWSALPCGLLLDELLASPEFLQQAQPLLETEAPGELEASEEAASLEAPLSEEEYRAL 420
sp|P0CJ85|DU4L2_HUMAN     EPAPWSALPCGLLLDELLASPEFLQQAQPLLETEAPGELEASEEAASLEAPLSEEEYRAL 420
sp|P0CJ89|DU4L6_HUMAN     EPAPWSALPCGLLLDELLASPEFLQQAQPLLETEAPGELEASEEAASLEAPLSEEEYRAL 420
sp|P0CJ90|DU4L7_HUMAN     EPAPWSALPCGLLLDELLASPEFLQQAQPLLETEAPGELEASEEAASLEAPLSEEEYRAL 420
sp|P0CJ88|DU4L5_HUMAN     EPAPWSALPCGLLLDELLASPEFLQQAQPLLETEAPGELEASEEAASLEAPLSEEEYRAL 420
                          ************************************************************

sp|Q9UBX2|DUX4_HUMAN      LEEL 424
sp|P0CJ86|DU4L3_HUMAN     LEEL 424
sp|P0CJ85|DU4L2_HUMAN     LEEL 424
sp|P0CJ89|DU4L6_HUMAN     LEEL 424
sp|P0CJ90|DU4L7_HUMAN     LEEL 424
sp|P0CJ88|DU4L5_HUMAN     LEEL 424
                          ****
```

B

| Accession Number | Q9UBX2 | P0CJ86 | P0CJ85 | P0CJ89 | P0CJ90 | P0CJ88 |
|---|---|---|---|---|---|---|
| **Q9UBX2** | 100% | 99.76% | 99.76% | 99.76% | 99.76% | 99.76% |
| **P0CJ86** | 99.76% | 100% | 100% | 100% | 100% | 100% |
| **P0CJ85** | 99.76% | 100% | 100% | 100% | 100% | 100% |
| **P0CJ89** | 99.76% | 100% | 100% | 100% | 100% | 100% |
| **P0CJ90** | 99.76% | 100% | 100% | 100% | 100% | 100% |
| **P0CJ88** | 99.76% | 100% | 100% | 100% | 100% | 100% |

**Figure S2. Multiple alignment of the Q9UBX2, P0CJ86, P0CJ85, P0CJ89, P0CJ90 and P0GJ88 proteins.**
(**A**) Amino acid sequence alignment of the indicated proteins. *: identical amino acid residues at the same position among protein sequences. Analysis indicates that these proteins bear the same sequences, except position 229, whereat Q9UBX2 contains Isoleucine (I), while all other proteins carry Valine (V). (**B**) Quantification of similarities/identities in between compared protein sequence pairs

A

```
sp|P0C7V4|F90AF_HUMAN      MMARRDPTSWAKRLVRAQTLQKQRRAPVGPRAPPPDEEDPRLKCKNCGAFGHTARSTRCP 60
sp|P0DV74|F90AH_HUMAN      MMARRDPKSWAKRLVRAQTLQKQRRAPVGPRAPPPDEEDPRLKCKNCGAFGHTARSTRCP 60
sp|P0DV73|F90AG_HUMAN      MMARRDPKSWAKRLVRAQTLQKQRRAPVGPRAPPPDEEDPRLKCKNCGAFGHTARSTRCP 60
sp|P0DV75|F90AI_HUMAN      MMARRDPKSWAKRLVRAQTLQKQRRAPVGPRAPPPDEEDPRLKCKNCGAFGHTARSTRCP 60
sp|P0DV76|F90AJ_HUMAN      MMARRDPKSWAKRLVRAQTLQKQRRAPVGPRAPPPDEEDPRLKCKNCGAFGHTARSTRCP 60
                           *******.****************************************************

sp|P0C7V4|F90AF_HUMAN      MKCWKAALVPATLGKKEGKENLKPWKPRVEANPGPLNKDKGEKEERPRQQDPQRKALLHM 120
sp|P0DV74|F90AH_HUMAN      MKCWKAALVPATLGKKEGKENLKPWKPRVEANPGPLNKDKGEKEERPRQQDPQRKALLHM 120
sp|P0DV73|F90AG_HUMAN      MKCWKAALVPATLGKKEGKENLKPWKPRVEANPGPLNKDKGEKEERPRQQDPQRKALLHM 120
sp|P0DV75|F90AI_HUMAN      MKCWKAALVPATLGKKEGKENLKPWKPRVEANPGPLNKDKGEKEERPRQQDPQRKALLHM 120
sp|P0DV76|F90AJ_HUMAN      MKCWKAALVPATLGKKEGKENLKPWKPRVEANPGPLNKDKGEKEERPRQQDPQRKALLHM 120
                           ************************************************************

sp|P0C7V4|F90AF_HUMAN      FSGKPPEKPLPNGKGSTEPSDYLRVASGPMPVHTTSKRPRLDPVLADRSATEMSGRGSVL 180
sp|P0DV74|F90AH_HUMAN      FSGKPPEKPLPNGKGSTESSDHLRVASGPMPVHTTSKRPRVDPVLADRSAAEMSGRGSVL 180
sp|P0DV73|F90AG_HUMAN      FSGKPPEKPLPNGKGSTESSDHLRVASGPMPVHTTSKRPRVDPVLADRSAAEMSGRGSVL 180
sp|P0DV75|F90AI_HUMAN      FSGKPPEKPLPNGKGSTESSDHLRVASGPMPVHTTSKRPRVDPVLADRSAAEMSGRGSVL 180
sp|P0DV76|F90AJ_HUMAN      FSGKPPEKPLPNGKGSTESSDHLRVASGPMPVHTTSKRPRVDPVLADRSAAEMSGRGSVL 180
                           ****************** **:******************:*********:*********

sp|P0C7V4|F90AF_HUMAN      ASLSPLRKASLSSSSSLGPKERQTGAAADMPQPAVRHQGREPLLVVKPTHSRPEGGCREV 240
sp|P0DV74|F90AH_HUMAN      ASLSPLRKASLSSSSSLGPKERQTGAAADIPQPAVRHQGREPLLVVKPTHSRPEGGCREV 240
sp|P0DV73|F90AG_HUMAN      ASLSPLRKASLSSSSSLGPKERQTGAAADIPQPAVRHQGREPLLVVKPTHSRPEGGCREV 240
sp|P0DV75|F90AI_HUMAN      ASLSPLRKASLSSSSSLGPKERQTGAAADMPQPAVRHQGREPLLVVKPTHSSPEGGCREV 240
sp|P0DV76|F90AJ_HUMAN      ASLSPLRKASLSSSSSLGPKERQTGAAADMPQPAVRHQGREPLLVVKPTHSSPEGGCREV 240
                           *****************************:********************* ********

sp|P0C7V4|F90AF_HUMAN      PQAASKTHGLLQAARPQAQDKRPAVTSQPCPPAATHSLGLGSNLSFGPGAKRPAQAPIQA 300
sp|P0DV74|F90AH_HUMAN      PQAASKTHGLLQAARPQAQDKRPAVTSQPCPPAATHSLGLGSNLSFGPGAKRPAQAPIQA 300
sp|P0DV73|F90AG_HUMAN      PQAASKTHGLLQAARPQAQDKRPAVTSQPCPPAATHSLGLGSNLSFGPGAKRPAQAPIQA 300
sp|P0DV75|F90AI_HUMAN      PQAASKTHGLLQAARPQAQDKRPAVTSQPCPPAATHSLGLGSNLSFGPGAKRPAQAPIQA 300
sp|P0DV76|F90AJ_HUMAN      PQAASKTHGLLQAARPQAQDKRPAVTSQPCPPAATHSLGLGSNLSFGPGAKRPAQAPIQA 300
                           ************************************************************

sp|P0C7V4|F90AF_HUMAN      CLNFPKKPRLGPFQIPESAIQGGELGAPENLQPPPAATELGPSTSPQMGRRTPAQVPSVD 360
sp|P0DV74|F90AH_HUMAN      CLNFPKKPRLGPFQIPESAIQGGELGAPENLQPPPAATELGPSTSPQMGRRTPAQVPSVD 360
sp|P0DV73|F90AG_HUMAN      CLNFPKKPRLGPFQIPESAIQGGELGAPENLQPPPAATELGPSTSPQMGRRTPAQVPSVD 360
sp|P0DV75|F90AI_HUMAN      CLKFPKKPRLGPFQIPESAIQGGELGAPGNLQPPPAATELGPSTSPQMGRRTPAQVPSVD 360
sp|P0DV76|F90AJ_HUMAN      CLKFPKKPRLGPFQIPESAIQGGELGAPGNLQPPPAATELGPSTSPQMGRRTPAQVPSVD 360
                           **:************************* *******************************

sp|P0C7V4|F90AF_HUMAN      RQPPHSRPCLPTAQACTMSHHPAASHDGAQPLRVLFRRLENGRWSSSLLAAPSFHSPEKP 420
sp|P0DV74|F90AH_HUMAN      RQPPHSTPCLPTAQACTMSHHSAASHDGAQPLRVLFRRLENGRWSSSLLAAPSFHSPEKP 420
sp|P0DV73|F90AG_HUMAN      RQPPHSTPCLPTAQACTMSHHSAASHDGAQPLRVLFRRLENGRWSSSLLAAPSFHSPEKP 420
sp|P0DV75|F90AI_HUMAN      WQPPHSTPCLPTAQACTMSHHSAASHDGAQPLRVLFRRLENGRWSSSLLAAPSFHSPEKP 420
sp|P0DV76|F90AJ_HUMAN      WQPPHSTPCLPTAQACTMSHHSAASHDGAQPLRVLFRRLENGRWSSSLLAAPSFHSPEKP 420
                            ***** ************** **************************************

sp|P0C7V4|F90AF_HUMAN      GAFLAQSPHVSEKSEAPCVRVPPSVLYEDLQVSSSSEDSDSDLE 464
sp|P0DV74|F90AH_HUMAN      GAFLAQSPHVSEKSEAPCVRVPPSVLYEDLQVSSSSEDSDSDLE 464
sp|P0DV73|F90AG_HUMAN      GAFLAQSPHVSEKSEAPCVRVPPSVLYEDLQVSSSSEDSDSDLE 464
sp|P0DV75|F90AI_HUMAN      GAFLAQSPHVSEKSEAPCVRVPPSVLYEDLQVSSSSEDSDSDLE 464
sp|P0DV76|F90AJ_HUMAN      GAFLAQSPHVSEKSEAPCVRVPPSVLYEDLQVSSSSEDSDSDLE 464
                           ********************************************
```

B

| Accession Number | P0C7V4 | P0DV74 | P0DV73 | P0DV75 | P0DV76 |
|---|---|---|---|---|---|
| **P0C7V4** | 100% | 98.28% | 98.28% | 97.63% | 97.63% |
| **P0DV74** | 98,28% | 100% | 100% | 98.92% | 98.92% |
| **P0DV73** | 98,28% | 100% | 100% | 98.92% | 98.92% |
| **P0DV75** | 97.63% | 98.92% | 98.92% | 100% | 100% |
| **P0DV76** | 97.63% | 98.92% | 98.92% | 100% | 100% |

**Figure S3. Multiple alignment of the P0C7V4, P0DV74, P0DV73, P0DV75 and P0DV76 proteins.**
(**A**) Amino acid sequence alignment of the indicated proteins. *: identical amino acid residues of the protein sequences analyzed. (**B**) Quantification of compared protein similarities/identities.

A

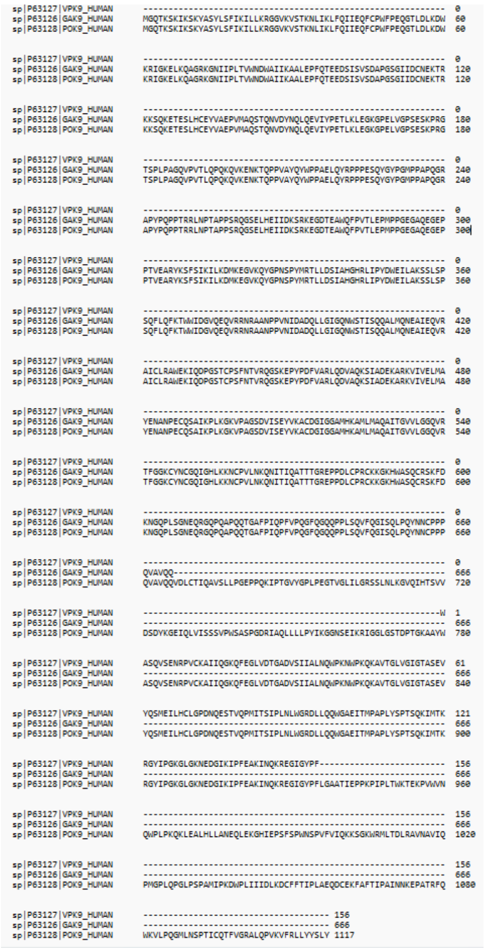

B

| Accession Number | P63127 | P63126 | P63128 |
|---|---|---|---|
| **P63127** | 100% | NAN | 100% |
| **P63126** | NAN | 100% | 100% |
| **P63128** | 100% | 100% | 100% |

**Figure S4. Alignment of the P63127, P63126 and P63128 proteins.** (**A**) Amino acid sequence comparison. Analysis indicates that the P63126 and P63127 proteins are fragments of the P63128 protein. (**B**) Quantification of compared protein similarities/identities. NAN: Not A Number; NAN indicates the lack of homology.

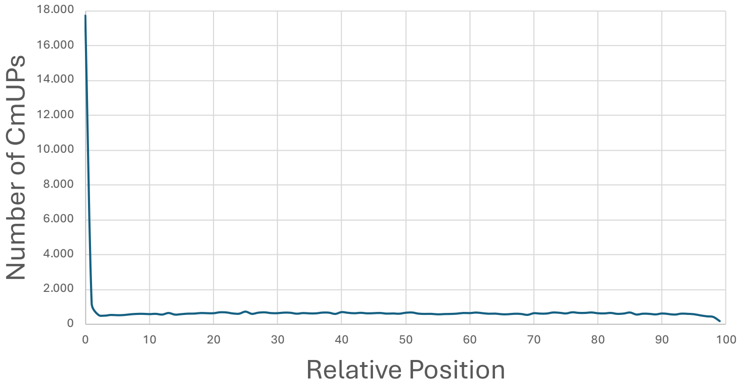

**Figure S5. Relative position of CmUPs in human proteins.**
An accumulation of CmUPs is detected at the $NH_2$-terminus (amino-terminal end, starting at position +1) of the examined proteins. Beyond this position, CmUPs are distributed almost equally within all the different positions of human proteins.

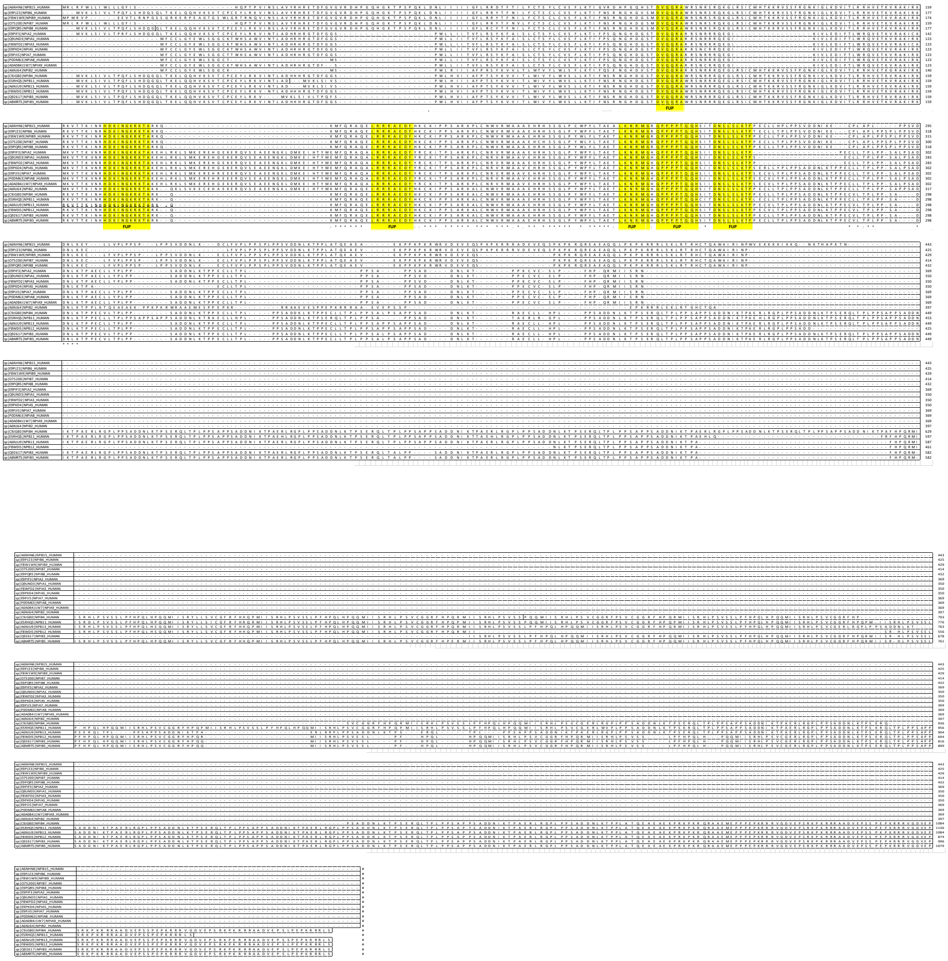

**Figure S6. Multiple alignment of the NPIP protein family.**

The 19-protein members of the NPIP family are aligned. Yellow color indicates the common peptides among all members of the NPIP family. These 6 peptides are characterized as FUPs, since, although they are common between members of the NPIP family, they are unique only for proteins that belong to the NPIP family.

A

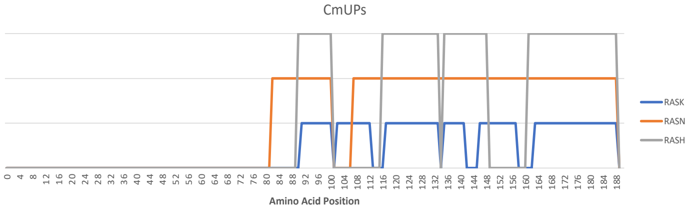

B

| | | |
|---|---|---|
| sp\|P01112\|RASH_HUMAN | M T E Y K L V V V G A G G V G K S A L T I Q L I Q N H F V D E Y D P T I E D S Y R K Q V V I D G E T C L L D I L D T A G | 60 |
| sp\|P01111\|RASN_HUMAN | M T E Y K L V V V G A G G V G K S A L T I Q L I Q N H F V D E Y D P T I E D S Y R K Q V V I D G E T C L L D I L D T A G | 60 |
| sp\|P01116\|RASK_HUMAN | M T E Y K L V V V G A G G V G K S A L T I Q L I Q N H F V D E Y D P T I E D S Y R K Q V V I D G E T C L L D I L D T A G | 60 |

| | | |
|---|---|---|
| sp\|P01112\|RASH_HUMAN | Q E E Y S A M R D Q Y M R T G E G F L C V F A I N N T K S F E D I H Q Y R E Q I K R V K D S D D V P M V L V G N K C D L | 120 |
| sp\|P01111\|RASN_HUMAN | Q E E Y S A M R D Q Y M R T G E G F L C V F A I N N S K S F A D I N L Y R E Q I K R V K D S D D V P M V L V G N K C D L | 120 |
| sp\|P01116\|RASK_HUMAN | Q E E Y S A M R D Q Y M R T G E G F L C V F A I N N T K S F E D I H H Y R E Q I K R V K D S E D V P M V L V G N K C D L | 120 |

FUP

| | | |
|---|---|---|
| sp\|P01112\|RASH_HUMAN | A A R T V E S R Q A Q D L A R S Y G I P Y I E T S A K T R Q G V E D A F Y T L V R E I R Q H K L R K L N P P D E S G P G | 180 |
| sp\|P01111\|RASN_HUMAN | P T R T V D T K Q A H E L A K S Y G I P F I E T S A K T R Q G V E D A F Y T L V R E I R Q Y R M K K L N S S D D G T Q G | 180 |
| sp\|P01116\|RASK_HUMAN | P S R T V D T K Q A Q D L A R S Y G I P F I E T S A K T R Q R V E D A F Y T L V R E I R Q Y R L K K I S K E E - K T P G | 179 |

| | | |
|---|---|---|
| sp\|P01112\|RASH_HUMAN | C M S C K - C V L S | 189 |
| sp\|P01111\|RASN_HUMAN | C M G L P - C V V M | 189 |
| sp\|P01116\|RASK_HUMAN | C V K I K K C I I M | 189 |

**Figure S7. The RAS family.**

(**A**) Schematic presentation of the RAS family members, showing the positions whereat CrUPs and CmUPs are being generated. 1-80 amino acid sequences are identical and CrUPs are being identified within the 81-189 amino acid sequences of the 3 proteins. (**B**) Sequence alignment of the RAS protein family members. Yellow color indicates the "EEYSAM" peptide, which has herein proved to serve as a typical FUP for the RAS family.

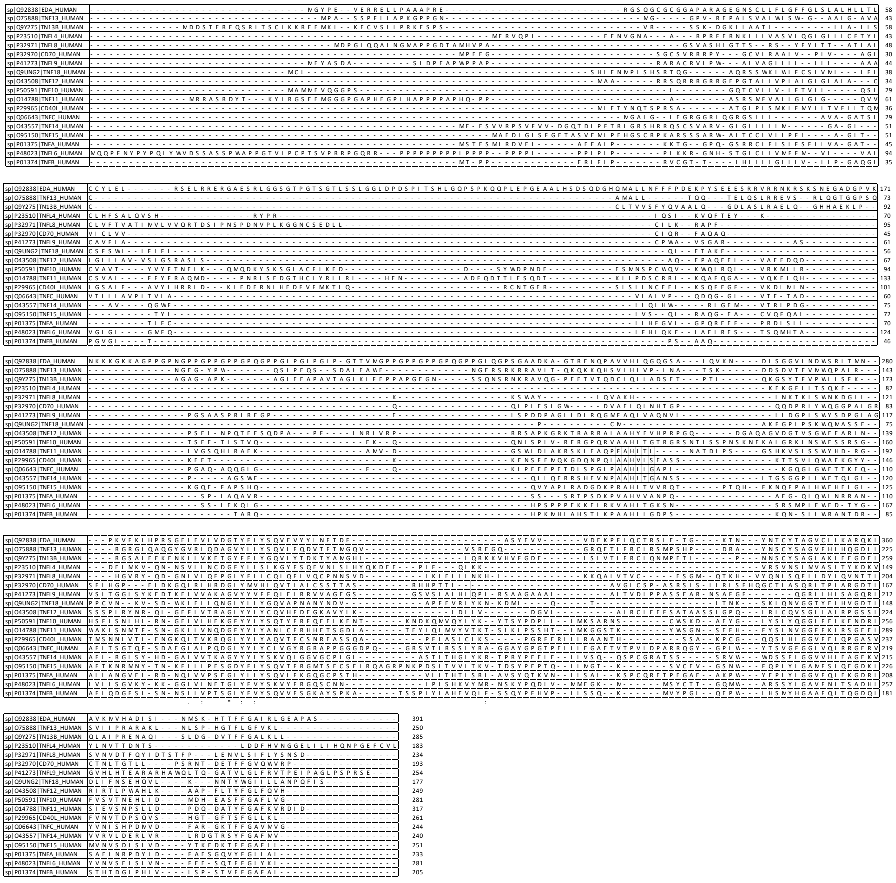

**Figure S8. The TNF family.**

Multiple alignment of the proteins belonging to the TNF family. The TNF family is characterized by complete coverage with CrUPs, since the TNF family members do not show any protein sequence similarities.

**Table S1**. Proteins without Core Unique Peptides (CrUPs)

| Accession Number | Protein Name | Protein Length (Amino Acid number) |
|---|---|---|
| P0DPR3 | T cell receptor delta diversity 1 | 2 |
| P01858 | Phagocytosis-stimulating peptide | 4 |
| P0DPI4 | T cell receptor beta diversity 1 | 4 |
| P0DOY5 | Immunoglobulin heavy diversity 1-1 | 5 |
| A0A0A0MT78 | T cell receptor beta joining 2-7 | 15 |
| P0CJ75 | HTumanin-like 8 | 24 |
| P0DMP1 | Humanin-like 12 | 24 |
| P0CG35 | Thymosin beta-15B | 45 |
| P0DX04 | Thymosin beta-15C | 45 |
| P0CG34 | Thymosin beta-15A | 45 |
| Q3ZM63 | Embryonic testis differentiation protein homolog A | 59 |
| P0DPP9 | Embryonic testis differentiation protein homolog B | 59 |
| P0DMW4 | Small integral membrane protein 10-like protein 2A | 78 |
| P0DMW5 | Small integral membrane protein 10-like protein 2B | 78 |
| P0DP74 | Beta-defensin 130A | 79 |
| A0A1B0GTK5 | Protein FAM236D | 79 |
| P0DP73 | Beta-defensin 130B | 79 |
| P0DP71 | Protein FAM236C | 79 |
| A0A1B0GV22 | Protein FAM236B | 79 |
| A0A1B0GUQ0 | Protein FAM236A | 79 |
| B3EWG5 | Protein FAM25C | 89 |
| B3EWG3 | Protein FAM25A | 89 |
| B3EWG6 | Protein FAM25G | 89 |
| Q69383 | Endogenous retrovirus group K member 6 Rec protein | 105 |
| P61572 | Endogenous retrovirus group K member 19 Rec protein | 105 |
| P61579 | Endogenous retrovirus group K member 25 Rec protein | 105 |
| P61573 | Endogenous retrovirus group K member 9 Rec protein | 105 |
| P0CG04 | Immunoglobulin lambda constant 1 | 106 |
| P0DPH9 | Uncharacterized protein CXorf51B | 108 |
| A0A1B0GTR3 | Uncharacterized protein CXorf51A | 108 |
| Q4V326 | G antigen 2E | 110 |
| A0A0J9YXY3 | T cell receptor beta variable 6-2 | 114 |
| A0A075B6N1 | T cell receptor beta variable 19 | 114 |
| P0DPF7 | T cell receptor beta variable 6-3 | 114 |
| A0JD32 | T cell receptor alpha variable 38-2/delta variable 8 | 116 |
| P01597 | Immunoglobulin kappa variable 1-39 | 117 |
| P01768 | Immunoglobulin heavy variable 3-30 | 117 |
| P01593 | Immunoglobulin kappa variable 1D-33 | 117 |
| P01594 | Immunoglobulin kappa variable 1-33 | 117 |
| P04432 | Immunoglobulin kappa variable 1D-39 | 117 |
| P0CL81 | G antigen 12G | 117 |
| A0A075B6S9 | Probable non-functional immunoglobulinn kappa variable 1-37 | 117 |

**Table S2.** Number of CrUPs, according to their length (number of amino acids) and their percentage (%) to the total number of CrUPs

| Amino Acids Number | CrUPs (number) | Percentage | | Amino Acids Number | CrUPs (number) | Percentage |
|---|---|---|---|---|---|---|
| 4 | 759 | 0.01% | | 53 | 13 | 0.00% |
| 5 | 626,878 | 8.62% | | 54 | 16 | 0.00% |
| 6 | 5,019,502 | 69.03% | | 55 | 18 | 0.00% |
| 7 | 1,498,004 | 20.60% | | 56 | 14 | 0.00% |
| 8 | 86,039 | 1.18% | | 57 | 9 | 0.00% |
| 9 | 14,939 | 0.21% | | 58 | 13 | 0.00% |
| 10 | 6,987 | 0.10% | | 59 | 7 | 0.00% |
| 11 | 4,628 | 0.06% | | 60 | 7 | 0.00% |
| 12 | 2,972 | 0.04% | | 61 | 10 | 0.00% |
| 13 | 2,427 | 0.03% | | 62 | 5 | 0.00% |
| 14 | 1,598 | 0.02% | | 63 | 18 | 0.00% |
| 15 | 1,239 | 0.02% | | 64 | 7 | 0.00% |
| 16 | 1,031 | 0.01% | | 65 | 7 | 0.00% |
| 17 | 635 | 0.01% | | 66 | 6 | 0.00% |
| 18 | 658 | 0.01% | | 67 | 6 | 0.00% |
| 19 | 473 | 0.01% | | 68 | 9 | 0.00% |
| 20 | 410 | 0.01% | | 69 | 5 | 0.00% |
| 21 | 214 | 0.00% | | 70 | 8 | 0.00% |
| 22 | 237 | 0.00% | | 71 | 3 | 0.00% |
| 23 | 220 | 0.00% | | 72 | 4 | 0.00% |
| 24 | 152 | 0.00% | | 73 | 6 | 0.00% |
| 25 | 136 | 0.00% | | 74 | 7 | 0.00% |
| 26 | 121 | 0.00% | | 75 | 9 | 0.00% |
| 27 | 105 | 0.00% | | 76 | 3 | 0.00% |
| 28 | 96 | 0.00% | | 77 | 5 | 0.00% |
| 29 | 76 | 0.00% | | 78 | 6 | 0.00% |
| 30 | 83 | 0.00% | | 79 | 4 | 0.00% |
| 31 | 62 | 0.00% | | 80 | 4 | 0.00% |
| 32 | 59 | 0.00% | | 81 | 4 | 0.00% |
| 33 | 62 | 0.00% | | 82 | 1 | 0.00% |
| 34 | 49 | 0.00% | | 83 | 3 | 0.00% |
| 35 | 37 | 0.00% | | 84 | 10 | 0.00% |
| 36 | 37 | 0.00% | | 85 | 4 | 0.00% |
| 37 | 35 | 0.00% | | 86 | 6 | 0.00% |
| 38 | 37 | 0.00% | | 87 | 8 | 0.00% |
| 39 | 35 | 0.00% | | 88 | 3 | 0.00% |
| 40 | 22 | 0.00% | | 89 | 4 | 0.00% |
| 41 | 25 | 0.00% | | 90 | 4 | 0.00% |
| 42 | 26 | 0.00% | | 91 | 3 | 0.00% |
| 43 | 24 | 0.00% | | 92 | 3 | 0.00% |
| 44 | 17 | 0.00% | | 93 | 3 | 0.00% |
| 45 | 21 | 0.00% | | 94 | 5 | 0.00% |
| 46 | 18 | 0.00% | | 95 | 3 | 0.00% |
| 47 | 22 | 0.00% | | 96 | 6 | 0.00% |
| 48 | 16 | 0.00% | | 97 | 3 | 0.00% |
| 49 | 26 | 0.00% | | 98 | 3 | 0.00% |
| 50 | 25 | 0.00% | | 99 | 0 | 0.00% |
| 51 | 13 | 0.00% | | 100 | 3 | 0.00% |
| 52 | 12 | 0.00% | | | | |

**Table S3.** Number of CmUPs, according to their length (number of amino acids) and their percentage (%) to the total number of CmUPs

| Amino Acids Number | CmUPs (number) | Percentage | | Amino Acids Number | CmUPs (number) | Percentage |
|---|---|---|---|---|---|---|
| 6 | 16 | 0.02% | | 53 | 309 | 0.47% |
| 7 | 194 | 0.29% | | 54 | 286 | 0.43% |
| 8 | 282 | 0.42% | | 55 | 301 | 0.45% |
| 9 | 414 | 0.62% | | 56 | 255 | 0.38% |
| 10 | 1,204 | 1.81% | | 57 | 286 | 0.43% |
| 11 | 3,796 | 5.71% | | 58 | 282 | 0.42% |
| 12 | 3,034 | 4.57% | | 59 | 304 | 0.46% |
| 13 | 1,699 | 2.56% | | 60 | 267 | 0.40% |
| 14 | 1,147 | 1.73% | | 61 | 216 | 0.33% |
| 15 | 1,021 | 1.54% | | 62 | 258 | 0.39% |
| 16 | 1,024 | 1.54% | | 63 | 227 | 0.34% |
| 17 | 966 | 1.45% | | 64 | 216 | 0.33% |
| 18 | 958 | 1.44% | | 65 | 213 | 0.32% |
| 19 | 951 | 1.43% | | 66 | 195 | 0.29% |
| 20 | 851 | 1.28% | | 67 | 225 | 0.34% |
| 21 | 892 | 1.34% | | 68 | 211 | 0.32% |
| 22 | 845 | 1.27% | | 69 | 197 | 0.30% |
| 23 | 778 | 1.17% | | 70 | 203 | 0.31% |
| 24 | 716 | 1.08% | | 71 | 205 | 0.31% |
| 25 | 697 | 1.05% | | 72 | 193 | 0.29% |
| 26 | 669 | 1.01% | | 73 | 189 | 0.28% |
| 27 | 628 | 0.95% | | 74 | 199 | 0.30% |
| 28 | 608 | 0.92% | | 75 | 184 | 0.28% |
| 29 | 638 | 0.96% | | 76 | 203 | 0.31% |
| 30 | 565 | 0.85% | | 77 | 179 | 0.27% |
| 31 | 557 | 0.84% | | 78 | 204 | 0.31% |
| 32 | 492 | 0.74% | | 79 | 177 | 0.27% |
| 33 | 499 | 0.75% | | 80 | 165 | 0.25% |
| 34 | 497 | 0.75% | | 81 | 197 | 0.30% |
| 35 | 502 | 0.76% | | 82 | 165 | 0.25% |
| 36 | 451 | 0.68% | | 83 | 172 | 0.26% |
| 37 | 460 | 0.69% | | 84 | 165 | 0.25% |
| 38 | 453 | 0.68% | | 85 | 154 | 0.23% |
| 39 | 402 | 0.60% | | 86 | 151 | 0.23% |
| 40 | 398 | 0.60% | | 87 | 168 | 0.25% |
| 41 | 427 | 0.64% | | 88 | 155 | 0.23% |
| 42 | 356 | 0.54% | | 89 | 147 | 0.22% |
| 43 | 348 | 0.52% | | 90 | 144 | 0.22% |
| 44 | 352 | 0.53% | | 91 | 145 | 0.22% |
| 45 | 377 | 0.57% | | 92 | 126 | 0.19% |
| 46 | 355 | 0.53% | | 93 | 144 | 0.22% |
| 47 | 313 | 0.47% | | 94 | 137 | 0.21% |
| 48 | 310 | 0.47% | | 95 | 131 | 0.20% |
| 49 | 302 | 0.45% | | 96 | 116 | 0.17% |
| 50 | 283 | 0.43% | | 97 | 140 | 0.21% |
| 51 | 279 | 0.42% | | 98 | 119 | 0.18% |
| 52 | 262 | 0.39% | | 99 | 141 | 0.21% |
| 51 | 13 | 0.00% | | 100 | 132 | 0.20% |
| 52 | 12 | 0.00% | | | | |

**Table S4.** Number of CrUPs per CmUP assembly

| Number of CrUPs per CmUP | CmUPs (number) | Percentage (%) | | Number of CrUPs per CmUP | CmUPs (number) | Percentage (%) |
|---|---|---|---|---|---|---|
| 2 | 908 | 1.37% | | 52 | 218 | 0.33% |
| 3 | 945 | 1.42% | | 53 | 236 | 0.36% |
| 4 | 2,476 | 3.73% | | 54 | 213 | 0.32% |
| 5 | 5,086 | 7.65% | | 55 | 225 | 0.34% |
| 6 | 3,355 | 5.05% | | 56 | 205 | 0.31% |
| 7 | 1,601 | 2.41% | | 57 | 207 | 0.31% |
| 8 | 1,535 | 2.31% | | 58 | 179 | 0.27% |
| 9 | 1,552 | 2.34% | | 59 | 185 | 0.28% |
| 10 | 1,532 | 2.31% | | 60 | 205 | 0.31% |
| 11 | 1,452 | 2.19% | | 61 | 176 | 0.26% |
| 12 | 1,258 | 1.89% | | 62 | 181 | 0.27% |
| 13 | 1,056 | 1.59% | | 63 | 184 | 0.28% |
| 14 | 983 | 1.48% | | 64 | 191 | 0.29% |
| 15 | 936 | 1.41% | | 65 | 198 | 0.30% |
| 16 | 849 | 1.28% | | 66 | 181 | 0.27% |
| 17 | 750 | 1.13% | | 67 | 169 | 0.25% |
| 18 | 774 | 1.16% | | 68 | 158 | 0.24% |
| 19 | 715 | 1.08% | | 69 | 157 | 0.24% |
| 20 | 670 | 1.01% | | 70 | 178 | 0.27% |
| 21 | 631 | 0.95% | | 71 | 153 | 0.23% |
| 22 | 607 | 0.91% | | 72 | 163 | 0.25% |
| 23 | 605 | 0.91% | | 73 | 176 | 0.26% |
| 24 | 554 | 0.83% | | 74 | 175 | 0.26% |
| 25 | 515 | 0.78% | | 75 | 159 | 0.24% |
| 26 | 498 | 0.75% | | 76 | 157 | 0.24% |
| 27 | 460 | 0.69% | | 77 | 149 | 0.22% |
| 28 | 421 | 0.63% | | 78 | 170 | 0.26% |
| 29 | 394 | 0.59% | | 79 | 155 | 0.23% |
| 30 | 417 | 0.63% | | 80 | 139 | 0.21% |
| 31 | 412 | 0.62% | | 81 | 139 | 0.21% |
| 32 | 365 | 0.55% | | 82 | 135 | 0.20% |
| 33 | 393 | 0.59% | | 83 | 144 | 0.22% |
| 34 | 318 | 0.48% | | 84 | 156 | 0.23% |
| 35 | 343 | 0.52% | | 85 | 142 | 0.21% |
| 36 | 336 | 0.51% | | 86 | 146 | 0.22% |
| 37 | 314 | 0.47% | | 87 | 129 | 0.19% |
| 38 | 332 | 0.50% | | 88 | 136 | 0.20% |
| 39 | 345 | 0.52% | | 89 | 157 | 0.24% |
| 40 | 331 | 0.50% | | 90 | 135 | 0.20% |
| 41 | 303 | 0.46% | | 91 | 114 | 0.17% |
| 42 | 307 | 0.46% | | 92 | 143 | 0.22% |
| 43 | 274 | 0.41% | | 93 | 107 | 0.16% |
| 44 | 258 | 0.39% | | 94 | 122 | 0.18% |
| 45 | 277 | 0.42% | | 95 | 128 | 0.19% |
| 46 | 264 | 0.40% | | 96 | 121 | 0.18% |
| 47 | 274 | 0.41% | | 97 | 139 | 0.21% |
| 48 | 233 | 0.35% | | 98 | 119 | 0.18% |
| 49 | 259 | 0.39% | | 99 | 119 | 0.18% |
| 50 | 250 | 0.38% | | 100 | 121 | 0.18% |
| 51 | 228 | 0.34% | | | | |

**Table S5.** Distribution of CrUPs and CmUPs on human chromosomes

| **Chromosomes** | **Reviewed Proteins (number)** | **CrUPs (number)** | **CmUPs (number)** | **CrUP Density (%)** | **CmUP Density (%)** | **Coverage by CrUPs (%)** | **Proteins without CrUPs (number)** | **Proteins without CrUPs (%)** |
|---|---|---|---|---|---|---|---|---|
| Chromosome 1 | 1,991 | 718,473 | 6,228 | 64% | 0.55% | 93% | 13 | 0.65% |
| Chromosome 2 | 1,262 | 532,490 | 4,116 | 65% | 0.50% | 93% | 15 | 1.19% |
| Chromosome 3 | 1,038 | 429,359 | 3,248 | 67% | 0.51% | 96% | 1 | 0.10% |
| Chromosome 4 | 734 | 292,476 | 2,336 | 66% | 0.53% | 94% | 6 | 0.82% |
| Chromosome 5 | 862 | 328,639 | 3,339 | 63% | 0.64% | 92% | 5 | 0.58% |
| Chromosome 6 | 979 | 369,853 | 2,844 | 66% | 0.51% | 95% | 5 | 0.51% |
| Chromosome 7 | 955 | 328,581 | 3,129 | 62% | 0.60% | 92% | 18 | 1.88% |
| Chromosome 8 | 669 | 246,557 | 2,034 | 65% | 0.54% | 94% | 9 | 1.35% |
| Chromosome 9 | 746 | 284,972 | 2,401 | 64% | 0.54% | 91% | 0 | 0.00% |
| Chromosome 10 | 714 | 282,011 | 2,279 | 65% | 0.53% | 94% | 5 | 0.70% |
| Chromosome 11 | 1,263 | 426,576 | 3,784 | 65% | 0.58% | 95% | 2 | 0.16% |
| Chromosome 12 | 991 | 371,118 | 3,536 | 65% | 0.62% | 94% | 0 | 0.00% |
| Chromosome 13 | 313 | 136,156 | 1,104 | 67% | 0.55% | 95% | 6 | 1.92% |
| Chromosome 14 | 704 | 236,879 | 2,104 | 65% | 0.58% | 94% | 5 | 0.71% |
| Chromosome 15 | 569 | 247,781 | 2,030 | 64% | 0.52% | 91% | 2 | 0.35% |
| Chromosome 16 | 805 | 300,095 | 2,170 | 65% | 0.47% | 93% | 7 | 0.87% |
| Chromosome 17 | 1,132 | 402,250 | 3,919 | 63% | 0.62% | 92% | 6 | 0.53% |
| Chromosome 18 | 261 | 116,977 | 916 | 68% | 0.53% | 96% | 0 | 0.00% |
| Chromosome 19 | 1,385 | 432,058 | 5,203 | 59% | 0.72% | 93% | 3 | 0.22% |
| Chromosome 20 | 527 | 176,951 | 1,558 | 67% | 0.59% | 96% | 0 | 0.00% |
| Chromosome 21 | 215 | 69,956 | 584 | 65% | 0.54% | 94% | 0 | 0.00% |
| Chromosome 22 | 469 | 148,071 | 1,398 | 63% | 0.60% | 91% | 0 | 0.00% |
| Chromosome X | 803 | 257,257 | 3,148 | 61% | 0.75% | 90% | 28 | 3.49% |
| Chromosome Y | 38 | 3,615 | 275 | 18% | 1.37% | 32% | 6 | 15.79% |
| Mitochondrion | 15 | 2,821 | 15 | 74% | 0.39% | 100% | 0 | 0.00% |

In each human chromosome, the number of reviewed proteins, the number of CrUPs and the number of CmUPs are shown. Table, also, describes the density (%) of CrUPs and CmUPs, and the coverage (%) of each chromosome by CrUPS. The number of proteins per chromosome that do not contain CrUPs, and their percentage to the reviewed proteins, are also presented.

**Table S6.** Analysis of human protein families by CrUP profiling

| Family Name | Proteins per Family (number) | CrUPs per Family (number) | Density of CrUPs per Family (%) | Family Coverage by CrUPs (%) |
|---|---|---|---|---|
| G-protein coupled receptor 1 family | 724 | 137,161 | 54% | 88% |
| Krueppel C2H2-type zinc-finger protein family | 543 | 170,767 | 50% | 94% |
| Protein kinase superfamily | 491 | 265,955 | 63% | 93% |
| Small GTPase superfamily | 162 | 19,838 | 52% | 82% |
| Immunoglobulin superfamily | 130 | 44,970 | 59% | 90% |
| Peptidase S1 family | 119 | 33,097 | 63% | 93% |
| Protein-tyrosine phosphatase family | 93 | 47,978 | 66% | 96% |
| Tyr protein kinase family | 90 | 55,715 | 65% | 96% |
| TRAFAC class myosin-kinesin ATPase superfamily | 85 | 74,902 | 59% | 90% |
| Major facilitator superfamily | 81 | 28,347 | 67% | 96% |
| CAMK Ser/Thr protein kinase family | 75 | 59,158 | 63% | 94% |
| Peptidase C19 family | 75 | 37,070 | 55% | 80% |
| Intermediate filament family | 75 | 19,566 | 48% | 83% |
| Rab family | 74 | 9,262 | 51% | 81% |
| Ser/Thr protein kinase family | 67 | 34,552 | 67% | 95% |
| TRIM/RBCC family | 65 | 20,811 | 60% | 87% |
| Class I-like SAM-binding methyltransferase superfamily | 62 | 19,805 | 69% | 95% |
| CMGC Ser/Thr protein kinase family | 62 | 19,831 | 56% | 86% |
| DEAD box helicase family | 60 | 31,097 | 63% | 91% |
| Cytochrome P450 family | 60 | 16,195 | 54% | 84% |
| AGC Ser/Thr protein kinase family | 58 | 26,169 | 56% | 87% |
| STE Ser/Thr protein kinase family | 56 | 27,858 | 60% | 91% |
| BZIP family | 55 | 13,492 | 66% | 97% |
| Short-chain dehydrogenases/reductases (SDR) family | 54 | 11,644 | 68% | 96% |
| Mitochondrial carrier (TC 2,A,29) family | 53 | 11,063 | 61% | 91% |
| AB hydrolase superfamily | 51 | 13,908 | 68% | 96% |
| G-protein coupled receptor 2 family | 49 | 37,118 | 69% | 97% |
| ABC transporter superfamily | 48 | 42,703 | 67% | 97% |
| Nuclear hormone receptor family | 48 | 14,346 | 58% | 88% |
| Class V-like SAM-binding methyltransferase superfamily | 46 | 40,268 | 69% | 98% |
| Ligand-gated ion channel (TC 1,A,9) family | 46 | 11,787 | 53% | 84% |
| Kinesin family | 45 | 36,421 | 65% | 94% |
| Paired homeobox family | 44 | 7,803 | 57% | 86% |
| Potassium channel family | 44 | 18,153 | 55% | 87% |
| Methyltransferase superfamily | 43 | 10,351 | 66% | 92% |
| Ubiquitin-conjugating enzyme family | 43 | 9,251 | 63% | 89% |
| Serpin family | 38 | 10,406 | 68% | 96% |

| | | | | |
|---|---|---|---|---|
| Calycin superfamily | 38 | 3,858 | 62% | 90% |
| TGF-beta family | 37 | 8,819 | 61% | 89% |
| Myosin family | 37 | 35,246 | 54% | 86% |
| Cation transport ATPase (P-type) (TC 3,A,3) family | 36 | 23,404 | 55% | 87% |
| AAA ATPase family | 35 | 19,409 | 67% | 95% |
| Beta-defensin family | 35 | 1,860 | 63% | 89% |
| TKL Ser/Thr protein kinase family | 34 | 16,565 | 67% | 96% |
| Type I cytokine receptor family | 32 | 14,355 | 74% | 100% |
| Cyclin family | 32 | 9,706 | 67% | 96% |
| SNF2/RAD54 helicase family | 32 | 33,819 | 62% | 92% |
| Sulfotransferase 1 family | 32 | 7,292 | 53% | 82% |
| Tetraspanin (TM4SF) family | 31 | 5,524 | 70% | 98% |
| Sorting nexin family | 31 | 10,107 | 69% | 97% |
| TRAFAC class dynamin-like GTPase superfamily | 30 | 12,299 | 59% | 90% |
| ETS family | 29 | 7,535 | 62% | 92% |
| STE20 subfamily | 29 | 13,008 | 56% | 87% |
| CDC2/CDKX subfamily | 29 | 8,521 | 54% | 84% |
| Arf family | 28 | 3,509 | 58% | 88% |
| Actin family | 28 | 4,909 | 44% | 68% |
| Non-receptor class dual specificity subfamily | 27 | 6,115 | 67% | 97% |
| ATP-dependent AMP-binding enzyme family | 27 | 11,581 | 65% | 93% |
| G-protein coupled receptor T2R family | 27 | 4,615 | 55% | 89% |
| GAGE family | 27 | 605 | 20% | 43% |
| Adhesion G-protein coupled receptor (ADGR) subfamily | 26 | 23,105 | 70% | 98% |
| Glycosyltransferase 2 family | 26 | 9,448 | 65% | 95% |
| Splicing factor SR family | 26 | 7,161 | 62% | 96% |
| Intercrine beta (chemokine CC) family | 26 | 1,654 | 61% | 90% |
| Antp homeobox family | 26 | 4,316 | 57% | 89% |
| Ras family | 26 | 2,524 | 47% | 76% |
| PRAME family | 26 | 2,302 | 19% | 41% |
| Metallo-dependent hydrolases superfamily | 24 | 9,132 | 66% | 96% |
| Nudix hydrolase family | 24 | 3,943 | 65% | 89% |
| DHHC palmitoyltransferase family | 24 | 6,281 | 64% | 92% |
| Claudin family | 24 | 3,411 | 63% | 92% |
| TRAFAC class TrmE-Era-EngA-EngB-Septin-like GTPase superfamily | 24 | 5,749 | 60% | 93% |
| Tubulin family | 24 | 1,850 | 17% | 42% |
| Helicase family | 23 | 20,260 | 72% | 99% |
| Glycosyltransferase 31 family | 23 | 6,115 | 71% | 99% |
| Peptidase M14 family | 23 | 10,659 | 69% | 99% |
| Major facilitator (TC 2,A,1) superfamily | 23 | 8,659 | 68% | 97% |
| Organic cation transporter (TC 2,A,1,19) family | 23 | 8,659 | 68% | 97% |

| | | | | |
|---|---|---|---|---|
| S-100 family | 23 | 2,104 | 68% | 96% |
| Peptidase M10A family | 23 | 7,912 | 67% | 97% |
| GST superfamily | 23 | 2,818 | 50% | 77% |
| Speedy/Ringo family | 23 | 745 | 10% | 28% |
| Lipase family | 22 | 7,182 | 69% | 99% |
| G-protein coupled receptor 3 family | 22 | 13,279 | 66% | 96% |
| DEAH subfamily | 22 | 16,111 | 65% | 93% |
| Heparin-binding growth factors family | 22 | 3,120 | 64% | 96% |
| Transient receptor (TC 1,A,4) family | 22 | 13,331 | 62% | 92% |
| TRAFAC class translation factor GTPase superfamily | 22 | 9,502 | 61% | 84% |
| CEA family | 22 | 3,425 | 40% | 79% |
| UDP-glycosyltransferase family | 22 | 3,333 | 29% | 56% |
| Histone-lysine methyltransferase family | 21 | 24,426 | 69% | 98% |
| Connexin family | 21 | 4,528 | 64% | 96% |
| Lipocalin family | 21 | 2,463 | 61% | 87% |
| Cyclic nucleotide phosphodiesterase family | 21 | 10,101 | 61% | 93% |
| Cyclophilin-type PPIase family | 21 | 2,989 | 54% | 82% |
| Nucleosome assembly protein (NAP) family | 21 | 3,271 | 44% | 68% |
| Histone H2B family | 21 | 475 | 17% | 41% |
| FAM90 family | 21 | 761 | 8% | 25% |
| USP17 subfamily | 21 | 365 | 3% | 17% |
| Glycosyltransferase 29 family | 20 | 5,375 | 71% | 99% |
| PI3/PI4-kinase family | 20 | 24,378 | 71% | 96% |
| Semaphorin family | 20 | 11,747 | 69% | 98% |
| GalNAc-T subfamily | 20 | 7,675 | 63% | 95% |
| Classic translation factor GTPase family | 20 | 8,002 | 59% | 82% |
| Rho family | 20 | 2,529 | 49% | 79% |
| Class-I aminoacyl-tRNA synthetase family | 19 | 11,320 | 74% | 100% |
| Class-II aminoacyl-tRNA synthetase family | 19 | 8,081 | 70% | 97% |
| PMP-22/EMP/MP20 family | 19 | 2,968 | 67% | 97% |
| Acetyltransferase family | 19 | 4,753 | 66% | 94% |
| Adenylyl cyclase class-4/guanylyl cyclase family | 19 | 13,172 | 65% | 94% |
| Sodium:neurotransmitter symporter (SNF) (TC 2,A,22) family | 19 | 7,926 | 64% | 96% |
| Sp1 C2H2-type zinc-finger protein family | 19 | 5,524 | 63% | 95% |
| Wnt family | 19 | 4,101 | 59% | 93% |
| Synaptotagmin family | 19 | 4,974 | 58% | 87% |
| NR1 subfamily | 19 | 5,222 | 58% | 89% |
| BTN/MOG family | 19 | 4,634 | 56% | 89% |
| Gamma-aminobutyric acid receptor (TC 1,A,9,5) subfamily | 19 | 4,647 | 51% | 82% |
| LCE family | 19 | 482 | 24% | 61% |
| Histone H2A family | 19 | 434 | 16% | 50% |

| | | | | |
|---|---|---|---|---|
| NPIP family | 19 | 348 | 3% | 16% |
| Tumor necrosis factor family | 18 | 3,301 | 73% | 100% |
| Ankyrin SOCS box (ASB) family | 18 | 5,211 | 71% | 100% |
| Integrin alpha chain family | 18 | 14,203 | 70% | 98% |
| MAP kinase kinase kinase subfamily | 18 | 11,816 | 67% | 96% |
| HAD-like hydrolase superfamily | 18 | 3,824 | 65% | 94% |
| Glutamate-gated ion channel (TC 1,A,10,1) family | 18 | 10,840 | 57% | 87% |
| NLRP family | 17 | 13,083 | 71% | 98% |
| Small leucine-rich proteoglycan (SLRP) family | 17 | 5,173 | 70% | 100% |
| PP2C family | 17 | 5,744 | 70% | 98% |
| ARTD/PARP family | 17 | 10,307 | 68% | 96% |
| MAGUK family | 17 | 10,649 | 66% | 96% |
| Sulfatase family | 17 | 6,630 | 66% | 97% |
| Fatty-acid binding protein (FABP) family | 17 | 1,395 | 62% | 95% |
| Intercrine alpha (chemokine CxC) family | 17 | 1,140 | 57% | 89% |
| Heat shock protein 70 family | 17 | 5,572 | 49% | 76% |
| G-alpha family | 17 | 2,742 | 40% | 71% |
| GOLGA6 family | 17 | 933 | 8% | 30% |
| Bcl-2 family | 16 | 2,799 | 72% | 99% |
| MS4A family | 16 | 2,988 | 71% | 99% |
| Dynein heavy chain family | 16 | 46,996 | 69% | 97% |
| Alpha-carbonic anhydrase family | 16 | 3,320 | 69% | 99% |
| Syntaxin family | 16 | 3,111 | 67% | 96% |
| Amino acid/polyamine transporter 2 family | 16 | 5,628 | 67% | 96% |
| Aldehyde dehydrogenase family | 16 | 5,335 | 64% | 96% |
| CD225/Dispanin family | 16 | 1,876 | 62% | 93% |
| Abd-B homeobox family | 16 | 3,080 | 61% | 92% |
| Zinc-containing alcohol dehydrogenase family | 16 | 3,582 | 60% | 89% |
| Acetylcholine receptor (TC 1,A,9,1) subfamily | 16 | 4,609 | 58% | 88% |
| Cytidine and deoxycytidylate deaminase family | 16 | 2,568 | 57% | 83% |
| Inward rectifier-type potassium channel (TC 1,A,2,1) family | 16 | 3,502 | 53% | 83% |
| POU transcription factor family | 16 | 3,537 | 51% | 81% |
| MIP/aquaporin (TC 1,A,8) family | 16 | 2,351 | 50% | 78% |
| MHC class I family | 16 | 2,508 | 49% | 83% |
| Alpha/beta interferon family | 16 | 1,072 | 35% | 72% |
| NBPF family | 16 | 1,149 | 5% | 35% |
| Beta/gamma-crystallin family | 15 | 6,168 | 71% | 98% |
| Peptidase C1 family | 15 | 3,900 | 70% | 98% |
| Peptidase C2 family | 15 | 8,374 | 70% | 99% |
| SnRNP Sm proteins family | 15 | 1,205 | 68% | 95% |
| LDLR family | 15 | 16,397 | 67% | 98% |
| Non-receptor class myotubularin subfamily | 15 | 8,999 | 67% | 97% |

| | | | | |
|---|---|---|---|---|
| Two pore domain potassium channel (TC 1,A,1,8) family | 15 | 3,782 | 66% | 96% |
| Kallikrein subfamily | 15 | 2,549 | 65% | 98% |
| Cystatin family | 15 | 1,301 | 63% | 92% |
| Amino acid-polyamine-organocation (APC) superfamily | 15 | 4,739 | 61% | 94% |
| Type-B carboxylesterase/lipase family | 15 | 6,665 | 55% | 84% |
| SIGLEC (sialic acid binding Ig-like lectin) family | 15 | 4,943 | 54% | 85% |
| Aldo/keto reductase family | 15 | 1,822 | 39% | 70% |
| GOLGA8 family | 15 | 390 | 4% | 33% |
| Type 2 subfamily | 14 | 7,741 | 75% | 100% |
| BPI/LBP/Plunc superfamily | 14 | 4,383 | 75% | 100% |
| ZIP transporter (TC 2,A,5) family | 14 | 4,819 | 70% | 98% |
| Monocarboxylate porter (TC 2,A,1,13) family | 14 | 4,824 | 69% | 99% |
| Monovalent cation:proton antiporter 1 (CPA1) transporter (TC 2,A,36) family | 14 | 6,785 | 65% | 94% |
| Sugar transporter (TC 2,A,1,1) family | 14 | 4,676 | 63% | 91% |
| TCP-1 chaperonin family | 14 | 4,978 | 62% | 87% |
| Type IV subfamily | 14 | 10,641 | 61% | 93% |
| SNF1 subfamily | 14 | 6,006 | 57% | 86% |
| Ephrin receptor subfamily | 14 | 7,941 | 57% | 93% |
| Dynamin/Fzo/YdjA family | 14 | 5,488 | 56% | 87% |
| G protein gamma family | 14 | 532 | 54% | 90% |
| MAP kinase subfamily | 14 | 3,610 | 54% | 85% |
| Annexin family | 14 | 2,809 | 53% | 77% |
| MHC class II family | 14 | 1,885 | 52% | 85% |
| Humanin family | 14 | 93 | 27% | 69% |
| Metallothionein superfamily | 14 | 219 | 26% | 70% |
| Type 1 family | 14 | 219 | 26% | 70% |
| SPATA31 family | 14 | 3,114 | 19% | 37% |
| TAF11 family | 14 | 351 | 13% | 32% |
| Peptidase C14A family | 13 | 3,448 | 69% | 96% |
| Pancreatic ribonuclease family | 13 | 1,467 | 68% | 98% |
| ATF subfamily | 13 | 3,516 | 67% | 97% |
| Peptidase M1 family | 13 | 7,382 | 67% | 92% |
| Formin homology family | 13 | 10,335 | 65% | 96% |
| Glucose transporter subfamily | 13 | 4,186 | 62% | 90% |
| Ov-serpin subfamily | 13 | 3,019 | 60% | 90% |
| Organo anion transporter (TC 2,A,60) family | 13 | 5,377 | 59% | 83% |
| Septin GTPase family | 13 | 2,941 | 53% | 88% |
| CaMK subfamily | 13 | 3,473 | 50% | 84% |
| PPP phosphatase family | 13 | 2,470 | 44% | 71% |
| KRTAP type 10 family | 13 | 912 | 24% | 71% |
| Beta type-B retroviral envelope protein family | 13 | 753 | 10% | 36% |

| | | | | |
|---|---|---|---|---|
| Phospholipase A2 family | 12 | 1,822 | 71% | 100% |
| RBR family | 12 | 5,191 | 70% | 97% |
| 1-acyl-sn-glycerol-3-phosphate acyltransferase family | 12 | 3,434 | 69% | 98% |
| PA-phosphatase related phosphoesterase family | 12 | 3,044 | 68% | 98% |
| ABCC family | 12 | 12,023 | 68% | 97% |
| CRISP family | 12 | 2,503 | 67% | 99% |
| Insulin receptor subfamily | 12 | 9,530 | 67% | 96% |
| SNF7 family | 12 | 1,831 | 66% | 95% |
| Sodium:solute symporter (SSF) (TC 2,A,21) family | 12 | 5,058 | 65% | 97% |
| OSBP family | 12 | 6,320 | 64% | 95% |
| GLI C2H2-type zinc-finger protein family | 12 | 5,566 | 60% | 92% |
| TGFB receptor subfamily | 12 | 4,003 | 59% | 91% |
| Bicoid subfamily | 12 | 2,268 | 58% | 90% |
| Gamma type-C retroviral envelope protein family | 12 | 3,821 | 58% | 82% |
| PKC subfamily | 12 | 5,076 | 57% | 90% |
| Calcium channel alpha-1 subunit (TC 1,A,1,11) family | 12 | 12,775 | 53% | 85% |
| CK1 Ser/Thr protein kinase family | 12 | 3,458 | 51% | 80% |
| NR2 subfamily | 12 | 2,759 | 49% | 80% |
| Globin family | 12 | 880 | 49% | 75% |
| TFS-II family | 12 | 1,194 | 48% | 80% |
| Neurexin family | 12 | 5,862 | 40% | 62% |
| KRTAP type 4 family | 12 | 409 | 18% | 70% |
| HERV class-II K(HML-2) subfamily | 12 | 106 | 6% | 27% |
| Peptidase A2 family | 12 | 106 | 6% | 27% |
| Type II cytokine receptor family | 11 | 3,732 | 74% | 100% |
| Metallo-beta-lactamase superfamily | 11 | 3,532 | 74% | 100% |
| IL-1 family | 11 | 1,615 | 74% | 100% |
| Acyl-CoA dehydrogenase family | 11 | 4,459 | 73% | 100% |
| SLC26A/SulP transporter (TC 2,A,53) family | 11 | 5,872 | 73% | 100% |
| Membrane-bound acyltransferase family | 11 | 3,943 | 73% | 99% |
| Complex I LYR family | 11 | 856 | 72% | 100% |
| Protein disulfide isomerase family | 11 | 4,204 | 72% | 100% |
| Peptidase T1B family | 11 | 1,949 | 71% | 100% |
| Non-receptor class subfamily | 11 | 7,198 | 71% | 99% |
| Shisa family | 11 | 2,379 | 70% | 99% |
| Class-I pyridoxal-phosphate-dependent aminotransferase family | 11 | 3,595 | 70% | 98% |
| NEK Ser/Thr protein kinase family | 11 | 5,459 | 69% | 97% |
| NIMA subfamily | 11 | 5,459 | 69% | 97% |
| Toll-like receptor family | 11 | 6,615 | 69% | 96% |
| Conjugate transporter (TC 3,A,1,208) subfamily | 11 | 10,914 | 67% | 97% |
| ADIPOR family | 11 | 2,387 | 66% | 95% |

| | | | | |
|---|---|---|---|---|
| DOCK family | 11 | 14,620 | 66% | 96% |
| ABCA family | 11 | 14,144 | 66% | 97% |
| Synaptobrevin family | 11 | 1,263 | 65% | 94% |
| Plexin family | 11 | 11,706 | 65% | 96% |
| ABCB family | 11 | 6,779 | 65% | 94% |
| E2F/DP family | 11 | 3,454 | 64% | 93% |
| Importin beta family | 11 | 6,978 | 63% | 93% |
| Histone deacetylase family | 11 | 5,060 | 62% | 92% |
| Adaptor complexes large subunit family | 11 | 6,193 | 58% | 86% |
| G-protein coupled receptor Fz/Smo family | 11 | 3,896 | 56% | 89% |
| Secretoglobin family | 11 | 573 | 56% | 82% |
| Histone H1/H5 family | 11 | 1,397 | 55% | 88% |
| Fibrillar collagen family | 11 | 9,840 | 55% | 96% |
| GB1 subfamily | 11 | 3,384 | 51% | 86% |
| GB1/RHD3 GTPase family | 11 | 3,384 | 51% | 86% |
| Recoverin family | 11 | 1,101 | 47% | 80% |
| Opsin subfamily | 11 | 1,865 | 46% | 66% |
| Heat shock protein 90 family | 11 | 2,187 | 37% | 71% |
| Cornifin (SPRR) family | 11 | 273 | 29% | 61% |
| KRTAP type 5 family | 11 | 347 | 16% | 72% |
| HERV class-II K(HML-2) env subfamily | 11 | 297 | 4% | 30% |
| Tubulin--tyrosine ligase family | 10 | 5,743 | 73% | 99% |
| Cation diffusion facilitator (CDF) transporter (TC 2,A,4) family | 10 | 3,438 | 73% | 100% |
| SLC30A subfamily | 10 | 3,438 | 73% | 100% |
| Interleukin-1 receptor family | 10 | 4,039 | 72% | 100% |
| Peptidase S8 family | 10 | 6,827 | 72% | 99% |
| Nectin family | 10 | 3,274 | 71% | 99% |
| XK family | 10 | 3,436 | 71% | 100% |
| Anoctamin family | 10 | 6,363 | 68% | 97% |
| Arrestin family | 10 | 2,696 | 67% | 98% |
| Cullin family | 10 | 6,753 | 67% | 94% |
| TPT transporter family | 10 | 2,307 | 64% | 89% |
| Eukaryotic diacylglycerol kinase family | 10 | 5,955 | 62% | 92% |
| NDK family | 10 | 1,543 | 62% | 89% |
| PMG family | 10 | 1,078 | 62% | 89% |
| Insulin family | 10 | 991 | 61% | 89% |
| TRAPP small subunits family | 10 | 991 | 60% | 83% |
| DNA mismatch repair MutL/HexB family | 10 | 3,273 | 59% | 87% |
| Anion exchanger (TC 2,A,31) family | 10 | 6,330 | 58% | 91% |
| S6 kinase subfamily | 10 | 3,927 | 55% | 90% |
| SRC subfamily | 10 | 2,761 | 53% | 91% |
| Peptidase A1 family | 10 | 2,164 | 51% | 73% |

| | | | | |
|---|---|---|---|---|
| Phosphoglycerate mutase family | 10 | 1,784 | 50% | 81% |
| Sodium channel (TC 1,A,1,10) family | 10 | 8,226 | 43% | 81% |
| KRTAP type 28 family | 10 | 105 | 11% | 60% |
| Beta type-B retroviral Gag protein family | 10 | 481 | 7% | 22% |
| HERV class-II K(HML-2) gag subfamily | 10 | 481 | 7% | 22% |
| Beta type-B retroviral polymerase family | 10 | 429 | 4% | 15% |
| HERV class-II K(HML-2) pol subfamily | 10 | 429 | 4% | 15% |
| BPI/LBP family | 9 | 3,334 | 75% | 100% |
| SMC family | 9 | 8,606 | 74% | 100% |
| Enoyl-CoA hydratase/isomerase family | 9 | 2,157 | 73% | 100% |
| Nucleotide-sugar transporter family | 9 | 2,405 | 73% | 99% |
| Tigger transposable element derived protein family | 9 | 3,572 | 73% | 99% |
| Adenylate kinase family | 9 | 3,056 | 73% | 100% |
| PDGF/VEGF growth factor family | 9 | 1,879 | 72% | 100% |
| Thiolase-like superfamily | 9 | 2,983 | 72% | 99% |
| PNMA family | 9 | 2,611 | 71% | 100% |
| Transglutaminase family | 9 | 4,548 | 71% | 100% |
| Transglutaminase superfamily | 9 | 4,548 | 71% | 100% |
| Chemokine-like factor family | 9 | 1,228 | 71% | 100% |
| Protein arginine N-methyltransferase family | 9 | 3,445 | 71% | 97% |
| Small heat shock protein (HSP20) family | 9 | 1,103 | 70% | 100% |
| IRF family | 9 | 2,692 | 70% | 99% |
| Amiloride-sensitive sodium channel (TC 1,A,6) family | 9 | 3,751 | 70% | 98% |
| Glycosyltransferase 8 family | 9 | 4,066 | 69% | 97% |
| Beta-catenin family | 9 | 5,632 | 68% | 97% |
| Band 7/mec-2 family | 9 | 2,170 | 67% | 96% |
| Fatty acid desaturase type 1 family | 9 | 2,334 | 67% | 97% |
| RsmB/NOP family | 9 | 2,868 | 65% | 92% |
| Fibril-associated collagens with interrupted helices (FACIT) family | 9 | 8,120 | 65% | 100% |
| EMP24/GP25L family | 9 | 1,313 | 65% | 95% |
| NR3 subfamily | 9 | 3,855 | 64% | 93% |
| CACNG subfamily | 9 | 1,649 | 63% | 94% |
| Peptidase M28 family | 9 | 4,024 | 63% | 88% |
| Adrenergic receptor subfamily | 9 | 2,642 | 62% | 97% |
| Ca(2+):cation antiporter (CaCA) (TC 2,A,19) family | 9 | 4,179 | 60% | 90% |
| Sodium/anion cotransporter family | 9 | 2,689 | 60% | 90% |
| Group II decarboxylase family | 9 | 3,054 | 59% | 85% |
| Mas subfamily | 9 | 1,720 | 59% | 92% |
| SLC12A transporter family | 9 | 5,333 | 57% | 89% |
| Copine family | 9 | 2,849 | 56% | 93% |
| Chloride channel (TC 2,A,49) family | 9 | 4,035 | 55% | 84% |

| | | | | |
|---|---|---|---|---|
| Adaptor complexes small subunit family | 9 | 827 | 54% | 85% |
| Elastase subfamily | 9 | 1,169 | 49% | 79% |
| TFA subfamily | 9 | 631 | 41% | 73% |
| Gamma-glutamyltransferase family | 9 | 1,478 | 36% | 60% |
| SPAN-X family | 9 | 305 | 32% | 72% |
| Centaurin gamma-like family | 9 | 1,972 | 29% | 61% |
| SSX family | 9 | 478 | 28% | 72% |
| Histone H3 family | 9 | 200 | 16% | 33% |
| POTE family | 9 | 439 | 9% | 31% |
| CT45 family | 9 | 32 | 2% | 7% |
| Type 1 subfamily | 8 | 3,117 | 75% | 100% |
| Class IV-like SAM-binding methyltransferase superfamily | 8 | 3,043 | 75% | 100% |
| DNA2/NAM7 helicase family | 8 | 9,439 | 75% | 100% |
| Quinone oxidoreductase subfamily | 8 | 2,200 | 74% | 100% |
| AlkB family | 8 | 2,042 | 74% | 100% |
| Polycystin family | 8 | 11,839 | 74% | 100% |
| Patched family | 8 | 6,856 | 73% | 100% |
| Ubiquitin-activating E1 family | 8 | 4,007 | 73% | 100% |
| MCM family | 8 | 4,957 | 73% | 100% |
| STXBP/unc-18/SEC1 family | 8 | 3,535 | 72% | 100% |
| FAM83 family | 8 | 4,145 | 72% | 100% |
| TMC family | 8 | 4,841 | 72% | 99% |
| IL-6 superfamily | 8 | 1,213 | 72% | 100% |
| Villin/gelsolin family | 8 | 4,980 | 71% | 100% |
| Integrin beta chain family | 8 | 5,226 | 71% | 99% |
| Diacylglycerol acyltransferase family | 8 | 1,936 | 71% | 100% |
| GARIN family | 8 | 2,603 | 71% | 99% |
| Peptidase S54 family | 8 | 2,734 | 70% | 98% |
| AIG1/Toc34/Toc159-like paraseptin GTPase family | 8 | 1,921 | 70% | 100% |
| IAN subfamily | 8 | 1,921 | 70% | 100% |
| GDA1/CD39 NTPase family | 8 | 2,851 | 69% | 98% |
| CSF-1/PDGF receptor subfamily | 8 | 6,278 | 68% | 98% |
| FXYD family | 8 | 532 | 68% | 99% |
| DMRT family | 8 | 2,252 | 67% | 97% |
| LTrpC subfamily | 8 | 8,071 | 66% | 95% |
| Chondroitin N-acetylgalactosaminyltransferase family | 8 | 4,191 | 66% | 97% |
| RFX family | 8 | 4,397 | 66% | 94% |
| Peptidase T1A family | 8 | 1,280 | 64% | 93% |
| CD300 family | 8 | 1,242 | 64% | 95% |
| Ephrin family | 8 | 1,340 | 64% | 96% |
| Histidine acid phosphatase family | 8 | 3,373 | 64% | 91% |

| | | | | |
|---|---|---|---|---|
| Adaptor complexes medium subunit family | 8 | 2,269 | 64% | 93% |
| LDH/MDH superfamily | 8 | 1,779 | 61% | 95% |
| Tenascin family | 8 | 12,064 | 60% | 96% |
| Actin-binding proteins ADF family | 8 | 971 | 60% | 91% |
| Glutathione peroxidase family | 8 | 990 | 60% | 95% |
| HSF family | 8 | 2,152 | 59% | 81% |
| Beta-type (group I) subfamily | 8 | 1,182 | 59% | 92% |
| Peroxidase family | 8 | 5,314 | 58% | 85% |
| NOTCH family | 8 | 6,018 | 58% | 88% |
| MAP kinase kinase subfamily | 8 | 1,755 | 57% | 87% |
| Glycosyltransferase 10 family | 8 | 1,860 | 56% | 83% |
| Glycosyl hydrolase 22 family | 8 | 697 | 56% | 82% |
| H (Eag) (TC 1,A,1,20) subfamily | 8 | 4,636 | 54% | 86% |
| Argonaute family | 8 | 3,751 | 54% | 85% |
| Dwarfin/SMAD family | 8 | 1,699 | 45% | 74% |
| A (Shaker) (TC 1,A,1,2) subfamily | 8 | 1,923 | 44% | 82% |
| G(i/o/t/z) subfamily | 8 | 1,032 | 36% | 74% |
| ATG8 family | 8 | 344 | 35% | 65% |
| KRTAP type 19 family | 8 | 189 | 34% | 87% |
| Glycoprotein hormones subunit beta family | 8 | 388 | 32% | 50% |
| Ubiquitin family | 8 | 209 | 14% | 26% |
| KRTAP type 9 family | 8 | 182 | 13% | 60% |
| PPIase A subfamily | 8 | 100 | 8% | 27% |
| Peptidase M16 family | 7 | 3,852 | 75% | 100% |
| Type 5 subfamily | 7 | 2,047 | 75% | 100% |
| NAD(P)-dependent epimerase/dehydratase family | 7 | 1,726 | 74% | 100% |
| TRAFAC class YlqF/YawG GTPase family | 7 | 3,067 | 74% | 100% |
| Glycerophosphoryl diester phosphodiesterase family | 7 | 2,499 | 73% | 100% |
| Exportin family | 7 | 5,620 | 73% | 99% |
| Bile acid:sodium symporter (BASS) (TC 2,A,28) family | 7 | 2,013 | 73% | 100% |
| Glyoxalase II family | 7 | 1,506 | 73% | 100% |
| Sirtuin family | 7 | 2,118 | 73% | 100% |
| Peptidase C48 family | 7 | 3,583 | 73% | 99% |
| RecA family | 7 | 1,731 | 72% | 100% |
| TRAFAC class OBG-HflX-like GTPase superfamily | 7 | 2,207 | 72% | 99% |
| P2X receptor family | 7 | 2,226 | 72% | 100% |
| Nucleotide pyrophosphatase/phosphodiesterase family | 7 | 3,199 | 71% | 99% |
| Cytochrome b5 family | 7 | 967 | 71% | 99% |
| Glycosyl hydrolase 18 family | 7 | 2,244 | 71% | 100% |
| ELO family | 7 | 1,415 | 71% | 100% |
| Sulfotransferase 2 family | 7 | 1,903 | 70% | 98% |

| | | | | |
|---|---|---|---|---|
| DNA polymerase type-B-like family | 7 | 4,529 | 70% | 98% |
| Carnitine/choline acetyltransferase family | 7 | 3,509 | 70% | 98% |
| Glycosyltransferase 14 family | 7 | 2,787 | 70% | 98% |
| Flavin monoamine oxidase family | 7 | 3,050 | 70% | 98% |
| Apolipoprotein L family | 7 | 1,779 | 70% | 99% |
| Glycosyltransferase group 1 family | 7 | 2,004 | 70% | 96% |
| Glycosyl hydrolase 47 family | 7 | 3,306 | 69% | 98% |
| YIP1 family | 7 | 1,365 | 69% | 98% |
| 5'(3')-deoxyribonucleotidase family | 7 | 2,016 | 69% | 98% |
| Inositol phosphokinase (IPK) family | 7 | 2,574 | 68% | 98% |
| Spectrin family | 7 | 12,236 | 68% | 97% |
| Glycosyltransferase 7 family | 7 | 1,749 | 67% | 96% |
| Thrombospondin family | 7 | 6,993 | 67% | 97% |
| Tropomodulin family | 7 | 2,065 | 66% | 99% |
| FKBP-type PPIase family | 7 | 1,395 | 66% | 93% |
| CCR4/nocturin family | 7 | 2,454 | 66% | 95% |
| LN-TM7 subfamily | 7 | 9,046 | 66% | 96% |
| Alpha-type (group II) subfamily | 7 | 1,943 | 65% | 97% |
| Gal/GlcNAc/GalNAc subfamily | 7 | 1,934 | 62% | 91% |
| Dicarboxylate/amino acid:cation symporter (DAACS) (TC 2,A,23) family | 7 | 2,382 | 62% | 94% |
| TALE/IRO homeobox family | 7 | 2,005 | 62% | 93% |
| Glycosyl hydrolase 31 family | 7 | 6,480 | 61% | 96% |
| Peptidase M13 family | 7 | 3,274 | 61% | 85% |
| ClpA/ClpB family | 7 | 1,664 | 60% | 85% |
| M28B subfamily | 7 | 3,005 | 60% | 85% |
| Distal-less homeobox family | 7 | 1,067 | 59% | 91% |
| Schlafen family | 7 | 3,013 | 58% | 87% |
| Transcription factor STAT family | 7 | 3,143 | 57% | 81% |
| Plakin or cytolinker family | 7 | 13,504 | 56% | 87% |
| NK-2 homeobox family | 7 | 1,245 | 56% | 90% |
| CUT homeobox family | 7 | 3,258 | 55% | 83% |
| GPRK subfamily | 7 | 2,280 | 54% | 84% |
| SCC3 family | 7 | 2,282 | 53% | 80% |
| L-type amino acid transporter (LAT) (TC 2,A,3,8) family | 7 | 1,545 | 53% | 88% |
| FGGY kinase family | 7 | 1,866 | 50% | 72% |
| MOB1/phocein family | 7 | 761 | 49% | 77% |
| WD repeat Groucho/TLE family | 7 | 2,014 | 47% | 78% |
| Importin alpha family | 7 | 1,695 | 46% | 78% |
| 14-3-3 family | 7 | 708 | 41% | 77% |
| TALE/MEIS homeobox family | 7 | 1,004 | 36% | 64% |
| POM121 family | 7 | 1,833 | 31% | 59% |
| Casein kinase I subfamily | 7 | 810 | 29% | 56% |

| | | | | |
|---|---|---|---|---|
| RALY subfamily | 7 | 517 | 25% | 49% |
| RRM HNRPC family | 7 | 517 | 25% | 49% |
| UPF0607 family | 7 | 421 | 19% | 66% |
| NUT family | 7 | 948 | 16% | 33% |
| SSU72 phosphatase family | 7 | 193 | 14% | 45% |
| CTAGE family | 7 | 374 | 8% | 18% |
| Peptidase M67A family | 6 | 1,823 | 75% | 100% |
| PLAC1 family | 6 | 767 | 74% | 100% |
| Type 4 subfamily | 6 | 2,805 | 74% | 100% |
| Gasdermin family | 6 | 1,991 | 74% | 100% |
| Sterol desaturase family | 6 | 1,483 | 74% | 100% |
| Complement C6/C7/C8/C9 family | 6 | 2,993 | 74% | 100% |
| Prefoldin subunit beta family | 6 | 1,071 | 74% | 100% |
| Thiolase family | 6 | 1,962 | 74% | 100% |
| Dynein light chain Tctex-type family | 6 | 711 | 73% | 100% |
| CALHM family | 6 | 1,426 | 73% | 100% |
| ITIH family | 6 | 4,325 | 73% | 100% |
| ELL/occludin family | 6 | 2,180 | 73% | 100% |
| 'GDXG' lipolytic enzyme family | 6 | 2,249 | 73% | 99% |
| RAD51 subfamily | 6 | 1,484 | 72% | 100% |
| BI1 family | 6 | 1,311 | 72% | 100% |
| IL-10 family | 6 | 783 | 72% | 100% |
| Peptidase S9B family | 6 | 3,561 | 72% | 99% |
| SLC35F solute transporter family | 6 | 1,884 | 72% | 99% |
| OBG GTPase family | 6 | 1,836 | 72% | 99% |
| MAPEG family | 6 | 658 | 72% | 100% |
| RNase PH family | 6 | 1,261 | 72% | 100% |
| Small Tim family | 6 | 399 | 72% | 99% |
| Tektin family | 6 | 1,703 | 71% | 99% |
| Ferlin family | 6 | 8,460 | 71% | 99% |
| CCN family | 6 | 1,466 | 71% | 100% |
| Fibulin family | 6 | 2,639 | 71% | 100% |
| Phospholipase D family | 6 | 2,681 | 71% | 99% |
| MAL family | 6 | 914 | 71% | 100% |
| Alpha-type protein kinase family | 6 | 7,021 | 71% | 98% |
| Eutherian X-chromosome-specific Armcx family | 6 | 3,246 | 70% | 100% |
| Carbon-nitrogen hydrolase superfamily | 6 | 1,802 | 70% | 99% |
| IL-17 family | 6 | 755 | 70% | 100% |
| CNC subfamily | 6 | 2,823 | 70% | 99% |
| Glucagon family | 6 | 637 | 70% | 100% |
| Cyclin AB subfamily | 6 | 2,732 | 70% | 99% |
| L6 tetraspanin family | 6 | 866 | 70% | 100% |

| | | | | |
|---|---|---|---|---|
| DOK family | 6 | 1,641 | 70% | 98% |
| Lipoxygenase family | 6 | 2,830 | 69% | 98% |
| SLITRK family | 6 | 3,553 | 69% | 99% |
| H-rev107 family | 6 | 799 | 69% | 98% |
| Zyxin/ajuba family | 6 | 2,258 | 68% | 98% |
| Glypican family | 6 | 2,323 | 68% | 97% |
| Adenosine and AMP deaminases family | 6 | 2,465 | 68% | 97% |
| NIPA family | 6 | 1,546 | 67% | 98% |
| Contactin family | 6 | 4,170 | 67% | 98% |
| Class-I pyridine nucleotide-disulfide oxidoreductase family | 6 | 2,235 | 67% | 96% |
| XPO subfamily | 6 | 4,026 | 67% | 96% |
| HD type 2 subfamily | 6 | 3,922 | 65% | 96% |
| Cyclic nucleotide-gated cation channel (TC 1,A,1,5) family | 6 | 3,016 | 64% | 96% |
| SLC24A subfamily | 6 | 2,634 | 64% | 91% |
| Class-II pyridoxal-phosphate-dependent aminotransferase family | 6 | 2,067 | 64% | 93% |
| FMO family | 6 | 2,044 | 63% | 96% |
| TrpV subfamily | 6 | 2,991 | 63% | 92% |
| WD repeat EMAP family | 6 | 4,548 | 63% | 96% |
| PtdIns transfer protein family | 6 | 2,772 | 62% | 95% |
| BTG family | 6 | 929 | 62% | 92% |
| Granzyme subfamily | 6 | 946 | 62% | 94% |
| Peroxiredoxin family | 6 | 844 | 62% | 95% |
| WD repeat coronin family | 6 | 2,077 | 62% | 95% |
| Protease inhibitor I39 (alpha-2-macroglobulin) family | 6 | 5,486 | 61% | 89% |
| SEC23/SEC24 family | 6 | 3,676 | 61% | 91% |
| DP1 family | 6 | 825 | 60% | 95% |
| IAP family | 6 | 1,434 | 60% | 90% |
| LHFP family | 6 | 808 | 60% | 94% |
| Chloride channel CLIC family | 6 | 1,245 | 60% | 95% |
| NPY family | 6 | 244 | 59% | 96% |
| Calponin family | 6 | 904 | 59% | 94% |
| 3-beta-HSD family | 6 | 1,353 | 59% | 84% |
| Dpy-19 family | 6 | 1,970 | 56% | 86% |
| Grh/CP2 family | 6 | 1,908 | 56% | 89% |
| Torsin subfamily | 6 | 1,149 | 56% | 80% |
| SIX/Sine oculis homeobox family | 6 | 1,485 | 56% | 85% |
| Type IV collagen family | 6 | 5,607 | 55% | 98% |
| Liprin family | 6 | 3,679 | 55% | 86% |
| JHDM3 histone demethylase family | 6 | 2,642 | 54% | 84% |
| Hydantoinase/dihydropyrimidinase family | 6 | 1,812 | 54% | 90% |
| Maf subfamily | 6 | 820 | 54% | 91% |

| | | | | |
|---|---|---|---|---|
| STrpC subfamily | 6 | 2,845 | 52% | 84% |
| Tim17/Tim22/Tim23 family | 6 | 628 | 52% | 77% |
| CELF/BRUNOL family | 6 | 1,380 | 47% | 82% |
| C/M/P thioester hydrolase family | 6 | 1,114 | 45% | 68% |
| WD repeat G protein beta family | 6 | 916 | 44% | 78% |
| Type IIC subfamily | 6 | 2,515 | 41% | 78% |
| Calmodulin family | 6 | 346 | 36% | 52% |
| PRR23 family | 6 | 443 | 28% | 52% |
| Somatotropin/prolactin family | 6 | 332 | 25% | 48% |
| Thymosin beta family | 6 | 52 | 19% | 40% |
| Glycosyl hydrolase 13 family | 6 | 580 | 18% | 30% |
| SIMIBI class G3E GTPase family | 6 | 376 | 16% | 35% |
| VCX/VCY family | 6 | 144 | 14% | 49% |
| RRM DAZ family | 6 | 365 | 12% | 26% |
| PRR20 family | 6 | 150 | 11% | 16% |
| MBD3L family | 6 | 139 | 11% | 28% |
| Class-V pyridoxal-phosphate-dependent aminotransferase family | 5 | 1,931 | 76% | 100% |
| Glycosyltransferase 4 subfamily | 5 | 1,652 | 75% | 100% |
| Metallophosphoesterase superfamily | 5 | 1,438 | 75% | 100% |
| FAST kinase family | 5 | 2,689 | 74% | 100% |
| LAMP family | 5 | 1,396 | 74% | 100% |
| DNA mismatch repair MutS family | 5 | 3,867 | 74% | 100% |
| Peptidase M20A family | 5 | 1,725 | 74% | 100% |
| GcvT family | 5 | 2,525 | 74% | 100% |
| Plunc family | 5 | 1,049 | 74% | 100% |
| Steroid 5-alpha reductase family | 5 | 1,107 | 74% | 100% |
| SEC6 family | 5 | 2,409 | 74% | 100% |
| Glycosyltransferase 47 family | 5 | 2,489 | 73% | 100% |
| Peroxisomal membrane protein PXMP2/4 family | 5 | 722 | 73% | 100% |
| SKI2 subfamily | 5 | 5,092 | 73% | 100% |
| TNF receptor-associated factor family | 5 | 1,914 | 73% | 100% |
| Activator 1 small subunits family | 5 | 1,745 | 73% | 100% |
| DRAM/TMEM150 family | 5 | 918 | 73% | 100% |
| Glycosyl hydrolase 1 family | 5 | 3,661 | 73% | 100% |
| ICAM family | 5 | 1,857 | 73% | 100% |
| SelWTH family | 5 | 630 | 73% | 99% |
| X(+)/potassium ATPases subunit beta family | 5 | 1,107 | 73% | 100% |
| TUBGCP family | 5 | 3,854 | 72% | 100% |
| Glycosyltransferase 6 family | 5 | 1,024 | 72% | 100% |
| RecQ subfamily | 5 | 4,115 | 72% | 99% |
| DIPK family | 5 | 1,545 | 72% | 100% |
| METTL21 family | 5 | 870 | 72% | 100% |

| | | | | |
|---|---|---|---|---|
| DCC family | 5 | 4,409 | 72% | 100% |
| Glycosyl hydrolase 56 family | 5 | 1,667 | 72% | 100% |
| Type 3 subfamily | 5 | 1,377 | 72% | 100% |
| Inositol monophosphatase superfamily | 5 | 1,174 | 72% | 100% |
| Multicopper oxidase family | 5 | 5,725 | 72% | 100% |
| ADCK protein kinase family | 5 | 2,105 | 72% | 99% |
| Flavoprotein pyridine nucleotide cytochrome reductase family | 5 | 1,229 | 72% | 99% |
| Glycosyl hydrolase 38 family | 5 | 3,827 | 71% | 99% |
| GDNFR family | 5 | 1,445 | 71% | 99% |
| Class-III pyridoxal-phosphate-dependent aminotransferase family | 5 | 1,714 | 71% | 99% |
| Peptidase A22B family | 5 | 1,824 | 71% | 100% |
| CTL (choline transporter-like) family | 5 | 2,451 | 71% | 98% |
| SLC35A subfamily | 5 | 1,284 | 71% | 99% |
| Calcium channel subunit alpha-2/delta family | 5 | 4,084 | 71% | 99% |
| CDC5/Polo subfamily | 5 | 2,294 | 71% | 99% |
| MINDY deubiquitinase family | 5 | 1,945 | 71% | 99% |
| BET3 subfamily | 5 | 612 | 71% | 99% |
| SLRP class II subfamily | 5 | 1,316 | 70% | 100% |
| CREC family | 5 | 1,159 | 70% | 99% |
| Phosphohexose mutase family | 5 | 2,034 | 70% | 98% |
| DAN family | 5 | 692 | 70% | 98% |
| VPS10-related sortilin family | 5 | 4,612 | 70% | 97% |
| PTEN phosphatase protein family | 5 | 4,060 | 70% | 98% |
| Type VI collagen family | 5 | 7,057 | 70% | 99% |
| Proton-dependent oligopeptide transporter (POT/PTR) (TC 2,A,17) family | 5 | 2,217 | 70% | 98% |
| Leprecan family | 5 | 2,106 | 70% | 99% |
| TUB family | 5 | 2,467 | 69% | 98% |
| NADC subfamily | 5 | 2,066 | 69% | 99% |
| SLC13A/DASS transporter (TC 2,A,47) family | 5 | 2,066 | 69% | 99% |
| TRX/MLL subfamily | 5 | 13,133 | 69% | 98% |
| Dynein light intermediate chain family | 5 | 2,050 | 69% | 98% |
| CAF1 family | 5 | 1,549 | 69% | 96% |
| CDP-alcohol phosphatidyltransferase class-I family | 5 | 1,185 | 68% | 97% |
| Secreted frizzled-related protein (sFRP) family | 5 | 1,091 | 68% | 97% |
| Type V subfamily | 5 | 4,114 | 68% | 98% |
| MyBP family | 5 | 2,987 | 68% | 99% |
| TDE1 family | 5 | 1,579 | 68% | 99% |
| Syntrophin family | 5 | 1,794 | 68% | 98% |
| Vasopressin/oxytocin receptor subfamily | 5 | 1,338 | 68% | 99% |
| C/EBP subfamily | 5 | 949 | 68% | 98% |
| BORG/CEP family | 5 | 919 | 68% | 99% |

| | | | | |
|---|---|---|---|---|
| Uridine kinase family | 5 | 1,019 | 67% | 96% |
| TMEM132 family | 5 | 3,602 | 67% | 96% |
| JHDM1 histone demethylase family | 5 | 3,738 | 67% | 97% |
| Deltex family | 5 | 1,969 | 67% | 98% |
| ABCG family | 5 | 2,204 | 67% | 94% |
| Eye pigment precursor importer (TC 3,A,1,204) subfamily | 5 | 2,204 | 67% | 94% |
| SCAMP family | 5 | 986 | 67% | 99% |
| Isocitrate and isopropylmalate dehydrogenases family | 5 | 1,339 | 67% | 95% |
| Unc-5 family | 5 | 2,776 | 66% | 98% |
| Sideroflexin family | 5 | 1,085 | 66% | 93% |
| Prickle / espinas / testin family | 5 | 2,018 | 66% | 97% |
| Cyclin C subfamily | 5 | 1,742 | 66% | 95% |
| Lipophilin subfamily | 5 | 297 | 66% | 97% |
| SAPAP family | 5 | 3,186 | 66% | 97% |
| Protein arginine deiminase family | 5 | 2,182 | 65% | 97% |
| Cationic amino acid transporter (CAT) (TC 2,A,3,3) family | 5 | 2,157 | 65% | 96% |
| LRRC8 family | 5 | 2,642 | 65% | 96% |
| TEC subfamily | 5 | 2,020 | 65% | 99% |
| Lysyl oxidase family | 5 | 2,115 | 65% | 97% |
| KHDC1 family | 5 | 545 | 64% | 96% |
| Ferritin family | 5 | 633 | 64% | 97% |
| HDGF family | 5 | 1,216 | 64% | 95% |
| Calcineurin regulatory subunit family | 5 | 603 | 64% | 95% |
| EGR C2H2-type zinc-finger protein family | 5 | 1,555 | 64% | 96% |
| MYST (SAS/MOZ) family | 5 | 3,589 | 63% | 94% |
| LRFN family | 5 | 2,243 | 63% | 98% |
| DAP kinase subfamily | 5 | 1,925 | 63% | 95% |
| Phospholipid scramblase family | 5 | 951 | 63% | 97% |
| Glycine N-acyltransferase family | 5 | 924 | 62% | 92% |
| IFIT family | 5 | 1,488 | 62% | 95% |
| ERF2/ZDHHC9 subfamily | 5 | 1,678 | 62% | 94% |
| Perilipin family | 5 | 1,987 | 62% | 100% |
| KQT (TC 1,A,1,15) subfamily | 5 | 2,491 | 62% | 93% |
| PDK/BCKDK protein kinase family | 5 | 1,275 | 62% | 96% |
| Multidrug resistance exporter (TC 3,A,1,201) subfamily | 5 | 3,614 | 61% | 92% |
| ING family | 5 | 989 | 61% | 92% |
| 5-hydroxytryptamine receptor (TC 1,A,9,2) subfamily | 5 | 1,394 | 61% | 92% |
| Endophilin family | 5 | 1,100 | 60% | 91% |
| TCP11 family | 5 | 1,353 | 60% | 85% |
| ATPase alpha/beta chains family | 5 | 1,630 | 60% | 84% |

| | | | | |
|---|---|---|---|---|
| Vinculin/alpha-catenin family | 5 | 2,763 | 60% | 89% |
| IgLON family | 5 | 1,026 | 60% | 89% |
| HMGN family | 5 | 391 | 59% | 94% |
| Unc-13 family | 5 | 4,597 | 59% | 88% |
| STEAP family | 5 | 1,190 | 59% | 86% |
| MNB/DYRK subfamily | 5 | 1,824 | 59% | 90% |
| Muscarinic acetylcholine receptor subfamily | 5 | 1,481 | 59% | 92% |
| Potassium channel KCNE family | 5 | 420 | 58% | 85% |
| Hexokinase family | 5 | 2,421 | 58% | 94% |
| Ikaros C2H2-type zinc-finger protein family | 5 | 1,479 | 58% | 89% |
| TAFA family | 5 | 384 | 57% | 93% |
| Mab-21 family | 5 | 1,191 | 57% | 81% |
| SLC35E subfamily | 5 | 989 | 57% | 78% |
| Snail C2H2-type zinc-finger protein family | 5 | 809 | 55% | 87% |
| Folate receptor family | 5 | 662 | 54% | 85% |
| KRTAP type 20 family | 5 | 136 | 54% | 100% |
| AP-2 family | 5 | 1,176 | 52% | 85% |
| Synaptogyrin family | 5 | 591 | 52% | 80% |
| Ammonium transporter (TC 2,A,49) family | 5 | 1,126 | 52% | 82% |
| Rh subfamily | 5 | 1,126 | 52% | 82% |
| LDH family | 5 | 882 | 52% | 91% |
| Type IIA subfamily | 5 | 2,411 | 49% | 80% |
| Phosphatase 2A regulatory subunit B56 family | 5 | 1,185 | 46% | 78% |
| ANP32 family | 5 | 519 | 46% | 88% |
| BEX family | 5 | 270 | 45% | 72% |
| BPG-dependent PGAM subfamily | 5 | 594 | 45% | 71% |
| FAM47 family | 5 | 1,465 | 45% | 87% |
| TMEM14 family | 5 | 244 | 43% | 72% |
| Alpha-defensin family | 5 | 202 | 42% | 64% |
| HMGB family | 5 | 402 | 39% | 80% |
| Polyadenylate-binding protein type-1 family | 5 | 1,093 | 38% | 74% |
| Plasminogen subfamily | 5 | 1,805 | 36% | 79% |
| Yippee family | 5 | 218 | 36% | 66% |
| SLC35G solute transporter family | 5 | 604 | 34% | 72% |
| Mu family | 5 | 368 | 34% | 63% |
| Ribose-phosphate pyrophosphokinase family | 5 | 553 | 33% | 65% |
| Alpha family | 5 | 359 | 32% | 65% |
| SUMO subfamily | 5 | 156 | 32% | 59% |
| cAMP subfamily | 5 | 498 | 30% | 54% |
| SPIN/STSY family | 5 | 349 | 27% | 51% |
| RIMBP family | 5 | 2,091 | 27% | 40% |
| Tryptase subfamily | 5 | 343 | 25% | 48% |

| | | | | |
|---|---|---|---|---|
| EEF2KMT family | 5 | 219 | 17% | 40% |
| PP-1 subfamily | 5 | 222 | 14% | 27% |
| DIPP subfamily | 5 | 113 | 13% | 26% |
| BAGE family | 5 | 26 | 8% | 25% |
| WASH1 family | 5 | 167 | 7% | 31% |
| ZNG1 subfamily | 5 | 58 | 3% | 21% |
| Eukaryotic ribosomal protein eL8 family | 4 | 631 | 76% | 100% |
| FAD-binding oxidoreductase/transferase type 4 family | 4 | 1,671 | 76% | 100% |
| DNA polymerase type-B family | 4 | 6,046 | 76% | 100% |
| SLC35B subfamily | 4 | 1,121 | 75% | 100% |
| Arg-specific ADP-ribosyltransferase family | 4 | 995 | 75% | 100% |
| Thioredoxin family | 4 | 390 | 75% | 100% |
| Nonaspanin (TM9SF) (TC 9,A,2) family | 4 | 1,867 | 75% | 100% |
| EIF-2B alpha/beta/delta subunits family | 4 | 1,156 | 75% | 100% |
| Peptidase C65 family | 4 | 905 | 75% | 100% |
| Izumo family | 4 | 777 | 75% | 100% |
| PIH1 family | 4 | 1,233 | 74% | 100% |
| GCN2 subfamily | 4 | 2,937 | 74% | 100% |
| DNA polymerase type-Y family | 4 | 2,660 | 74% | 100% |
| ALB/AFP/VDB family | 4 | 1,705 | 74% | 100% |
| RNA-metabolizing metallo-beta-lactamase-like family | 4 | 2,026 | 74% | 100% |
| VPS13 family | 4 | 11,407 | 74% | 100% |
| FRAS1 family | 4 | 8,547 | 74% | 100% |
| XPG/RAD2 endonuclease family | 4 | 2,464 | 74% | 100% |
| Acyl-CoA oxidase family | 4 | 1,919 | 74% | 100% |
| Glycosyltransferase 22 family | 4 | 1,654 | 74% | 100% |
| PATE family | 4 | 322 | 74% | 100% |
| Serine esterase family | 4 | 1,102 | 74% | 99% |
| Glycosyl hydrolase 33 family | 4 | 1,261 | 74% | 100% |
| AKAP110 family | 4 | 3,919 | 74% | 100% |
| Pseudouridine synthase RluA family | 4 | 1,170 | 74% | 100% |
| Cation channel sperm-associated (TC 1,A,1,19) family | 4 | 1,609 | 74% | 100% |
| FPP/GGPP synthase family | 4 | 1,131 | 74% | 100% |
| 2-oxoacid dehydrogenase family | 4 | 1,533 | 74% | 100% |
| MTERF family | 4 | 1,162 | 73% | 100% |
| R-spondin family | 4 | 743 | 73% | 100% |
| Dus family | 4 | 1,419 | 73% | 100% |
| STKL subfamily | 4 | 1,275 | 73% | 100% |
| WAL family | 4 | 5,220 | 73% | 100% |
| EF-G/EF-2 subfamily | 4 | 2,466 | 73% | 100% |
| KCNMB (TC 8,A,14,1) family | 4 | 670 | 73% | 100% |

| | | | | |
|---|---|---|---|---|
| Glutaredoxin family | 4 | 413 | 73% | 100% |
| Peptidase M24B family | 4 | 1,679 | 73% | 100% |
| Gamma-glutamylcyclotransferase family | 4 | 546 | 73% | 100% |
| Pelle subfamily | 4 | 1,749 | 73% | 100% |
| Klotho subfamily | 4 | 2,259 | 73% | 100% |
| GHMP kinase family | 4 | 1,701 | 73% | 100% |
| AB hydrolase 4 family | 4 | 1,246 | 73% | 100% |
| DNA polymerase type-X family | 4 | 1,396 | 73% | 100% |
| Reduced folate carrier (RFC) transporter (TC 2,A,48) family | 4 | 1,264 | 73% | 100% |
| Nesprin family | 4 | 12,422 | 73% | 99% |
| SLC29A/ENT transporter (TC 2,A,57) family | 4 | 1,395 | 73% | 100% |
| Mastermind family | 4 | 2,970 | 73% | 100% |
| Myelin P0 protein family | 4 | 703 | 73% | 100% |
| A subfamily | 4 | 1,560 | 73% | 100% |
| Very long-chain fatty acids dehydratase HACD family | 4 | 825 | 73% | 100% |
| Glycoside-pentoside-hexuronide (GPH) cation symporter transporter (TC 2,A,2) family | 4 | 1,911 | 73% | 100% |
| CAVIN family | 4 | 1,045 | 73% | 100% |
| SOWAH family | 4 | 1,583 | 73% | 100% |
| CIMAP family | 4 | 775 | 72% | 100% |
| Piwi subfamily | 4 | 2,580 | 72% | 99% |
| 'GDSL' lipolytic enzyme family | 4 | 1,564 | 72% | 99% |
| TMTC family | 4 | 2,430 | 72% | 99% |
| SET2 subfamily | 4 | 5,996 | 72% | 99% |
| EPS8 family | 4 | 2,048 | 72% | 100% |
| LFG subfamily | 4 | 887 | 72% | 100% |
| L1/neurofascin/NgCAM family | 4 | 3,667 | 72% | 99% |
| C5-methyltransferase family | 4 | 2,703 | 72% | 98% |
| Profilin family | 4 | 391 | 72% | 100% |
| Protease inhibitor I35 (TIMP) family | 4 | 617 | 72% | 99% |
| Galactose-3-O-sulfotransferase family | 4 | 1,244 | 72% | 100% |
| Shroom family | 4 | 4,261 | 72% | 99% |
| Atypical chemokine receptor subfamily | 4 | 1,024 | 72% | 100% |
| Peptidase C12 family | 4 | 1,080 | 71% | 100% |
| ZP domain family | 4 | 1,676 | 71% | 97% |
| PC-esterase family | 4 | 1,934 | 71% | 99% |
| Paralemmin family | 4 | 1,420 | 71% | 99% |
| DPPIV subfamily | 4 | 2,369 | 71% | 99% |
| MAP7 family | 4 | 2,279 | 71% | 100% |
| 17-beta-HSD 3 subfamily | 4 | 892 | 71% | 98% |
| Prokaryotic/mitochondrial release factor family | 4 | 852 | 71% | 100% |
| Gfo/Idh/MocA family | 4 | 1,000 | 71% | 98% |

| | | | | |
|---|---|---|---|---|
| VPS37 family | 4 | 916 | 71% | 100% |
| Transglutaminase-like superfamily | 4 | 1,446 | 71% | 98% |
| CAS family | 4 | 2,167 | 71% | 98% |
| Sauvagine/corticotropin-releasing factor/urotensin I family | 4 | 421 | 71% | 100% |
| Apolipoprotein A1/A4/E family | 4 | 954 | 71% | 100% |
| TPD52 family | 4 | 548 | 71% | 100% |
| Phosducin family | 4 | 727 | 71% | 100% |
| Suvar3-9 subfamily | 4 | 2,351 | 71% | 99% |
| JAK subfamily | 4 | 3,247 | 71% | 99% |
| Neuregulin family | 4 | 1,642 | 71% | 99% |
| Glycosyl hydrolase 2 family | 4 | 1,371 | 71% | 98% |
| Purine/pyrimidine phosphoribosyltransferase family | 4 | 804 | 71% | 99% |
| Receptor class 3 subfamily | 4 | 3,995 | 71% | 100% |
| LIMR family | 4 | 1,558 | 70% | 98% |
| GAB family | 4 | 1,778 | 70% | 99% |
| APG1/unc-51/ULK1 subfamily | 4 | 2,693 | 70% | 98% |
| AF4 family | 4 | 3,445 | 70% | 99% |
| Dickkopf family | 4 | 771 | 70% | 99% |
| WD repeat PROPPIN family | 4 | 1,125 | 70% | 99% |
| Inositol 1,4,5-trisphosphate 5-phosphatase family | 4 | 3,866 | 70% | 98% |
| Peptidase C54 family | 4 | 1,207 | 70% | 99% |
| Canopy family | 4 | 560 | 70% | 99% |
| RASGRP family | 4 | 1,938 | 70% | 98% |
| CLCR family | 4 | 2,124 | 70% | 99% |
| NGF-beta family | 4 | 667 | 70% | 100% |
| Inositol 1,4,5-trisphosphate 5-phosphatase type II family | 4 | 2,338 | 70% | 99% |
| SLRP class I subfamily | 4 | 1,259 | 70% | 99% |
| Syndecan proteoglycan family | 4 | 801 | 70% | 99% |
| GAS2 family | 4 | 1,786 | 70% | 99% |
| FHIP family | 4 | 2,447 | 70% | 99% |
| GDNF subfamily | 4 | 545 | 70% | 100% |
| NF-kappa-B inhibitor family | 4 | 1,032 | 69% | 100% |
| SCM family | 4 | 1,460 | 69% | 99% |
| I-kappa-B kinase subfamily | 4 | 2,043 | 69% | 97% |
| MAPR subfamily | 4 | 592 | 69% | 98% |
| DNase I family | 4 | 823 | 69% | 99% |
| Sarcoglycan beta/delta/gamma/zeta family | 4 | 829 | 69% | 97% |
| Organophosphate:Pi antiporter (OPA) (TC 2,A,1,4) family | 4 | 1,350 | 69% | 97% |
| Furin subfamily | 4 | 2,028 | 69% | 98% |
| LU7TM family | 4 | 1,552 | 69% | 97% |
| Glycosyltransferase 54 family | 4 | 1,329 | 69% | 95% |

| | | | | |
|---|---|---|---|---|
| ROBO family | 4 | 3,722 | 69% | 98% |
| LRRTM family | 4 | 1,516 | 69% | 98% |
| N-acetylmuramoyl-L-alanine amidase 2 family | 4 | 1,019 | 69% | 95% |
| FAM3 family | 4 | 627 | 68% | 99% |
| Calreticulin family | 4 | 1,371 | 68% | 98% |
| FAM110 family | 4 | 857 | 68% | 99% |
| 2-5A synthase family | 4 | 1,854 | 68% | 99% |
| TNFAIP8 family | 4 | 586 | 68% | 100% |
| Choline/ethanolamine kinase family | 4 | 1,148 | 68% | 97% |
| TSC-22/Dip/Bun family | 4 | 1,617 | 68% | 97% |
| Vestigial family | 4 | 808 | 68% | 97% |
| LTBP family | 4 | 4,388 | 68% | 99% |
| SH3BGR family | 4 | 375 | 68% | 99% |
| Pecanex family | 4 | 5,199 | 68% | 96% |
| Sprouty family | 4 | 825 | 68% | 99% |
| EGF receptor subfamily | 4 | 3,455 | 68% | 97% |
| HAPLN family | 4 | 979 | 67% | 99% |
| Class I subfamily | 4 | 1,332 | 67% | 97% |
| ABCD family | 4 | 1,842 | 67% | 95% |
| Peroxisomal fatty acyl CoA transporter (TC 3,A,1,203) subfamily | 4 | 1,842 | 67% | 95% |
| Aggrecan/versican proteoglycan family | 4 | 5,459 | 67% | 100% |
| Eukaryotic initiation factor 4E family | 4 | 620 | 67% | 98% |
| SEC24 subfamily | 4 | 2,995 | 67% | 96% |
| Phosphatase and actin regulator family | 4 | 1,652 | 67% | 97% |
| IGFL family | 4 | 319 | 67% | 99% |
| Synaptophysin/synaptobrevin family | 4 | 739 | 67% | 98% |
| TEX28 family | 4 | 1,490 | 66% | 97% |
| Fos subfamily | 4 | 870 | 66% | 97% |
| Aldolase class II family | 4 | 1,595 | 66% | 98% |
| Peptidase C64 family | 4 | 2,157 | 66% | 94% |
| Junctophilin family | 4 | 1,798 | 66% | 98% |
| PP1 inhibitor family | 4 | 397 | 66% | 98% |
| WD repeat L(2)GL family | 4 | 2,886 | 65% | 95% |
| Glycosyl hydrolase 35 family | 4 | 1,710 | 65% | 93% |
| Rootletin family | 4 | 2,644 | 65% | 94% |
| JIP scaffold family | 4 | 2,710 | 65% | 95% |
| MIA/OTOR family | 4 | 2,313 | 65% | 94% |
| WNK subfamily | 4 | 4,990 | 65% | 95% |
| Taxilin family | 4 | 1,216 | 64% | 94% |
| G (TC 1,A,1,2) subfamily | 4 | 1,244 | 64% | 95% |
| Epsin family | 4 | 1,589 | 64% | 95% |
| Peptidase S1C family | 4 | 1,198 | 64% | 94% |

| | | | | |
|---|---|---|---|---|
| V-ATPase 116 kDa subunit family | 4 | 2,148 | 64% | 93% |
| Anion channel-forming bestrophin (TC 1,A,46) family | 4 | 1,424 | 64% | 95% |
| Calcium-sensitive chloride channel subfamily | 4 | 1,424 | 64% | 95% |
| SPSB family | 4 | 741 | 64% | 96% |
| PIAS family | 4 | 1,533 | 64% | 95% |
| RGK family | 4 | 790 | 64% | 97% |
| SKI family | 4 | 2,151 | 63% | 93% |
| DMPK subfamily | 4 | 3,564 | 63% | 94% |
| Biopterin-dependent aromatic amino acid hydroxylase family | 4 | 1,211 | 63% | 96% |
| ZC3H12 family | 4 | 1,798 | 63% | 93% |
| MiT/TFE family | 4 | 1,215 | 63% | 94% |
| Dynein intermediate chain family | 4 | 1,620 | 63% | 90% |
| Sal C2H2-type zinc-finger protein family | 4 | 2,924 | 62% | 92% |
| D-isomer specific 2-hydroxyacid dehydrogenase family | 4 | 1,088 | 62% | 89% |
| Calcium-activated (TC 1,A,1,3) subfamily | 4 | 2,956 | 62% | 90% |
| Cornichon family | 4 | 375 | 62% | 92% |
| HEY family | 4 | 749 | 62% | 94% |
| NDRG family | 4 | 917 | 61% | 95% |
| Complexin/synaphin family | 4 | 359 | 61% | 95% |
| Receptor class 2B subfamily | 4 | 3,526 | 61% | 95% |
| ACDP family | 4 | 2,015 | 61% | 94% |
| Teneurin subfamily | 4 | 6,670 | 61% | 95% |
| MEF2 family | 4 | 1,126 | 60% | 89% |
| NXPE family | 4 | 1,320 | 60% | 88% |
| IFI6/IFI27 family | 4 | 289 | 59% | 98% |
| TENT family | 4 | 978 | 59% | 96% |
| Camello family | 4 | 571 | 59% | 93% |
| HIPK subfamily | 4 | 2,499 | 59% | 88% |
| BAR homeobox family | 4 | 734 | 59% | 90% |
| Subgroup III subfamily | 4 | 2,097 | 58% | 85% |
| Receptor class 2A subfamily | 4 | 4,705 | 58% | 89% |
| TMEM7 family | 4 | 560 | 58% | 85% |
| Neurexophilin family | 4 | 633 | 58% | 87% |
| Lammer subfamily | 4 | 1,215 | 58% | 90% |
| Pyrroline-5-carboxylate reductase family | 4 | 735 | 58% | 88% |
| NKAIN family | 4 | 468 | 57% | 92% |
| EYA family | 4 | 1,328 | 57% | 90% |
| Cyclin Y subfamily | 4 | 795 | 57% | 86% |
| Cytochrome c oxidase VIIa family | 4 | 215 | 56% | 91% |
| EMX homeobox family | 4 | 655 | 56% | 86% |
| TCF/LEF family | 4 | 1,117 | 56% | 89% |

| | | | | |
|---|---|---|---|---|
| AhpC/Prx1 subfamily | 4 | 518 | 56% | 93% |
| CDKN2 cyclin-dependent kinase inhibitor family | 4 | 348 | 55% | 80% |
| TEX13 family | 4 | 1,338 | 55% | 95% |
| Stathmin family | 4 | 384 | 55% | 95% |
| NipSnap family | 4 | 576 | 54% | 88% |
| RRM CPEB family | 4 | 1,381 | 53% | 79% |
| CAMP-dependent kinase regulatory chain family | 4 | 844 | 53% | 83% |
| Deformed subfamily | 4 | 578 | 53% | 90% |
| GRB2/sem-5/DRK family | 4 | 466 | 53% | 78% |
| NXF family | 4 | 1,123 | 52% | 77% |
| Fibroblast growth factor receptor subfamily | 4 | 1,669 | 51% | 87% |
| CTF/NF-I family | 4 | 987 | 51% | 82% |
| Rab GDI family | 4 | 1,111 | 50% | 80% |
| Kinesin light chain family | 4 | 1,170 | 50% | 83% |
| Enolase family | 4 | 963 | 50% | 81% |
| HD type 1 subfamily | 4 | 887 | 50% | 76% |
| Type III PI4K subfamily | 4 | 1,879 | 50% | 74% |
| Calcium channel beta subunit family | 4 | 1,126 | 50% | 85% |
| Liprin-alpha subfamily | 4 | 2,388 | 49% | 83% |
| IMPDH/GMPR family | 4 | 849 | 49% | 83% |
| JARID1 histone demethylase family | 4 | 3,111 | 49% | 78% |
| NAC-alpha family | 4 | 1,083 | 49% | 85% |
| NDST subfamily | 4 | 1,706 | 49% | 82% |
| PDE4 subfamily | 4 | 1,512 | 48% | 87% |
| Potassium channel KCNN family | 4 | 1,089 | 48% | 78% |
| Receptors of complement activation (RCA) family | 4 | 1,914 | 48% | 83% |
| Potassium channel HCN family | 4 | 1,772 | 47% | 83% |
| Class-3 subfamily | 4 | 809 | 46% | 77% |
| TALE/TGIF homeobox family | 4 | 490 | 46% | 70% |
| ATP:guanido phosphotransferase family | 4 | 722 | 45% | 79% |
| EHD subfamily | 4 | 968 | 45% | 77% |
| Glycine receptor (TC 1,A,9,3) subfamily | 4 | 809 | 43% | 76% |
| S100-fused protein family | 4 | 3,148 | 43% | 100% |
| KRTAP type 12 family | 4 | 191 | 42% | 75% |
| C (Shaw) (TC 1,A,1,2) subfamily | 4 | 1,068 | 42% | 77% |
| CNKSR family | 4 | 1,318 | 41% | 58% |
| FAH family | 4 | 516 | 41% | 60% |
| G(q) subfamily | 4 | 557 | 38% | 70% |
| Lambda interferon family | 4 | 298 | 38% | 69% |
| Alpha-actinin family | 4 | 1,379 | 38% | 76% |
| Calcitonin family | 4 | 185 | 38% | 59% |
| Parvalbumin family | 4 | 176 | 38% | 62% |

| | | | | |
|---|---|---|---|---|
| TFII-I family | 4 | 1,429 | 37% | 52% |
| Universal ribosomal protein uL10 family | 4 | 417 | 37% | 57% |
| Uroplakin-3 family | 4 | 416 | 37% | 50% |
| RRM elav family | 4 | 527 | 37% | 67% |
| Theta family | 4 | 355 | 37% | 52% |
| Type IIB subfamily | 4 | 1,785 | 36% | 71% |
| METL family | 4 | 476 | 36% | 54% |
| TALE/PBX homeobox family | 4 | 589 | 35% | 72% |
| COE family | 4 | 819 | 35% | 64% |
| Ago subfamily | 4 | 1,171 | 34% | 70% |
| Protein phosphatase inhibitor 2 family | 4 | 271 | 33% | 65% |
| DSH family | 4 | 915 | 32% | 59% |
| Phosphatase 2A regulatory subunit B family | 4 | 564 | 32% | 71% |
| Alkaline phosphatase family | 4 | 656 | 31% | 61% |
| Tropomyosin family | 4 | 333 | 30% | 72% |
| EF-Tu/EF-1A subfamily | 4 | 483 | 26% | 42% |
| GTR/RAG GTP-binding protein family | 4 | 365 | 25% | 44% |
| Archaeal Rpo11/eukaryotic RPB11/RPC19 RNA polymerase subunit family | 4 | 117 | 24% | 56% |
| Gfa family | 4 | 215 | 20% | 31% |
| KRTAP type 1 family | 4 | 124 | 19% | 62% |
| FAM74 family | 4 | 65 | 11% | 24% |
| CCDC144 family | 4 | 382 | 11% | 31% |
| EGF-CFC (Cripto-1/FRL1/Cryptic) family | 4 | 73 | 9% | 19% |
| FAM72 family | 4 | 22 | 4% | 14% |
| POTE family; Actin family | 4 | 132 | 3% | 24% |
| KRTAP type 2 family | 4 | 4 | 1% | 21% |
| FAM236 family | 4 | 0 | 0% | 0% |
| FPG family | 3 | 1,012 | 76% | 100% |
| Carotenoid oxygenase family | 3 | 1,261 | 76% | 100% |
| Copper type II ascorbate-dependent monooxygenase family | 3 | 1,313 | 76% | 100% |
| Uracil-DNA glycosylase (UDG) superfamily | 3 | 750 | 76% | 100% |
| TRM10 family | 3 | 799 | 76% | 100% |
| TBCC family | 3 | 946 | 75% | 100% |
| ATP-dependent DNA ligase family | 3 | 2,135 | 75% | 100% |
| DNA polymerase type-A family | 3 | 3,551 | 75% | 100% |
| Phosphatidylethanolamine-binding protein family | 3 | 595 | 75% | 100% |
| Amidase family | 3 | 1,227 | 75% | 100% |
| RNA methyltransferase TrmH family | 3 | 1,792 | 75% | 100% |
| Protein prenyltransferase subunit alpha family | 3 | 1,009 | 75% | 100% |
| CFAP97 family | 3 | 594 | 75% | 100% |
| Peptidase C78 family | 3 | 889 | 75% | 100% |
| GPAT/DAPAT family | 3 | 1,720 | 75% | 100% |

| | | | | |
|---|---|---|---|---|
| ITPRIP family | 3 | 1,222 | 75% | 100% |
| Peptidase M24A family | 3 | 894 | 75% | 100% |
| ALPK subfamily | 3 | 3,964 | 74% | 100% |
| Pseudouridine synthase TruB family | 3 | 889 | 74% | 100% |
| CD36 family | 3 | 1,118 | 74% | 100% |
| ZC2HC1 family | 3 | 746 | 74% | 100% |
| FAD-dependent oxidoreductase family | 3 | 1,183 | 74% | 100% |
| Protein prenyltransferase subunit beta family | 3 | 851 | 74% | 100% |
| PIGG/PIGN/PIGO family | 3 | 2,231 | 74% | 100% |
| GABA-B receptor subfamily | 3 | 2,017 | 74% | 100% |
| XPF family | 3 | 2,161 | 74% | 100% |
| RNR ribonuclease family | 3 | 2,150 | 74% | 100% |
| UPL family | 3 | 6,659 | 74% | 100% |
| Unc-93 family | 3 | 1,115 | 74% | 100% |
| Fascin family | 3 | 1,100 | 74% | 100% |
| MT-A70-like family | 3 | 1,118 | 74% | 100% |
| FAM188 subfamily | 3 | 1,232 | 74% | 100% |
| Chromogranin/secretogranin protein family | 3 | 1,297 | 74% | 100% |
| Otopetrin family | 3 | 1,310 | 74% | 100% |
| Acid sphingomyelinase family | 3 | 1,139 | 74% | 100% |
| Tectonic family | 3 | 1,399 | 74% | 100% |
| MICU1 family | 3 | 1,065 | 74% | 100% |
| Popeye family | 3 | 750 | 74% | 100% |
| Eukaryotic cobalamin transport proteins family | 3 | 943 | 74% | 100% |
| rRNA adenine N(6)-methyltransferase family | 3 | 779 | 74% | 100% |
| Myozenin family | 3 | 601 | 74% | 100% |
| RIO-type Ser/Thr kinase family | 3 | 1,210 | 74% | 100% |
| UbiA prenyltransferase family | 3 | 850 | 74% | 100% |
| Pannexin family | 3 | 1,103 | 74% | 100% |
| RMDN family | 3 | 880 | 74% | 100% |
| GPN-loop GTPase family | 3 | 713 | 74% | 100% |
| DNA/RNA non-specific endonuclease family | 3 | 858 | 74% | 100% |
| TTC39 family | 3 | 1,383 | 74% | 100% |
| SLC46A family | 3 | 1,027 | 74% | 100% |
| MTFR1 family | 3 | 743 | 74% | 100% |
| CIMIP2 family | 3 | 583 | 74% | 100% |
| Type A subfamily | 3 | 1,021 | 74% | 100% |
| UBR1 family | 3 | 3,961 | 73% | 100% |
| TFIIB family | 3 | 1,037 | 73% | 100% |
| DNA repair metallo-beta-lactamase (DRMBL) family | 3 | 1,662 | 73% | 100% |
| RELT family | 3 | 737 | 73% | 100% |
| Universal ribosomal protein uL1 family | 3 | 757 | 73% | 100% |

| | | | | |
|---|---|---|---|---|
| RNA methyltransferase RlmE family | 3 | 1,043 | 73% | 100% |
| BolA/IbaG family | 3 | 242 | 73% | 99% |
| TRNA pseudouridine synthase TruA family | 3 | 888 | 73% | 100% |
| Nucleoplasmin family | 3 | 503 | 73% | 100% |
| Retinoblastoma protein (RB) family | 3 | 2,298 | 73% | 100% |
| AKAP95 family | 3 | 1,407 | 73% | 100% |
| TatD-type hydrolase family | 3 | 976 | 73% | 100% |
| Universal ribosomal protein uL30 family | 3 | 486 | 73% | 100% |
| Tyrosinase family | 3 | 1,160 | 73% | 100% |
| Bacterial ribosomal protein bS18 family | 3 | 436 | 73% | 100% |
| Natural cytotoxicity receptor (NCR) family | 3 | 571 | 73% | 100% |
| Ntn-hydrolase family | 3 | 785 | 73% | 100% |
| Alkaline ceramidase family | 3 | 589 | 73% | 100% |
| RLR subfamily | 3 | 1,920 | 73% | 100% |
| Niban family | 3 | 1,731 | 73% | 100% |
| Tachykinin family | 3 | 265 | 73% | 100% |
| Peptidase S28 family | 3 | 1,096 | 73% | 100% |
| Glyoxalase I family | 3 | 479 | 73% | 100% |
| Clarin family | 3 | 503 | 73% | 100% |
| MsrB Met sulfoxide reductase family | 3 | 357 | 73% | 100% |
| Carbohydrate kinase PfkB family | 3 | 715 | 73% | 100% |
| Chibby family | 3 | 594 | 73% | 100% |
| Tweety family | 3 | 1,097 | 73% | 100% |
| TDD superfamily | 3 | 665 | 73% | 100% |
| TAS1R subfamily | 3 | 1,842 | 73% | 100% |
| CD164 family | 3 | 421 | 73% | 100% |
| Aspartyl/asparaginyl beta-hydroxylase family | 3 | 1,103 | 73% | 99% |
| Dysbindin family | 3 | 558 | 73% | 99% |
| VRK subfamily | 3 | 1,001 | 73% | 100% |
| RNA polymerase beta chain family | 3 | 2,500 | 73% | 100% |
| Metaxin family | 3 | 756 | 73% | 100% |
| RAMP family | 3 | 342 | 73% | 100% |
| Fibroblast growth factor-binding protein family | 3 | 519 | 73% | 100% |
| Zyg-11 family | 3 | 1,647 | 73% | 99% |
| ERGIC family | 3 | 762 | 73% | 100% |
| Serine/threonine dehydratase family | 3 | 723 | 73% | 99% |
| FAM171 family | 3 | 1,843 | 73% | 100% |
| CISD protein family | 3 | 268 | 72% | 100% |
| TIM family | 3 | 755 | 72% | 100% |
| Dapper family | 3 | 1,620 | 72% | 100% |
| XRCC4-XLF family | 3 | 607 | 72% | 100% |
| Cation-dependent O-methyltransferase family | 3 | 596 | 72% | 100% |

| | | | | |
|---|---|---|---|---|
| Histone-like Alba family | 3 | 363 | 72% | 100% |
| LISCH7 family | 3 | 1,326 | 72% | 100% |
| Synaptopodin family | 3 | 2,168 | 72% | 100% |
| CBF/MAK21 family | 3 | 1,713 | 72% | 100% |
| Nucleoredoxin family | 3 | 580 | 72% | 100% |
| Adrenomedullin family | 3 | 351 | 72% | 100% |
| RIN (Ras interaction/interference) family | 3 | 1,923 | 72% | 100% |
| SLC35D subfamily | 3 | 800 | 72% | 100% |
| FAM53 family | 3 | 875 | 72% | 100% |
| DCK/DGK family | 3 | 579 | 72% | 99% |
| Peptidase S33 family | 3 | 827 | 72% | 100% |
| Cu-Zn superoxide dismutase family | 3 | 482 | 72% | 100% |
| CDIP1/LITAF family | 3 | 318 | 72% | 99% |
| FAM13 family | 3 | 1,819 | 72% | 99% |
| NHS family | 3 | 3,234 | 72% | 100% |
| Smoothelin family | 3 | 1,349 | 72% | 99% |
| Orn/Lys/Arg decarboxylase class-II family | 3 | 986 | 72% | 99% |
| Peptidase S10 family | 3 | 1,014 | 72% | 100% |
| CARMIL family | 3 | 3,008 | 72% | 100% |
| ITM2 family | 3 | 573 | 72% | 100% |
| IRG family | 3 | 912 | 72% | 100% |
| ZHX family | 3 | 1,919 | 72% | 99% |
| Epoxide hydrolase family | 3 | 919 | 72% | 100% |
| Exonuclease superfamily | 3 | 526 | 72% | 100% |
| TET family | 3 | 4,269 | 72% | 99% |
| TMEM45 family | 3 | 592 | 72% | 100% |
| WD repeat CDC20/Fizzy family | 3 | 1,089 | 72% | 100% |
| TMEM8 family | 3 | 1,053 | 72% | 99% |
| ABCF family | 3 | 1,565 | 72% | 100% |
| EF3 subfamily | 3 | 1,565 | 72% | 100% |
| FAM240 family | 3 | 184 | 72% | 99% |
| TM2 family | 3 | 480 | 72% | 100% |
| POU2AF family | 3 | 571 | 72% | 100% |
| Nanos family | 3 | 433 | 72% | 100% |
| Pentraxin family | 3 | 419 | 72% | 99% |
| Dpy-30 family | 3 | 325 | 72% | 100% |
| BCLAF1/THRAP3 family | 3 | 1,855 | 72% | 99% |
| NSG family | 3 | 411 | 72% | 100% |
| GLTP family | 3 | 512 | 72% | 100% |
| ENTREP family | 3 | 1,188 | 72% | 100% |
| LAPTM4/LAPTM5 transporter family | 3 | 582 | 72% | 100% |
| SEZ6 family | 3 | 2,098 | 72% | 100% |

| | | | | |
|---|---|---|---|---|
| TACC family | 3 | 3,289 | 72% | 99% |
| SLRP class III subfamily | 3 | 682 | 72% | 99% |
| Pex2/pex10/pex12 family | 3 | 709 | 72% | 100% |
| CIDE family | 3 | 484 | 72% | 100% |
| OXR1 family | 3 | 1,454 | 72% | 100% |
| NNMT/PNMT/TEMT family | 3 | 579 | 72% | 100% |
| Cornifelin family | 3 | 289 | 72% | 100% |
| AAA ATPase family; Peptidase M41 family | 3 | 1,691 | 72% | 100% |
| Extended synaptotagmin family | 3 | 2,081 | 71% | 100% |
| RING-box family | 3 | 218 | 71% | 100% |
| CCDC88 family | 3 | 3,841 | 71% | 99% |
| Selectin/LECAM family | 3 | 1,293 | 71% | 100% |
| FAM174 family | 3 | 343 | 71% | 100% |
| Plasma kallikrein subfamily | 3 | 1,182 | 71% | 100% |
| CHP subfamily | 3 | 431 | 71% | 100% |
| RIPOR family | 3 | 2,306 | 71% | 99% |
| Aconitase/IPM isomerase family | 3 | 1,875 | 71% | 99% |
| Cytochrome c oxidase subunit 6B family | 3 | 213 | 71% | 99% |
| RRM Spen family | 3 | 3,940 | 71% | 99% |
| AMIGO family | 3 | 1,082 | 71% | 100% |
| Peroxin-11 family | 3 | 532 | 71% | 100% |
| Amer family | 3 | 1,899 | 71% | 99% |
| Eukaryotic NMN adenylyltransferase family | 3 | 596 | 71% | 98% |
| FAM20 family | 3 | 1,091 | 71% | 98% |
| Concentrative nucleoside transporter (CNT) (TC 2,A,41) family | 3 | 1,421 | 71% | 100% |
| Mrp/NBP35 ATP-binding proteins family | 3 | 647 | 71% | 99% |
| Natriuretic peptide family | 3 | 292 | 71% | 100% |
| Opioid neuropeptide precursor family | 3 | 495 | 71% | 100% |
| CSC1 (TC 1,A,17) family | 3 | 1,736 | 71% | 99% |
| TMEM200 family | 3 | 1,007 | 71% | 100% |
| APP family | 3 | 1,549 | 71% | 100% |
| Peroxiredoxin-like PRXL2 family | 3 | 463 | 71% | 100% |
| Peptidase M3 family | 3 | 1,493 | 71% | 98% |
| LanC-like protein family | 3 | 899 | 71% | 99% |
| PA-PLA1 family | 3 | 1,849 | 71% | 97% |
| BCOR family | 3 | 2,609 | 71% | 99% |
| SnRNP core protein family | 3 | 257 | 71% | 99% |
| TMEM106 family | 3 | 556 | 71% | 99% |
| Spinster (TC 2,A,1,49) family | 3 | 1,124 | 71% | 99% |
| SLRP class IV subfamily | 3 | 1,133 | 71% | 100% |
| Paxillin family | 3 | 1,017 | 71% | 99% |
| RETREG family | 3 | 1,065 | 71% | 100% |

| | | | | |
|---|---|---|---|---|
| Asx family | 3 | 3,692 | 71% | 99% |
| Parathyroid hormone family | 3 | 277 | 71% | 99% |
| Calsyntenin family | 3 | 2,043 | 71% | 99% |
| FAM131 family | 3 | 690 | 71% | 99% |
| SNAP-25 family | 3 | 476 | 71% | 100% |
| Myotilin/palladin family | 3 | 2,256 | 70% | 99% |
| MPI phosphatase family | 3 | 1,111 | 70% | 98% |
| Sterol o-acyltransferase subfamily | 3 | 1,098 | 70% | 97% |
| Transferrin family | 3 | 1,508 | 70% | 99% |
| WEE1 subfamily | 3 | 1,202 | 70% | 100% |
| RNA polymerase beta' chain family | 3 | 3,566 | 70% | 99% |
| Glycosyl hydrolase 20 family | 3 | 1,102 | 70% | 99% |
| MCF2 family | 3 | 2,226 | 70% | 99% |
| PA28 family | 3 | 520 | 70% | 99% |
| Non-receptor class 4 subfamily | 3 | 1,434 | 70% | 99% |
| CAMSAP1 family | 3 | 3,040 | 70% | 100% |
| ATR family | 3 | 1,178 | 70% | 97% |
| Hook family | 3 | 1,514 | 70% | 99% |
| SRC/p160 nuclear receptor coactivator family | 3 | 3,027 | 70% | 99% |
| TCL1 family | 3 | 244 | 70% | 100% |
| GW182 family | 3 | 3,834 | 70% | 99% |
| SFT2 family | 3 | 373 | 70% | 100% |
| Protein phosphatase inhibitor 1 family | 3 | 338 | 70% | 100% |
| ADP-ribosylglycohydrolase family | 3 | 750 | 70% | 100% |
| Polycomblike family | 3 | 1,215 | 70% | 100% |
| UDPGP type 1 family | 3 | 1,073 | 70% | 98% |
| P4HA family | 3 | 1,126 | 70% | 98% |
| Beta3-Gal-T subfamily | 3 | 695 | 70% | 99% |
| MAP1 family | 3 | 4,414 | 70% | 98% |
| Cortexin family | 3 | 170 | 70% | 100% |
| Colipase family | 3 | 232 | 70% | 99% |
| HINT family | 3 | 328 | 70% | 99% |
| WWC family | 3 | 2,365 | 70% | 99% |
| LAP (LRR and PDZ) protein family | 3 | 3,187 | 70% | 98% |
| BOD1 family | 3 | 2,371 | 70% | 97% |
| Myelin proteolipid protein family | 3 | 570 | 70% | 99% |
| Rad21 family | 3 | 1,205 | 69% | 97% |
| SLC43A transporter (TC 2,A,1,44) family | 3 | 1,125 | 69% | 98% |
| MDFI family | 3 | 473 | 69% | 98% |
| SAPS family | 3 | 1,885 | 69% | 99% |
| GSG1 family | 3 | 674 | 69% | 99% |
| ERG4/ERG24 family | 3 | 1,044 | 69% | 98% |

| | | | | |
|---|---|---|---|---|
| AXL/UFO subfamily | 3 | 1,926 | 69% | 98% |
| TORC family | 3 | 1,345 | 69% | 99% |
| Nogo receptor family | 3 | 922 | 69% | 100% |
| GRB7/10/14 family | 3 | 1,151 | 69% | 98% |
| JHDM2 histone demethylase family | 3 | 3,879 | 69% | 98% |
| SH2B adapter family | 3 | 1,354 | 69% | 98% |
| Glucose-6-phosphatase family | 3 | 729 | 69% | 99% |
| EVA1 family | 3 | 522 | 69% | 97% |
| ODC antizyme family | 3 | 449 | 69% | 98% |
| BTD/VNN family | 3 | 1,085 | 69% | 98% |
| MHC peptide exporter (TC 3,A,1,209) subfamily | 3 | 1,514 | 69% | 97% |
| SYCE family | 3 | 558 | 69% | 99% |
| Archaeal RpoM/eukaryotic RPA12/RPB9/RPC11 RNA polymerase family | 3 | 247 | 69% | 99% |
| CD99 family | 3 | 431 | 69% | 100% |
| FNBP1 family | 3 | 1,253 | 69% | 99% |
| CDC50/LEM3 family | 3 | 567 | 69% | 97% |
| Arginase family | 3 | 706 | 69% | 99% |
| IER family | 3 | 655 | 69% | 99% |
| Sphingomyelin synthase family | 3 | 819 | 69% | 96% |
| Eukaryotic initiation factor 4G family | 3 | 2,808 | 69% | 96% |
| Caveolin family | 3 | 337 | 69% | 99% |
| CCDC85 family | 3 | 805 | 69% | 100% |
| AXUD1 family | 3 | 1,177 | 69% | 99% |
| Ripply family | 3 | 321 | 68% | 100% |
| Vesicular transporter family | 3 | 1,074 | 68% | 99% |
| TMEM184 family | 3 | 860 | 68% | 97% |
| H2,0 homeobox family | 3 | 799 | 68% | 98% |
| Delta-type subfamily | 3 | 672 | 68% | 99% |
| HHIP family | 3 | 1,505 | 68% | 98% |
| Diaphanous subfamily | 3 | 2,429 | 68% | 98% |
| SOGA family | 3 | 2,910 | 68% | 98% |
| LRRC3 family | 3 | 538 | 68% | 98% |
| Mitochondrial pyruvate carrier (MPC) (TC 2,A,105) family | 3 | 253 | 68% | 99% |
| Polycystin subfamily | 3 | 1,153 | 68% | 96% |
| Iodothyronine deiodinase family | 3 | 560 | 68% | 98% |
| Repulsive guidance molecule (RGM) family | 3 | 890 | 68% | 98% |
| SH3RF family | 3 | 1,688 | 68% | 99% |
| PRA1 family | 3 | 372 | 68% | 99% |
| MEGF family | 3 | 2,173 | 67% | 99% |
| CDI family | 3 | 457 | 67% | 100% |
| PKI family | 3 | 155 | 67% | 100% |
| ANKRD34 family | 3 | 1,041 | 67% | 98% |

| | | | | |
|---|---|---|---|---|
| N-CoR nuclear receptor corepressors family | 3 | 3,406 | 67% | 97% |
| NOS family | 3 | 2,551 | 67% | 98% |
| FSH/LSH/TSH subfamily | 3 | 1,447 | 67% | 97% |
| GPR137 family | 3 | 834 | 67% | 97% |
| Peptidase C69 family | 3 | 846 | 67% | 98% |
| Secernin subfamily | 3 | 846 | 67% | 98% |
| JHDM1D subfamily | 3 | 2,071 | 67% | 96% |
| TRAM family | 3 | 744 | 67% | 97% |
| Nav/unc-53 family | 3 | 4,512 | 67% | 97% |
| Verprolin family | 3 | 953 | 67% | 100% |
| Angiomotin family | 3 | 1,883 | 67% | 98% |
| AB hydrolase 2 family | 3 | 466 | 67% | 97% |
| SLC34A transporter family | 3 | 1,287 | 67% | 97% |
| Nucleobase:cation symporter-2 (NCS2) (TC 2,A,40) family | 3 | 1,240 | 67% | 95% |
| GPRASP family | 3 | 1,853 | 67% | 95% |
| GGA protein family | 3 | 1,316 | 67% | 98% |
| TTC9 family | 3 | 421 | 67% | 99% |
| Tribbles subfamily | 3 | 714 | 67% | 97% |
| CSMD family | 3 | 7,152 | 66% | 98% |
| Glycosyltransferase 25 family | 3 | 1,225 | 66% | 99% |
| Glycosyltransferase 43 family | 3 | 659 | 66% | 98% |
| P53 family | 3 | 1,135 | 66% | 96% |
| WD repeat neurobeachin family | 3 | 5,571 | 66% | 96% |
| Endothelin/sarafotoxin family | 3 | 416 | 66% | 97% |
| Rho GDI family | 3 | 417 | 66% | 97% |
| SORCS subfamily | 3 | 2,349 | 66% | 95% |
| Paraoxonase family | 3 | 703 | 66% | 99% |
| SS18 family | 3 | 589 | 66% | 95% |
| PAR subfamily | 3 | 610 | 66% | 97% |
| Mical family | 3 | 3,319 | 66% | 95% |
| CITED family | 3 | 427 | 66% | 99% |
| Class II subfamily | 3 | 744 | 66% | 94% |
| Gamma-type subfamily | 3 | 731 | 66% | 95% |
| Copper/topaquinone oxidase family | 3 | 1,489 | 66% | 92% |
| TPPP family | 3 | 370 | 65% | 98% |
| FAM122 family | 3 | 477 | 65% | 98% |
| Universal ribosomal protein uL3 family | 3 | 757 | 65% | 94% |
| DACH/dachshund family | 3 | 1,480 | 65% | 98% |
| Kindlin family | 3 | 1,322 | 65% | 97% |
| TOM1 family | 3 | 963 | 65% | 95% |
| Adducin subfamily | 3 | 1,416 | 65% | 98% |
| SLC41A transporter family | 3 | 1,037 | 65% | 96% |

| | | | | |
|---|---|---|---|---|
| JAKMIP family | 3 | 1,483 | 65% | 98% |
| S (TC 1,A,1,2) subfamily | 3 | 971 | 65% | 98% |
| Type B subfamily | 3 | 620 | 64% | 96% |
| VPS26 family | 3 | 618 | 64% | 92% |
| Phosphorylase b kinase regulatory chain family | 3 | 2,280 | 64% | 95% |
| Transketolase family | 3 | 1,183 | 64% | 99% |
| Labial subfamily | 3 | 618 | 64% | 94% |
| HIN-200 family | 3 | 1,037 | 64% | 97% |
| PWWP3A family | 3 | 2,213 | 64% | 100% |
| Troponin T family | 3 | 537 | 64% | 97% |
| JADE family | 3 | 1,560 | 64% | 95% |
| Lipin family | 3 | 1,675 | 64% | 95% |
| Ena/VASP family | 3 | 881 | 64% | 97% |
| IRF2BP family | 3 | 1,249 | 63% | 92% |
| Malic enzymes family | 3 | 1,117 | 63% | 96% |
| Fem-1 family | 3 | 1,211 | 63% | 93% |
| Constitutive coactivator of PPAR-gamma family | 3 | 1,976 | 63% | 93% |
| GADD45 family | 3 | 306 | 63% | 98% |
| PDE6D/unc-119 family | 3 | 405 | 63% | 96% |
| GAMAD family | 3 | 200 | 63% | 91% |
| BRAG family | 3 | 2,292 | 63% | 95% |
| V-ATPase G subunit family | 3 | 223 | 63% | 91% |
| Sestrin family | 3 | 922 | 63% | 95% |
| Grainyhead subfamily | 3 | 1,176 | 63% | 93% |
| 5'-AMP-activated protein kinase gamma subunit family | 3 | 873 | 63% | 94% |
| FAM98 family | 3 | 816 | 63% | 94% |
| Homer family | 3 | 666 | 62% | 95% |
| PI transfer class IIA subfamily | 3 | 2,220 | 62% | 95% |
| MCAK/KIF2 subfamily | 3 | 1,309 | 62% | 94% |
| Troponin I family | 3 | 360 | 62% | 99% |
| PACSIN family | 3 | 840 | 62% | 92% |
| BRINP family | 3 | 1,430 | 62% | 92% |
| L-isoaspartyl/D-aspartyl protein methyltransferase family | 3 | 583 | 62% | 92% |
| WD repeat striatin family | 3 | 1,435 | 62% | 92% |
| EIF4E-binding protein family | 3 | 208 | 62% | 91% |
| Jun subfamily | 3 | 630 | 61% | 97% |
| GIPC family | 3 | 589 | 61% | 94% |
| Histatin/statherin family | 3 | 104 | 61% | 94% |
| Cytidylyltransferase family | 3 | 687 | 61% | 87% |
| Parvin family | 3 | 648 | 61% | 92% |
| RRM TET family | 3 | 1,077 | 61% | 97% |
| Derlin family | 3 | 439 | 61% | 94% |

| | | | | |
|---|---|---|---|---|
| MYT1 family | 3 | 2,029 | 60% | 94% |
| Class-2 subfamily | 3 | 1,003 | 60% | 89% |
| SCAR/WAVE family | 3 | 940 | 60% | 96% |
| Peptidase M19 family | 3 | 833 | 60% | 91% |
| HMX homeobox family | 3 | 587 | 60% | 93% |
| CEP170 family | 3 | 2,077 | 60% | 85% |
| CoREST family | 3 | 899 | 60% | 91% |
| RCAN family | 3 | 412 | 60% | 93% |
| Cyclin D subfamily | 3 | 521 | 59% | 93% |
| Synapsin family | 3 | 1,110 | 59% | 92% |
| PAR6 family | 3 | 650 | 59% | 94% |
| Katanin p60 subunit A1 subfamily | 3 | 900 | 59% | 88% |
| FAM9 family | 3 | 405 | 59% | 96% |
| Synuclein family | 3 | 236 | 59% | 93% |
| PDE1 subfamily | 3 | 1,046 | 59% | 93% |
| RAF subfamily | 3 | 1,187 | 59% | 94% |
| Sulfotransferase 6 family | 3 | 873 | 59% | 90% |
| Caudal homeobox family | 3 | 506 | 59% | 92% |
| BCL7 family | 3 | 369 | 59% | 91% |
| NR4 subfamily | 3 | 1,066 | 59% | 88% |
| Hedgehog family | 3 | 742 | 58% | 87% |
| MAPRE family | 3 | 512 | 58% | 93% |
| Fibrillin family | 3 | 5,015 | 58% | 95% |
| KHDRBS family | 3 | 663 | 58% | 91% |
| FAM27 family | 3 | 229 | 58% | 90% |
| ABI family | 3 | 803 | 58% | 88% |
| FMR1 family | 3 | 1,112 | 58% | 92% |
| Filamin family | 3 | 4,600 | 58% | 93% |
| SNAP family | 3 | 522 | 58% | 85% |
| Teashirt C2H2-type zinc-finger protein family | 3 | 1,839 | 58% | 88% |
| Luc7 family | 3 | 688 | 58% | 91% |
| Kinesin II subfamily | 3 | 1,289 | 58% | 94% |
| EF-1-beta/EF-1-delta family | 3 | 367 | 57% | 90% |
| Orai family | 3 | 487 | 57% | 92% |
| PI3K p85 subunit family | 3 | 1,094 | 57% | 91% |
| RRM IMP/VICKZ family | 3 | 999 | 57% | 90% |
| Di-Ras family | 3 | 356 | 57% | 87% |
| NodC/HAS family | 3 | 957 | 57% | 88% |
| PROL1/PROL3 family | 3 | 261 | 57% | 86% |
| Eukaryotic mitochondrial porin family | 3 | 485 | 56% | 93% |
| PIM subfamily | 3 | 534 | 56% | 89% |
| Ryanodine receptor (TC 1,A,3,1) family | 3 | 8,345 | 56% | 89% |

| | | | | |
|---|---|---|---|---|
| Alpha-ketoglutarate dehydrogenase family | 3 | 1,645 | 56% | 85% |
| InsP3 receptor family | 3 | 4,526 | 56% | 88% |
| Universal ribosomal protein uL13 family | 3 | 267 | 55% | 82% |
| CTAG/PCC1 family | 3 | 294 | 55% | 80% |
| CBFA2T family | 3 | 1,023 | 55% | 85% |
| ATP-dependent PFK group I subfamily | 3 | 1,288 | 55% | 90% |
| Eukaryotic two domain clade 'E' sub-subfamily | 3 | 1,288 | 55% | 90% |
| Phosphofructokinase type A (PFKA) family | 3 | 1,288 | 55% | 90% |
| SINA (Seven in absentia) family | 3 | 480 | 55% | 87% |
| SLC8 subfamily | 3 | 1,545 | 55% | 89% |
| SMARCD family | 3 | 834 | 55% | 86% |
| Ficolin lectin family | 3 | 504 | 54% | 88% |
| PUR DNA-binding protein family | 3 | 527 | 54% | 88% |
| Aurora subfamily | 3 | 567 | 54% | 90% |
| Muscleblind family | 3 | 594 | 53% | 87% |
| SMEK family | 3 | 1,336 | 53% | 87% |
| YTHDF family | 3 | 915 | 53% | 88% |
| Requiem/DPF family | 3 | 611 | 53% | 90% |
| C2CD4 family | 3 | 607 | 53% | 82% |
| Unc-104 subfamily | 3 | 2,390 | 52% | 84% |
| Pellino family | 3 | 677 | 52% | 89% |
| Riboflavin transporter family | 3 | 700 | 51% | 81% |
| Shaker potassium channel beta subunit family | 3 | 609 | 51% | 87% |
| BACURD family | 3 | 485 | 51% | 85% |
| TBP family | 3 | 454 | 50% | 79% |
| Centrin family | 3 | 256 | 50% | 84% |
| WD repeat DCAF8 family | 3 | 914 | 50% | 88% |
| TAF9 family | 3 | 326 | 50% | 79% |
| Type II pantothenate kinase family | 3 | 766 | 50% | 87% |
| GATS family | 3 | 407 | 50% | 77% |
| UTX family | 3 | 2,171 | 49% | 77% |
| DIP2 family | 3 | 2,323 | 49% | 83% |
| KRTAP type 6 family | 3 | 116 | 49% | 94% |
| Glucosamine/galactosamine-6-phosphate isomerase family | 3 | 401 | 49% | 79% |
| F-actin-capping protein alpha subunit family | 3 | 421 | 48% | 74% |
| CP2 subfamily | 3 | 732 | 48% | 84% |
| TCTP family | 3 | 214 | 47% | 86% |
| D (Shal) (TC 1,A,1,2) subfamily | 3 | 914 | 47% | 76% |
| Kinesin subfamily | 3 | 1,394 | 47% | 83% |
| ARD1 subfamily | 3 | 303 | 47% | 73% |
| Class I fructose-bisphosphate aldolase family | 3 | 514 | 47% | 80% |
| Poly(A) polymerase family | 3 | 995 | 47% | 75% |

| | | | | |
|---|---|---|---|---|
| PKD subfamily | 3 | 1,240 | 46% | 81% |
| MORF4 family-associated protein family | 3 | 172 | 46% | 73% |
| ORM family | 3 | 209 | 46% | 78% |
| ERD2 family | 3 | 287 | 45% | 79% |
| TVP23 family | 3 | 309 | 45% | 66% |
| Dynein light chain family | 3 | 126 | 45% | 68% |
| ABHD17 family | 3 | 411 | 44% | 80% |
| WD repeat DCAF12 family | 3 | 611 | 44% | 76% |
| Adenosylhomocysteinase family | 3 | 689 | 44% | 69% |
| Class-4 subfamily | 3 | 497 | 43% | 76% |
| Lin-7 family | 3 | 264 | 41% | 84% |
| VGLUT subfamily | 3 | 717 | 41% | 75% |
| SAA family | 3 | 153 | 41% | 68% |
| Non-receptor class CDC14 subfamily | 3 | 628 | 41% | 64% |
| Glycogen phosphorylase family | 3 | 1,018 | 40% | 75% |
| Melanin-concentrating hormone family | 3 | 135 | 40% | 77% |
| eIF4A subfamily | 3 | 488 | 40% | 70% |
| RAC subfamily | 3 | 570 | 40% | 78% |
| Ropporin family | 3 | 254 | 39% | 63% |
| Universal ribosomal protein uL16 family | 3 | 261 | 38% | 58% |
| SPT20 family | 3 | 908 | 38% | 60% |
| Class-5 subfamily | 3 | 390 | 37% | 62% |
| G(s) subfamily | 3 | 668 | 37% | 58% |
| MIF family | 3 | 135 | 37% | 51% |
| PP-2B subfamily | 3 | 548 | 35% | 73% |
| Universal ribosomal protein uL24 family | 3 | 177 | 35% | 51% |
| Peptidase S26B family | 3 | 180 | 34% | 65% |
| Securin family | 3 | 199 | 33% | 72% |
| ATPase C chain family | 3 | 134 | 32% | 46% |
| RRN3 family | 3 | 352 | 31% | 50% |
| ZXD family | 3 | 735 | 30% | 49% |
| WD repeat EBI family | 3 | 459 | 28% | 64% |
| FKBP1 subfamily | 3 | 91 | 28% | 56% |
| Universal ribosomal protein uS2 family | 3 | 245 | 28% | 41% |
| AQP11/AQP12 subfamily | 3 | 228 | 26% | 39% |
| KRTAP type 3 family | 3 | 74 | 25% | 47% |
| Aldo/keto reductase 2 subfamily | 3 | 254 | 25% | 59% |
| Sedlin subfamily | 3 | 103 | 25% | 33% |
| FAM246 family | 3 | 170 | 24% | 47% |
| Eukaryotic ribosomal protein eL39 family | 3 | 34 | 22% | 60% |
| Stereocilin family | 3 | 1,004 | 21% | 31% |
| Eukaryotic ribosomal protein eS4 family | 3 | 161 | 20% | 51% |

| | | | | |
|---|---|---|---|---|
| Paired-like homeobox family | 3 | 151 | 20% | 30% |
| PEPP subfamily | 3 | 151 | 20% | 30% |
| CK2 subfamily | 3 | 209 | 18% | 38% |
| EIF-5A family | 3 | 77 | 17% | 43% |
| TMEM191 family | 3 | 135 | 16% | 25% |
| FAM10 family | 3 | 148 | 15% | 40% |
| FAM127 family | 3 | 48 | 14% | 42% |
| ANKRD36 family | 3 | 684 | 14% | 41% |
| DDX11/CHL1 sub-subfamily | 3 | 356 | 13% | 42% |
| UPF0633 family | 3 | 35 | 12% | 34% |
| FRG2 family | 3 | 97 | 12% | 29% |
| FAM157 family | 3 | 101 | 9% | 34% |
| Nanog homeobox family | 3 | 60 | 7% | 24% |
| LRRC37A family | 3 | 224 | 4% | 15% |
| FAM25 family | 3 | 0 | 0% | 0% |
| Sulfotransferase 3 family | 2 | 589 | 77% | 100% |
| Universal ribosomal protein uL23 family | 2 | 237 | 77% | 99% |
| KAE1 / TsaD family | 2 | 574 | 77% | 100% |
| Universal ribosomal protein uL14 family | 2 | 218 | 76% | 99% |
| Clp1 family | 2 | 858 | 76% | 100% |
| MINAR family | 2 | 842 | 76% | 100% |
| Eukaryotic RPB7/RPC8 RNA polymerase subunit family | 2 | 286 | 76% | 100% |
| NCF2/NOXA1 family | 2 | 762 | 76% | 100% |
| TM6SF family | 2 | 568 | 76% | 100% |
| FAM187 family | 2 | 594 | 76% | 100% |
| IL-15/IL-21 family | 2 | 246 | 76% | 99% |
| Crooked-neck family | 2 | 1,293 | 76% | 100% |
| Glycoprotein hormones subunit alpha family | 2 | 186 | 76% | 100% |
| UPF0224 (FAM112) family | 2 | 239 | 76% | 100% |
| FAM111 family | 2 | 1,020 | 76% | 100% |
| Stanniocalcin family | 2 | 416 | 76% | 100% |
| B9D family | 2 | 287 | 76% | 100% |
| MDH type 2 family | 2 | 645 | 76% | 100% |
| PMEL/NMB family | 2 | 933 | 76% | 100% |
| TASOR family | 2 | 3,102 | 76% | 100% |
| Sclerostin family | 2 | 317 | 76% | 100% |
| Universal ribosomal protein uS11 family | 2 | 261 | 76% | 100% |
| BUB1 subfamily | 2 | 1,615 | 76% | 100% |
| Transferase hexapeptide repeat family | 2 | 590 | 76% | 100% |
| Eukaryotic-type primase small subunit family | 2 | 741 | 76% | 100% |
| TMEM131 family | 2 | 2,640 | 76% | 100% |
| RIB43A family | 2 | 575 | 76% | 100% |

| | | | | |
|---|---|---|---|---|
| Queuine tRNA-ribosyltransferase family | 2 | 618 | 76% | 100% |
| TPP enzyme family | 2 | 914 | 76% | 100% |
| TCP10 family | 2 | 1,171 | 75% | 100% |
| Universal ribosomal protein uS4 family | 2 | 285 | 75% | 99% |
| Shugoshin family | 2 | 1,376 | 75% | 100% |
| PNP/MTAP phosphorylase family | 2 | 431 | 75% | 100% |
| Universal ribosomal protein uS17 family | 2 | 217 | 75% | 100% |
| HSBP1 family | 2 | 113 | 75% | 99% |
| Palmitoyl-protein thioesterase family | 2 | 458 | 75% | 99% |
| Spermidine/spermine synthase family | 2 | 503 | 75% | 100% |
| TMEM126 family | 2 | 320 | 75% | 100% |
| ATM subfamily | 2 | 4,291 | 75% | 100% |
| EPO/TPO family | 2 | 411 | 75% | 100% |
| Glycosyltransferase 61 family | 2 | 833 | 75% | 100% |
| FAM227 family | 2 | 811 | 75% | 100% |
| Prominin family | 2 | 1,278 | 75% | 100% |
| Universal ribosomal protein uL4 family | 2 | 555 | 75% | 100% |
| Complex I NDUFA12 subunit family | 2 | 236 | 75% | 99% |
| DAMOX/DASOX family | 2 | 517 | 75% | 100% |
| GTP-binding SRP family | 2 | 858 | 75% | 100% |
| Peptidase C13 family | 2 | 622 | 75% | 100% |
| Type-1 seryl-tRNA synthetase subfamily | 2 | 775 | 75% | 100% |
| FAM217 family | 2 | 669 | 75% | 100% |
| Thymidylate kinase family | 2 | 496 | 75% | 100% |
| Methionine aminopeptidase type 1 subfamily | 2 | 541 | 75% | 100% |
| Very large inducible GTPase (VLIG) family | 2 | 2,515 | 75% | 100% |
| CWF19 family | 2 | 1,074 | 75% | 100% |
| FAM118 family | 2 | 531 | 75% | 100% |
| TMEM179 family | 2 | 339 | 75% | 100% |
| Pseudouridine synthase TruD family | 2 | 1,021 | 75% | 100% |
| AccD/PCCB family | 2 | 826 | 75% | 100% |
| UTP23/FCF1 family | 2 | 335 | 75% | 100% |
| Peptidase M54 family | 2 | 643 | 75% | 100% |
| Eukaryotic release factor 1 family | 2 | 616 | 75% | 100% |
| FAM172 family | 2 | 583 | 75% | 100% |
| Trm1 family | 2 | 1,043 | 75% | 100% |
| Nicastrin family | 2 | 953 | 75% | 100% |
| Universal ribosomal protein uL2 family | 2 | 421 | 75% | 100% |
| Threonine synthase family | 2 | 919 | 75% | 100% |
| PhyH family | 2 | 471 | 75% | 100% |
| DeSI family | 2 | 271 | 75% | 100% |
| Otulin subfamily | 2 | 530 | 75% | 100% |

| | | | | |
|---|---|---|---|---|
| SDE3 subfamily | 2 | 1,657 | 75% | 100% |
| Otogelin family | 2 | 3,949 | 75% | 100% |
| HIBADH-related family | 2 | 665 | 75% | 100% |
| Acid ceramidase family | 2 | 564 | 75% | 100% |
| CD3Z/FCER1G family | 2 | 187 | 75% | 100% |
| Phosphatidyl serine synthase family | 2 | 718 | 75% | 100% |
| CKAP2 family | 2 | 1,068 | 75% | 100% |
| CATSPERD family | 2 | 1,308 | 75% | 100% |
| Peptidase M17 family | 2 | 779 | 75% | 100% |
| FAM161 family | 2 | 977 | 75% | 100% |
| Zona pellucida-binding protein Sp38 family | 2 | 515 | 75% | 100% |
| Caprin family | 2 | 1,372 | 75% | 100% |
| FAM221 family | 2 | 523 | 75% | 100% |
| Nth/MutY family | 2 | 641 | 75% | 100% |
| FAM234 family | 2 | 877 | 75% | 100% |
| CEP43 family | 2 | 428 | 75% | 100% |
| NAD kinase family | 2 | 663 | 75% | 100% |
| FAM154 family | 2 | 651 | 75% | 100% |
| Cation-independent O-methyltransferase family | 2 | 721 | 75% | 100% |
| K-HECT subfamily | 2 | 3,434 | 75% | 100% |
| GASK family | 2 | 816 | 75% | 100% |
| RAD3/XPD subfamily | 2 | 1,476 | 75% | 100% |
| MFSD6 family | 2 | 1,027 | 75% | 100% |
| IF-2 subfamily | 2 | 1,452 | 75% | 100% |
| Purple acid phosphatase family | 2 | 569 | 75% | 100% |
| CCSER family | 2 | 1,293 | 75% | 100% |
| SIKE family | 2 | 343 | 75% | 99% |
| RNA 3'-terminal cyclase family | 2 | 551 | 75% | 100% |
| Sulfatase-modifying factor family | 2 | 503 | 75% | 100% |
| FAM175 family | 2 | 614 | 75% | 100% |
| FAM178 family | 2 | 1,379 | 74% | 100% |
| Glycosyltransferase 32 family | 2 | 516 | 74% | 100% |
| TMCO5 family | 2 | 443 | 74% | 100% |
| Peptidase S16 family | 2 | 1,348 | 74% | 100% |
| SCO1/2 family | 2 | 422 | 74% | 100% |
| 3-hydroxyacyl-CoA dehydrogenase family | 2 | 471 | 74% | 100% |
| Neutral sphingomyelinase family | 2 | 802 | 74% | 100% |
| Peptidase S9A family | 2 | 1,069 | 74% | 100% |
| Glycosyltransferase 13 family | 2 | 822 | 74% | 100% |
| NUP210 family | 2 | 2,808 | 74% | 100% |
| Ubiquitin conjugation factor E4 family | 2 | 1,761 | 74% | 100% |
| FAM222 family | 2 | 754 | 74% | 100% |

| | | | | |
|---|---|---|---|---|
| Universal ribosomal protein uS9 family | 2 | 403 | 74% | 100% |
| Glycosyltransferase 39 family | 2 | 1,113 | 74% | 100% |
| GOLM family | 2 | 622 | 74% | 100% |
| Type IA topoisomerase family | 2 | 1,384 | 74% | 99% |
| DNA repair enzymes AP/ExoA family | 2 | 621 | 74% | 100% |
| TRM61 family | 2 | 569 | 74% | 100% |
| FAM216 family | 2 | 306 | 74% | 100% |
| NFX1 family | 2 | 1,508 | 74% | 100% |
| FAM169 family | 2 | 640 | 74% | 100% |
| P23/wos2 family | 2 | 242 | 74% | 100% |
| Protease inhibitor I47 (latexin) family | 2 | 383 | 74% | 100% |
| NEMP family | 2 | 639 | 74% | 100% |
| Universal ribosomal protein uS7 family | 2 | 331 | 74% | 100% |
| HesB/IscA family | 2 | 210 | 74% | 100% |
| NMI family | 2 | 440 | 74% | 100% |
| TRNA-intron endonuclease family | 2 | 575 | 74% | 100% |
| Mesothelin family | 2 | 988 | 74% | 100% |
| Peptidase M8 family | 2 | 1,070 | 74% | 100% |
| GCF family | 2 | 1,259 | 74% | 100% |
| Motilin family | 2 | 172 | 74% | 100% |
| Translin family | 2 | 384 | 74% | 100% |
| TTC21 family | 2 | 1,954 | 74% | 100% |
| FAM124 family | 2 | 742 | 74% | 100% |
| Inositol 3,4-bisphosphate 4-phosphatase family | 2 | 1,409 | 74% | 99% |
| HEBP family | 2 | 292 | 74% | 100% |
| ADP-ribosyl cyclase family | 2 | 458 | 74% | 100% |
| ATOS family | 2 | 1,196 | 74% | 100% |
| Universal ribosomal protein uL15 family | 2 | 329 | 74% | 100% |
| Clusterin family | 2 | 678 | 74% | 100% |
| SPATA6 family | 2 | 652 | 74% | 100% |
| FAM104 family | 2 | 223 | 74% | 100% |
| TRM5/TYW2 family | 2 | 709 | 74% | 100% |
| EIF-2-beta/eIF-5 family | 2 | 566 | 74% | 100% |
| Phospholipase B-like family | 2 | 846 | 74% | 100% |
| Archaeal Rpo3/eukaryotic RPB3 RNA polymerase subunit family | 2 | 460 | 74% | 100% |
| Peptidase C15 family | 2 | 300 | 74% | 100% |
| Eukaryotic ribosomal protein eL24 family | 2 | 237 | 74% | 100% |
| 2H phosphoesterase superfamily | 2 | 508 | 74% | 100% |
| NAPRTase family | 2 | 762 | 74% | 100% |
| WD repeat SEC13 family | 2 | 505 | 74% | 100% |
| Peptidase M24 family | 2 | 1,067 | 74% | 100% |
| Two pore calcium channel subfamily | 2 | 1,161 | 74% | 100% |

| | | | | |
|---|---|---|---|---|
| Chondromodulin-1 family | 2 | 482 | 74% | 100% |
| Class IV subfamily | 2 | 559 | 74% | 100% |
| ZPB subfamily | 2 | 872 | 74% | 100% |
| HHAT subfamily | 2 | 738 | 74% | 100% |
| Universal ribosomal protein uS12 family | 2 | 208 | 74% | 100% |
| LCMT family | 2 | 755 | 74% | 100% |
| RNase Z family | 2 | 880 | 74% | 100% |
| MDM2/MDM4 family | 2 | 726 | 74% | 100% |
| MRAP family | 2 | 279 | 74% | 100% |
| LicD transferase family | 2 | 707 | 74% | 100% |
| FAM149 family | 2 | 1,002 | 74% | 100% |
| GNAT subfamily | 2 | 278 | 74% | 100% |
| SMN family | 2 | 393 | 74% | 100% |
| SKP1 family | 2 | 203 | 74% | 100% |
| MTUS1 family | 2 | 1,948 | 74% | 100% |
| LCA5 family | 2 | 1,009 | 74% | 100% |
| Prefoldin subunit alpha family | 2 | 259 | 74% | 99% |
| LIF/OSM family | 2 | 335 | 74% | 100% |
| RNA M5U methyltransferase family | 2 | 833 | 74% | 100% |
| ROX family | 2 | 816 | 74% | 100% |
| NOD1-NOD2 family | 2 | 1,470 | 74% | 100% |
| BAF family | 2 | 132 | 74% | 98% |
| CREG family | 2 | 376 | 74% | 100% |
| Crescerin family | 2 | 2,019 | 74% | 100% |
| SYCP2 family | 2 | 1,726 | 74% | 100% |
| CIBAR family | 2 | 437 | 74% | 100% |
| Universal ribosomal protein uL11 family | 2 | 263 | 74% | 100% |
| INSYN2 family | 2 | 747 | 74% | 100% |
| Dynactin subunits 5/6 family | 2 | 274 | 74% | 100% |
| Glycosyl hydrolase 23 family | 2 | 299 | 74% | 100% |
| Methylthiotransferase family | 2 | 869 | 74% | 100% |
| CoA-transferase III family | 2 | 609 | 74% | 100% |
| Apolipoprotein O/MICOS complex subunit Mic27 family | 2 | 343 | 74% | 100% |
| Copper transporter (Ctr) (TC 1,A,56) family | 2 | 245 | 74% | 99% |
| SLC31A subfamily | 2 | 245 | 74% | 99% |
| EIF-2B gamma/epsilon subunits family | 2 | 863 | 74% | 100% |
| Flotillin subfamily | 2 | 629 | 74% | 100% |
| Fetuin family | 2 | 551 | 74% | 100% |
| UPF0606 family | 2 | 2,794 | 74% | 100% |
| Prenylcysteine oxidase family | 2 | 734 | 73% | 99% |
| CNOT2/3/5 family | 2 | 950 | 73% | 100% |
| FIT family | 2 | 407 | 73% | 100% |

| | | | | |
|---|---|---|---|---|
| 5'-3' exonuclease family | 2 | 1,951 | 73% | 99% |
| EME1/MMS4 family | 2 | 697 | 73% | 100% |
| CEMIP family | 2 | 2,015 | 73% | 100% |
| PPase class C family | 2 | 2,600 | 73% | 99% |
| Prune subfamily | 2 | 2,600 | 73% | 99% |
| FILIP1 family | 2 | 1,724 | 73% | 100% |
| Stoned B family | 2 | 1,204 | 73% | 100% |
| Universal ribosomal protein uS3 family | 2 | 301 | 73% | 100% |
| WD repeat DDB2/WDR76 family | 2 | 773 | 73% | 100% |
| Dispatched family | 2 | 2,147 | 73% | 100% |
| CCM2 family | 2 | 745 | 73% | 100% |
| Carotenoid/retinoid oxidoreductase family | 2 | 874 | 73% | 100% |
| WD repeat TAF5 family | 2 | 1,019 | 73% | 100% |
| Neuropilin family | 2 | 1,360 | 73% | 100% |
| PAT1 family | 2 | 963 | 73% | 100% |
| Proline oxidase family | 2 | 833 | 73% | 100% |
| OKL38 family | 2 | 720 | 73% | 100% |
| Transposase 22 family | 2 | 882 | 73% | 100% |
| SAS10 family | 2 | 582 | 73% | 100% |
| Quiescin-sulfhydryl oxidase (QSOX) family | 2 | 1,059 | 73% | 100% |
| SAC3 family | 2 | 1,747 | 73% | 100% |
| TMEM47 family | 2 | 274 | 73% | 100% |
| Cytochrome c family | 2 | 315 | 73% | 100% |
| MLF family | 2 | 378 | 73% | 100% |
| SPOT14 family | 2 | 241 | 73% | 100% |
| Seminal plasma protein family | 2 | 260 | 73% | 100% |
| Lipase maturation factor family | 2 | 933 | 73% | 100% |
| CEP135/TSGA10 family | 2 | 1,346 | 73% | 100% |
| DNase II family | 2 | 528 | 73% | 100% |
| Peptidase M43B family | 2 | 2,503 | 73% | 100% |
| Glycosyl hydrolase 27 family | 2 | 615 | 73% | 100% |
| Rad9 family | 2 | 598 | 73% | 100% |
| SEC16 family | 2 | 2,501 | 73% | 100% |
| KsgA subfamily | 2 | 543 | 73% | 100% |
| MCC family | 2 | 1,121 | 73% | 100% |
| FAM81 family | 2 | 600 | 73% | 100% |
| Universal ribosomal protein uL18 family | 2 | 349 | 73% | 100% |
| WD repeat ATG16 family | 2 | 897 | 73% | 100% |
| TMEM54 family | 2 | 338 | 73% | 100% |
| MCAF family | 2 | 1,428 | 73% | 100% |
| AHA1 family | 2 | 466 | 73% | 100% |
| EPCAM family | 2 | 466 | 73% | 100% |

| | | | | |
|---|---|---|---|---|
| TMEM41 family | 2 | 406 | 73% | 100% |
| MCRIP family | 2 | 188 | 73% | 100% |
| Universal ribosomal protein uS10 family | 2 | 234 | 73% | 100% |
| NTM1 family | 2 | 370 | 73% | 100% |
| SKAP family | 2 | 525 | 73% | 100% |
| 6-phosphogluconolactonase subfamily | 2 | 767 | 73% | 100% |
| CCDC90 family | 2 | 448 | 73% | 100% |
| ECO subfamily | 2 | 1,053 | 73% | 99% |
| AKNA family | 2 | 1,662 | 73% | 100% |
| RPGRIP1 family | 2 | 1,900 | 73% | 100% |
| Universal ribosomal protein uS15 family | 2 | 298 | 73% | 100% |
| MFAP family | 2 | 260 | 73% | 100% |
| WD repeat HIR1 family | 2 | 1,151 | 73% | 100% |
| Peptidase S26 family | 2 | 249 | 73% | 100% |
| Gamma-BBH/TMLD family | 2 | 590 | 73% | 100% |
| CWC16 family | 2 | 525 | 73% | 100% |
| RuvB family | 2 | 671 | 73% | 100% |
| GTPBP1 subfamily | 2 | 928 | 73% | 100% |
| Ribonuclease III family | 2 | 1,245 | 73% | 100% |
| EMC8/EMC9 family | 2 | 305 | 73% | 100% |
| Peroxisomal targeting signal receptor family | 2 | 923 | 73% | 99% |
| GOT1 family | 2 | 197 | 73% | 100% |
| Enoyl-CoA hydratase/isomerase family; 3-hydroxyacyl-CoA dehydrogenase family | 2 | 1,084 | 73% | 100% |
| Xin family | 2 | 3,805 | 73% | 100% |
| DPH1/DPH2 family | 2 | 676 | 73% | 100% |
| Universal ribosomal protein uL29 family | 2 | 272 | 73% | 100% |
| ChaC subfamily | 2 | 296 | 73% | 100% |
| 4HPPD family | 2 | 557 | 73% | 100% |
| DoxX family | 2 | 234 | 73% | 100% |
| Dopey family | 2 | 3,472 | 73% | 100% |
| ZNRF3 family | 2 | 1,253 | 73% | 100% |
| Replication factor A protein 2 family | 2 | 387 | 73% | 100% |
| EMC6 family | 2 | 180 | 73% | 100% |
| AIG1 family | 2 | 341 | 73% | 100% |
| VTI1 family | 2 | 327 | 73% | 100% |
| ADAD family | 2 | 844 | 73% | 100% |
| Universal ribosomal protein uL22 family | 2 | 284 | 73% | 100% |
| Xanthine dehydrogenase family | 2 | 1,945 | 73% | 99% |
| Pyrimidine 5'-nucleotidase family | 2 | 463 | 73% | 100% |
| LRRC14 subfamily | 2 | 733 | 73% | 100% |
| UPF0389 family | 2 | 230 | 73% | 99% |
| TMEM161 family | 2 | 703 | 73% | 100% |

| | | | | |
|---|---|---|---|---|
| HUS1 family | 2 | 406 | 73% | 98% |
| CFAP73 family | 2 | 454 | 73% | 100% |
| Universal ribosomal protein uS5 family | 2 | 526 | 73% | 100% |
| Class-IV pyridoxal-phosphate-dependent aminotransferase family | 2 | 566 | 73% | 100% |
| TAF6 family | 2 | 945 | 73% | 100% |
| CALCOCO family | 2 | 827 | 73% | 100% |
| MICOS complex subunit Mic19 family | 2 | 336 | 73% | 100% |
| Mis18 family | 2 | 336 | 73% | 100% |
| Tom20 family | 2 | 216 | 73% | 100% |
| Bradykinin receptor subfamily | 2 | 541 | 73% | 100% |
| Radical SAM superfamily | 2 | 533 | 73% | 100% |
| CARF family | 2 | 506 | 73% | 100% |
| VMP family | 2 | 290 | 73% | 100% |
| Glutamine synthetase family | 2 | 641 | 73% | 100% |
| TMEM176 family | 2 | 367 | 73% | 100% |
| OXA1/ALB3/YidC family | 2 | 558 | 73% | 100% |
| FNDC3 family | 2 | 1,745 | 73% | 100% |
| FAM184 family | 2 | 1,598 | 73% | 100% |
| Translokin family | 2 | 697 | 73% | 100% |
| TMEM178 family | 2 | 429 | 73% | 100% |
| Isochorismatase family | 2 | 365 | 73% | 99% |
| SPATA2 family | 2 | 685 | 73% | 100% |
| RGS7BP/RGS9BP family | 2 | 357 | 73% | 100% |
| Vasculin family | 2 | 687 | 73% | 99% |
| IFI44 family | 2 | 650 | 73% | 99% |
| Dermatan-sulfate isomerase family | 2 | 1,574 | 73% | 98% |
| Jacalin lectin family | 2 | 272 | 73% | 100% |
| NARF family | 2 | 676 | 73% | 100% |
| SH3BP5 family | 2 | 615 | 73% | 100% |
| Succinate/malate CoA ligase beta subunit family | 2 | 649 | 73% | 100% |
| CbbY/CbbZ/Gph/YieH family | 2 | 398 | 72% | 100% |
| THEM4/THEM5 thioesterase family | 2 | 353 | 72% | 100% |
| DCP1 family | 2 | 869 | 72% | 100% |
| Ariadne subfamily | 2 | 761 | 72% | 100% |
| ATP synthase subunit s family | 2 | 342 | 72% | 100% |
| NifU family | 2 | 305 | 72% | 100% |
| MIP18 family | 2 | 234 | 72% | 100% |
| FAM170 family | 2 | 444 | 72% | 100% |
| LRATD family | 2 | 436 | 72% | 100% |
| Glycosyl hydrolase 79 family | 2 | 822 | 72% | 100% |
| Indoleamine 2,3-dioxygenase family | 2 | 596 | 72% | 100% |
| PTH2 family | 2 | 231 | 72% | 99% |

| | | | | |
|---|---|---|---|---|
| CLPTM1 family | 2 | 874 | 72% | 100% |
| ALG6/ALG8 glucosyltransferase family | 2 | 748 | 72% | 100% |
| SPARC family | 2 | 700 | 72% | 100% |
| TMEM134/TMEM230 family | 2 | 228 | 72% | 100% |
| TMEM38 family | 2 | 427 | 72% | 100% |
| NADP-dependent oxidoreductase L4BD family | 2 | 492 | 72% | 100% |
| DZIP C2H2-type zinc-finger protein family | 2 | 1,182 | 72% | 100% |
| Vacuolar ATPase subunit S1 family | 2 | 502 | 72% | 100% |
| CTNNBIP1 family | 2 | 196 | 72% | 100% |
| ODF2 family | 2 | 1,059 | 72% | 100% |
| Universal ribosomal protein uS14 family | 2 | 133 | 72% | 100% |
| NAD(P)H dehydrogenase (quinone) family | 2 | 365 | 72% | 100% |
| IL-7/IL-9 family | 2 | 232 | 72% | 100% |
| RRP1 family | 2 | 881 | 72% | 100% |
| DUTPase family | 2 | 284 | 72% | 99% |
| FAM228 family | 2 | 383 | 72% | 100% |
| TMEM121 family | 2 | 648 | 72% | 100% |
| CWC22 family | 2 | 1,277 | 72% | 100% |
| Geminin family | 2 | 429 | 72% | 100% |
| ZAR1 family | 2 | 538 | 72% | 100% |
| AUTS2 family | 2 | 1,663 | 72% | 99% |
| Cingulin family | 2 | 1,808 | 72% | 99% |
| PRPH2/ROM1 family | 2 | 503 | 72% | 100% |
| AspA/AstE family | 2 | 456 | 72% | 99% |
| Aspartoacylase subfamily | 2 | 456 | 72% | 99% |
| MVB12 family | 2 | 427 | 72% | 99% |
| Peptidase C56 family | 2 | 295 | 72% | 100% |
| IFRD family | 2 | 690 | 72% | 100% |
| JUPITER family | 2 | 248 | 72% | 100% |
| ATXN1 family | 2 | 1,084 | 72% | 100% |
| Speriolin family | 2 | 671 | 72% | 99% |
| Sel-1 family | 2 | 1,068 | 72% | 100% |
| ATG2 family | 2 | 2,894 | 72% | 99% |
| CEP85 family | 2 | 1,129 | 72% | 100% |
| CC2D1 family | 2 | 1,303 | 72% | 100% |
| FAM229 family | 2 | 149 | 72% | 100% |
| Isthmin family | 2 | 745 | 72% | 100% |
| GrpE family | 2 | 318 | 72% | 99% |
| FAM241 family | 2 | 182 | 72% | 100% |
| LRRC75 family | 2 | 474 | 72% | 100% |
| FAM181 family | 2 | 561 | 72% | 100% |
| INKA family | 2 | 420 | 72% | 100% |

| | | | | |
|---|---|---|---|---|
| KISH family | 2 | 105 | 72% | 100% |
| Bms1-like GTPase family | 2 | 1,500 | 72% | 97% |
| NSFL1C family | 2 | 504 | 72% | 99% |
| Adrenodoxin/putidaredoxin family | 2 | 266 | 72% | 99% |
| DapA family | 2 | 465 | 72% | 100% |
| Ubinuclein family | 2 | 1,783 | 72% | 98% |
| LRRC32/LRRC33 family | 2 | 973 | 72% | 100% |
| EFCAB4 family | 2 | 812 | 72% | 100% |
| Sarcoglycan alpha/epsilon family | 2 | 592 | 72% | 99% |
| Mitoguardin family | 2 | 880 | 72% | 100% |
| ODC antizyme inhibitor subfamily | 2 | 652 | 72% | 99% |
| Turtle family | 2 | 1,815 | 72% | 100% |
| Type 2 lipid phosphate phosphatase family | 2 | 603 | 72% | 100% |
| Eukaryotic/archaeal RNase P protein component 2 family | 2 | 206 | 72% | 100% |
| GalK subfamily | 2 | 610 | 72% | 100% |
| BCL9 family | 2 | 2,099 | 72% | 99% |
| Glycosyltransferase 18 family | 2 | 1,100 | 72% | 99% |
| Tomoregulin family | 2 | 541 | 72% | 100% |
| FCHO family | 2 | 1,219 | 72% | 100% |
| TIAM family | 2 | 2,360 | 72% | 99% |
| Meteorin family | 2 | 433 | 72% | 99% |
| VKOR family | 2 | 243 | 72% | 100% |
| TMEM86 family | 2 | 334 | 72% | 100% |
| ERICH6 family | 2 | 974 | 72% | 100% |
| V-ATPase proteolipid subunit family | 2 | 258 | 72% | 100% |
| KDELC family | 2 | 723 | 72% | 99% |
| NIT1/NIT2 family | 2 | 432 | 72% | 100% |
| Raftlin family | 2 | 773 | 72% | 100% |
| CEP63 family | 2 | 936 | 72% | 100% |
| FMN-dependent alpha-hydroxy acid dehydrogenase family | 2 | 516 | 72% | 100% |
| EBP family | 2 | 312 | 72% | 100% |
| TFIIA subunit 1 family | 2 | 611 | 72% | 97% |
| FAK subfamily | 2 | 1,474 | 72% | 99% |
| NOP5/NOP56 family | 2 | 803 | 72% | 98% |
| Cyclin E subfamily | 2 | 582 | 71% | 99% |
| Neurotensin receptor subfamily | 2 | 592 | 71% | 100% |
| Peptidase M2 family | 2 | 1,509 | 71% | 100% |
| FAM193 family | 2 | 1,549 | 71% | 99% |
| DIM1 family | 2 | 208 | 71% | 100% |
| PHOSPHO family | 2 | 363 | 71% | 100% |
| TREX family | 2 | 393 | 71% | 100% |
| FAM177 family | 2 | 265 | 71% | 100% |

| | | | | |
|---|---|---|---|---|
| Astrotactin family | 2 | 1,886 | 71% | 99% |
| Prospero homeodomain family | 2 | 949 | 71% | 99% |
| UPP synthase family | 2 | 447 | 71% | 100% |
| MISP family | 2 | 641 | 71% | 100% |
| Cyclin G subfamily | 2 | 456 | 71% | 100% |
| Themis family | 2 | 916 | 71% | 100% |
| FAM210 family | 2 | 331 | 71% | 100% |
| BICDR family | 2 | 771 | 71% | 100% |
| SMC1 subfamily | 2 | 1,760 | 71% | 99% |
| NSE4 family | 2 | 512 | 71% | 99% |
| Rabaptin family | 2 | 1,020 | 71% | 100% |
| Heavy Metal importer (TC 3,A,1,210) subfamily | 2 | 1,136 | 71% | 99% |
| BCAP29/BCAP31 family | 2 | 347 | 71% | 100% |
| IL-4/IL-13 family | 2 | 213 | 71% | 99% |
| TMEM39 family | 2 | 698 | 71% | 99% |
| Lyase 1 family | 2 | 675 | 71% | 100% |
| SLAIN motif-containing family | 2 | 818 | 71% | 100% |
| FOG (Friend of GATA) family | 2 | 1,535 | 71% | 99% |
| Eukaryotic ribosomal protein eS8 family | 2 | 333 | 71% | 100% |
| TMEM74 family | 2 | 399 | 71% | 100% |
| Podocalyxin family | 2 | 827 | 71% | 100% |
| Prohibitin family | 2 | 406 | 71% | 100% |
| NRK subfamily | 2 | 305 | 71% | 100% |
| Ninjurin family | 2 | 209 | 71% | 100% |
| TMEM87 subfamily | 2 | 789 | 71% | 98% |
| FAM151 family | 2 | 612 | 71% | 100% |
| DDIT4 family | 2 | 302 | 71% | 100% |
| SLC35C subfamily | 2 | 518 | 71% | 100% |
| Gastrokine family | 2 | 260 | 71% | 100% |
| PPP1R15 family | 2 | 985 | 71% | 100% |
| GRXCR1 family | 2 | 382 | 71% | 100% |
| MARCKS family | 2 | 374 | 71% | 100% |
| N4BP1 family | 2 | 1,117 | 71% | 99% |
| PPIL1 subfamily | 2 | 576 | 71% | 100% |
| Bubblegum subfamily | 2 | 986 | 71% | 99% |
| SPATS2 family | 2 | 782 | 71% | 98% |
| ABHD14 family | 2 | 341 | 71% | 99% |
| CRELD family | 2 | 548 | 71% | 99% |
| MCU (TC 1,A,77) family | 2 | 487 | 71% | 99% |
| WSCD family | 2 | 808 | 71% | 99% |
| Fatty acyl-CoA reductase family | 2 | 730 | 71% | 99% |
| Heme oxygenase family | 2 | 428 | 71% | 99% |

| | | | | |
|---|---|---|---|---|
| Suvar4-20 subfamily | 2 | 954 | 71% | 98% |
| DTD family | 2 | 267 | 71% | 100% |
| 2,4-dienoyl-CoA reductase subfamily | 2 | 444 | 71% | 100% |
| TAF4 family | 2 | 1,378 | 71% | 99% |
| Beta-microseminoprotein family | 2 | 179 | 71% | 100% |
| STOP family | 2 | 716 | 71% | 100% |
| Spire family | 2 | 1,040 | 71% | 100% |
| SPTSS family | 2 | 104 | 71% | 100% |
| Glycosyl hydrolase 29 family | 2 | 660 | 71% | 100% |
| Adipolin/erythroferrone family | 2 | 464 | 71% | 100% |
| Arkadia family | 2 | 947 | 71% | 99% |
| Nephronectin family | 2 | 790 | 71% | 100% |
| BRMS1 family | 2 | 402 | 71% | 100% |
| Selenoprotein M/F family | 2 | 219 | 71% | 100% |
| ABHD4/ABHD5 subfamily | 2 | 488 | 71% | 100% |
| Crumbs protein family | 2 | 1,900 | 71% | 100% |
| Class VI-like SAM-binding methyltransferase superfamily | 2 | 341 | 71% | 99% |
| LRRFIP family | 2 | 1,079 | 71% | 98% |
| MON1/SAND family | 2 | 846 | 71% | 100% |
| PAR3 family | 2 | 1,807 | 71% | 98% |
| HEM-1/HEM-2 family | 2 | 1,591 | 71% | 98% |
| CNF-like-inhibitor family | 2 | 311 | 71% | 100% |
| CDC37 family | 2 | 504 | 70% | 100% |
| Adenomatous polyposis coli (APC) family | 2 | 3,627 | 70% | 98% |
| Sesquipedalian family | 2 | 358 | 70% | 99% |
| ATG9 family | 2 | 1,242 | 70% | 99% |
| CDR2 family | 2 | 647 | 70% | 99% |
| NUPR family | 2 | 126 | 70% | 100% |
| V (TC 1,A,1,2) subfamily | 2 | 735 | 70% | 99% |
| APH-1 family | 2 | 367 | 70% | 100% |
| Receptor class 5 subfamily | 2 | 2,643 | 70% | 98% |
| DUOXA family | 2 | 466 | 70% | 97% |
| COLEC10/COLEC11 family | 2 | 385 | 70% | 100% |
| Lin-54 family | 2 | 883 | 70% | 98% |
| PX domain-containing GAP family | 2 | 2,370 | 70% | 99% |
| AFG2 subfamily | 2 | 1,156 | 70% | 100% |
| TspO/BZRP family | 2 | 238 | 70% | 100% |
| Acylphosphatase family | 2 | 139 | 70% | 99% |
| FAM107 family | 2 | 193 | 70% | 100% |
| ELAPOR family | 2 | 1,432 | 70% | 98% |
| Synembryn family | 2 | 737 | 70% | 99% |
| LIX1 family | 2 | 434 | 70% | 99% |

| | | | | |
|---|---|---|---|---|
| Neuritin family | 2 | 215 | 70% | 100% |
| SREBP family | 2 | 1,602 | 70% | 98% |
| WD repeat SEC31 family | 2 | 1,679 | 70% | 98% |
| Urotensin-2 family | 2 | 170 | 70% | 98% |
| Osteocalcin/matrix Gla protein family | 2 | 142 | 70% | 100% |
| CDC42SE/SPEC family | 2 | 114 | 70% | 99% |
| AKR/ZDHHC17 subfamily | 2 | 877 | 70% | 98% |
| SmF/LSm6 subfamily | 2 | 116 | 70% | 100% |
| Sidekick family | 2 | 3,064 | 70% | 99% |
| MRF family | 2 | 1,440 | 70% | 98% |
| Milton family | 2 | 1,304 | 70% | 99% |
| DDAH family | 2 | 398 | 70% | 98% |
| Ataxin-2 family | 2 | 1,667 | 70% | 99% |
| PNRC family | 2 | 325 | 70% | 99% |
| Multi antimicrobial extrusion (MATE) (TC 2,A,66,1) family | 2 | 817 | 70% | 97% |
| Cytochrome c oxidase IV family | 2 | 237 | 70% | 100% |
| ANT/ATPSC lysine N-methyltransferase family | 2 | 326 | 70% | 99% |
| TMEM229 family | 2 | 381 | 70% | 100% |
| Guanylin family | 2 | 158 | 70% | 100% |
| PAIP2 family | 2 | 174 | 70% | 98% |
| SLRP class V subfamily | 2 | 783 | 70% | 100% |
| Feline leukemia virus subgroup C receptor (TC 2,A,1,28,1) family | 2 | 752 | 70% | 98% |
| Refilin family | 2 | 299 | 70% | 99% |
| FAM135 family | 2 | 2,031 | 70% | 97% |
| Cytochrome c oxidase VIII family | 2 | 98 | 70% | 100% |
| STT3 family | 2 | 1,064 | 69% | 98% |
| PIEZO (TC 1,A,75) family | 2 | 3,664 | 69% | 98% |
| Pleiotrophin family | 2 | 216 | 69% | 100% |
| MRL family | 2 | 1,329 | 69% | 99% |
| Beclin family | 2 | 611 | 69% | 100% |
| TAF7 family | 2 | 562 | 69% | 100% |
| Chemokine-like receptor (CMKLR) family | 2 | 504 | 69% | 100% |
| PCP4 family | 2 | 90 | 69% | 100% |
| KIF27 subfamily | 2 | 1,899 | 69% | 100% |
| Hyccin family | 2 | 727 | 69% | 99% |
| SH3PXD2 family | 2 | 1,413 | 69% | 97% |
| ARMET family | 2 | 255 | 69% | 100% |
| RILPL family | 2 | 424 | 69% | 98% |
| HEXIM family | 2 | 445 | 69% | 97% |
| FARP (FMRFamide related peptide) family | 2 | 213 | 69% | 100% |
| FAM114 family | 2 | 736 | 69% | 99% |
| FAM43 family | 2 | 518 | 69% | 100% |

| | | | | |
|---|---|---|---|---|
| TMEM59 family | 2 | 458 | 69% | 100% |
| Nucleoporin NSP1/NUP62 family | 2 | 486 | 69% | 100% |
| RNF12 family | 2 | 901 | 69% | 100% |
| FAM180 family | 2 | 245 | 69% | 100% |
| SERBP1-HABP4 family | 2 | 565 | 69% | 98% |
| Calsequestrin family | 2 | 547 | 69% | 98% |
| Chitinase class II subfamily | 2 | 648 | 69% | 99% |
| PROTOR family | 2 | 520 | 69% | 100% |
| Somatostatin family | 2 | 152 | 69% | 100% |
| NALF family | 2 | 639 | 69% | 97% |
| Eukaryotic RPC7 RNA polymerase subunit family | 2 | 303 | 69% | 100% |
| YIF1 family | 2 | 417 | 69% | 99% |
| GREB1 family | 2 | 2,660 | 69% | 97% |
| Flagellar radial spoke RSP4/6 family | 2 | 984 | 69% | 100% |
| DENND6 family | 2 | 819 | 69% | 99% |
| Su(H) family | 2 | 698 | 69% | 95% |
| Calbindin family | 2 | 365 | 69% | 98% |
| Omega family | 2 | 332 | 69% | 100% |
| TOR1AIP family | 2 | 722 | 69% | 94% |
| TMEM170 family | 2 | 189 | 68% | 100% |
| RUNDC3 family | 2 | 629 | 68% | 98% |
| HEATR5 family | 2 | 2,813 | 68% | 97% |
| Liprin-beta subfamily | 2 | 1,291 | 68% | 95% |
| RENT3 family | 2 | 656 | 68% | 100% |
| Bombesin/neuromedin-B/ranatensin family | 2 | 184 | 68% | 100% |
| SWR1 subfamily | 2 | 4,369 | 68% | 95% |
| Cytospin-A family | 2 | 1,494 | 68% | 97% |
| RRM CPSF6/7 family | 2 | 698 | 68% | 100% |
| NKD family | 2 | 629 | 68% | 97% |
| cGMP subfamily | 2 | 978 | 68% | 99% |
| AK3 subfamily | 2 | 307 | 68% | 98% |
| Fes/fps subfamily | 2 | 1,121 | 68% | 97% |
| NFIL3 subfamily | 2 | 512 | 68% | 97% |
| FAM167 (SEC) family | 2 | 257 | 68% | 100% |
| Tom40 family | 2 | 456 | 68% | 99% |
| NYAP family | 2 | 1,018 | 68% | 100% |
| Fructosamine kinase family | 2 | 421 | 68% | 100% |
| CMC family | 2 | 126 | 68% | 100% |
| XylT subfamily | 2 | 1,242 | 68% | 97% |
| LDOC1 family | 2 | 262 | 68% | 100% |
| Platelet-activating factor acetylhydrolase IB beta/gamma subunits subfamily | 2 | 313 | 68% | 95% |
| Resistin/FIZZ family | 2 | 149 | 68% | 100% |

| | | | | |
|---|---|---|---|---|
| Glycosyl hydrolase 99 family | 2 | 625 | 68% | 97% |
| ASPP family | 2 | 1,506 | 68% | 97% |
| TPRG1 family | 2 | 371 | 68% | 99% |
| Twinfilin subfamily | 2 | 474 | 68% | 99% |
| SEC2 family | 2 | 581 | 68% | 95% |
| STAM family | 2 | 721 | 68% | 98% |
| Glycogenin subfamily | 2 | 576 | 68% | 98% |
| Unkempt family | 2 | 1,008 | 68% | 98% |
| STARD3 family | 2 | 459 | 68% | 96% |
| PSD family | 2 | 1,213 | 68% | 98% |
| ROR subfamily | 2 | 1,269 | 68% | 98% |
| Peptidase M67C family | 2 | 580 | 67% | 96% |
| KIF26 subfamily | 2 | 2,690 | 67% | 97% |
| MCTP family | 2 | 1,265 | 67% | 97% |
| TMEM88 family | 2 | 217 | 67% | 99% |
| SYK/ZAP-70 subfamily | 2 | 844 | 67% | 98% |
| NR0 subfamily | 2 | 489 | 67% | 100% |
| CSN7 subfamily | 2 | 362 | 67% | 98% |
| CSN7/EIF3M family | 2 | 362 | 67% | 98% |
| Mitochondrial Rho GTPase family | 2 | 830 | 67% | 96% |
| GIGYF family | 2 | 1,567 | 67% | 98% |
| PRP38 family | 2 | 576 | 67% | 100% |
| PPase family | 2 | 418 | 67% | 98% |
| AVIT (prokineticin) family | 2 | 157 | 67% | 100% |
| Mediator complex subunit 13 family | 2 | 2,940 | 67% | 97% |
| UGRP subfamily | 2 | 132 | 67% | 100% |
| Prostaglandin G/H synthase family | 2 | 806 | 67% | 98% |
| GnRH family | 2 | 142 | 67% | 100% |
| CSK subfamily | 2 | 641 | 67% | 99% |
| Tie subfamily | 2 | 1,514 | 67% | 94% |
| FNIP family | 2 | 1,524 | 67% | 93% |
| GORASP family | 2 | 596 | 67% | 97% |
| SLA2 family | 2 | 1,406 | 67% | 95% |
| Glutaminyl-peptide cyclotransferase family | 2 | 496 | 67% | 98% |
| CPSF4/YTH1 family | 2 | 299 | 67% | 97% |
| RNF144 subfamily | 2 | 397 | 67% | 96% |
| ABHD16 family | 2 | 685 | 67% | 97% |
| NmU family | 2 | 218 | 67% | 99% |
| Prion family | 2 | 286 | 67% | 100% |
| TMEM151 family | 2 | 689 | 67% | 98% |
| GAREM family | 2 | 1,166 | 67% | 97% |
| ESRP family | 2 | 937 | 67% | 96% |

| | | | | |
|---|---|---|---|---|
| Rheb family | 2 | 244 | 66% | 99% |
| Complex I NDUFA4 subunit family | 2 | 113 | 66% | 100% |
| PACS family | 2 | 1,231 | 66% | 98% |
| OB-RGRP/VPS55 family | 2 | 174 | 66% | 100% |
| Glycosyltransferase 11 family | 2 | 470 | 66% | 95% |
| Type II PI4K subfamily | 2 | 637 | 66% | 97% |
| Eukaryotic PMM family | 2 | 337 | 66% | 97% |
| Cyclin-dependent kinase 5 activator family | 2 | 447 | 66% | 96% |
| SID1 family | 2 | 1,100 | 66% | 94% |
| Non-receptor class 2 subfamily | 2 | 786 | 66% | 99% |
| PPDPF family | 2 | 131 | 66% | 98% |
| WD repeat POC1 family | 2 | 585 | 66% | 95% |
| FAM186 family | 2 | 2,143 | 66% | 100% |
| Receptor class 8 subfamily | 2 | 1,317 | 66% | 95% |
| DnaJ family | 2 | 498 | 66% | 98% |
| DNAJB12/DNAJB14 subfamily | 2 | 498 | 66% | 98% |
| NAB family | 2 | 668 | 66% | 95% |
| SAP30 family | 2 | 266 | 66% | 98% |
| Cappuccino subfamily | 2 | 2,073 | 66% | 100% |
| Vasohibin family | 2 | 475 | 66% | 95% |
| HMG-CoA synthase family | 2 | 678 | 66% | 96% |
| WFIKKN family | 2 | 741 | 66% | 96% |
| VHL family | 2 | 232 | 66% | 99% |
| Neuropeptide B/W family | 2 | 191 | 66% | 99% |
| SHANK family | 2 | 2,641 | 66% | 97% |
| CAND family | 2 | 1,624 | 66% | 94% |
| Multiplexin collagen family | 2 | 2,067 | 66% | 100% |
| CAP family | 2 | 626 | 66% | 98% |
| FUN14 family | 2 | 226 | 66% | 97% |
| EFR3 family | 2 | 1,076 | 66% | 96% |
| V-ATPase C subunit family | 2 | 531 | 66% | 95% |
| GOLGA2 family | 2 | 752 | 66% | 97% |
| TANC family | 2 | 2,525 | 66% | 97% |
| PDE3 subfamily | 2 | 1,477 | 66% | 94% |
| SGF11 family | 2 | 291 | 66% | 95% |
| V-ATPase V0D/AC39 subunit family | 2 | 459 | 65% | 97% |
| Furry protein family | 2 | 3,943 | 65% | 96% |
| FAM63 subfamily | 2 | 713 | 65% | 98% |
| Type IB subfamily | 2 | 1,938 | 65% | 94% |
| PDE7 subfamily | 2 | 609 | 65% | 92% |
| Slowmo family | 2 | 239 | 65% | 98% |
| PRPF40 family | 2 | 1,191 | 65% | 97% |

| | | | | |
|---|---|---|---|---|
| FAM89 family | 2 | 243 | 65% | 95% |
| Endothelin receptor subfamily | 2 | 566 | 65% | 98% |
| ALPK subfamily; Transient receptor (TC 1,A,4) family | 2 | 2,531 | 65% | 95% |
| Zygin family | 2 | 485 | 65% | 96% |
| PRR15 family | 2 | 151 | 65% | 100% |
| RRM NCBP2 family | 2 | 201 | 65% | 99% |
| TMEM255 family | 2 | 439 | 65% | 97% |
| Pterin-4-alpha-carbinolamine dehydratase family | 2 | 152 | 65% | 98% |
| NTE family | 2 | 1,746 | 65% | 94% |
| Galanin family | 2 | 155 | 65% | 97% |
| SGT family | 2 | 400 | 65% | 99% |
| Opioid growth factor receptor family | 2 | 731 | 65% | 99% |
| IPP isomerase type 1 family | 2 | 294 | 65% | 95% |
| FAM219 family | 2 | 248 | 65% | 97% |
| DEGS subfamily | 2 | 418 | 65% | 96% |
| Mediator complex subunit 25 family | 2 | 752 | 65% | 93% |
| SerB family | 2 | 192 | 65% | 91% |
| CWC21 family | 2 | 2,164 | 65% | 100% |
| LARP family | 2 | 1,298 | 65% | 96% |
| SBNO family | 2 | 1,780 | 65% | 93% |
| Vang family | 2 | 674 | 64% | 98% |
| FAM168 family | 2 | 283 | 64% | 95% |
| UPF0220 family | 2 | 203 | 64% | 98% |
| PICALM/SNAP91 family | 2 | 1,004 | 64% | 95% |
| Delta-EF1/ZFH-1 C2H2-type zinc-finger family | 2 | 1,505 | 64% | 90% |
| Non-receptor class 1 subfamily | 2 | 547 | 64% | 93% |
| REXO1/REXO3 family | 2 | 1,220 | 64% | 96% |
| USP20/USP33 subfamily | 2 | 1,194 | 64% | 96% |
| Urea transporter family | 2 | 842 | 64% | 99% |
| 5'-nucleotidase type 3 family | 2 | 628 | 64% | 94% |
| Reprimo family | 2 | 147 | 64% | 98% |
| Clathrin light chain family | 2 | 306 | 64% | 95% |
| SHMT family | 2 | 633 | 64% | 96% |
| COQ10 family | 2 | 311 | 64% | 95% |
| SNURF family | 2 | 123 | 64% | 100% |
| Myosin family; Protein kinase superfamily | 2 | 1,894 | 64% | 94% |
| Nucleobindin family | 2 | 564 | 64% | 94% |
| G(12) subfamily | 2 | 485 | 64% | 94% |
| Mitofusin subfamily | 2 | 958 | 64% | 93% |
| SERF family | 2 | 108 | 64% | 93% |
| Type IB topoisomerase family | 2 | 871 | 64% | 96% |
| Eukaryotic ribosomal protein P1/P2 family | 2 | 146 | 64% | 91% |

| | | | | |
|---|---|---|---|---|
| TCAF family | 2 | 1,172 | 64% | 95% |
| NRAMP family | 2 | 711 | 64% | 97% |
| Cytochrome c oxidase subunit 6A family | 2 | 131 | 64% | 97% |
| LSM14 family | 2 | 539 | 64% | 95% |
| Troponin C family | 2 | 204 | 64% | 98% |
| Class-6 subfamily | 2 | 630 | 64% | 95% |
| NK-3 homeobox family | 2 | 360 | 63% | 96% |
| Enhancer of polycomb family | 2 | 1,043 | 63% | 93% |
| ALKAL family | 2 | 178 | 63% | 95% |
| KappaB-Ras subfamily | 2 | 242 | 63% | 97% |
| PNP/UDP phosphorylase family | 2 | 394 | 63% | 94% |
| Mediator complex subunit 12 family | 2 | 2,710 | 63% | 94% |
| Hic subfamily | 2 | 844 | 63% | 95% |
| OST3/OST6 family | 2 | 427 | 63% | 94% |
| Alanine aminotransferase subfamily | 2 | 637 | 63% | 92% |
| ERF4 family | 2 | 190 | 63% | 98% |
| Gastrin/cholecystokinin family | 2 | 135 | 63% | 98% |
| TMEM9 family | 2 | 238 | 62% | 97% |
| HNF1 homeobox family | 2 | 742 | 62% | 92% |
| EEIG family | 2 | 463 | 62% | 94% |
| HMG-CoA lyase family | 2 | 432 | 62% | 93% |
| Glycosyltransferase 49 family; Glycosyltransferase 8 family | 2 | 916 | 62% | 93% |
| Tetrahydrofolate dehydrogenase/cyclohydrolase family | 2 | 432 | 62% | 93% |
| RasD family | 2 | 339 | 62% | 90% |
| EROs family | 2 | 579 | 62% | 91% |
| CAMTA family | 2 | 1,778 | 62% | 92% |
| Akirin family | 2 | 244 | 62% | 96% |
| CTCF zinc-finger protein family | 2 | 857 | 62% | 95% |
| Glyceraldehyde-3-phosphate dehydrogenase family | 2 | 458 | 62% | 96% |
| Pro/parathymosin family | 2 | 131 | 62% | 100% |
| HAPSTR1 family | 2 | 337 | 61% | 95% |
| NR5 subfamily | 2 | 616 | 61% | 90% |
| Tetrahydrofolate dehydrogenase/cyclohydrolase family; Formate--tetrahydrofolate ligase family | 2 | 1,176 | 61% | 89% |
| EAF family | 2 | 324 | 61% | 94% |
| ZBTB18 subfamily | 2 | 579 | 61% | 95% |
| AGR family | 2 | 209 | 61% | 91% |
| UPF0400 (RTT103) family | 2 | 391 | 61% | 89% |
| GOLPH3/VPS74 family | 2 | 357 | 61% | 94% |
| FAM78 family | 2 | 333 | 61% | 92% |
| WD repeat ARPC1 family | 2 | 454 | 61% | 92% |
| ARPC5 family | 2 | 186 | 61% | 95% |

| | | | | |
|---|---|---|---|---|
| Pyruvate kinase family | 2 | 675 | 61% | 93% |
| MTSS family | 2 | 917 | 61% | 95% |
| PDE8 subfamily | 2 | 1,045 | 61% | 90% |
| 3-oxoacid CoA-transferase family | 2 | 632 | 61% | 88% |
| BRE1 family | 2 | 1,204 | 61% | 91% |
| GPSM family | 2 | 828 | 61% | 91% |
| ENOX family | 2 | 763 | 61% | 92% |
| BicD family | 2 | 1,094 | 61% | 92% |
| FAM24 family | 2 | 121 | 61% | 90% |
| SMARCC family | 2 | 1,405 | 61% | 87% |
| RNF19 subfamily | 2 | 951 | 61% | 89% |
| Round spermatid basic protein 1 family | 2 | 998 | 61% | 91% |
| Protein sulfotransferase family | 2 | 452 | 61% | 94% |
| CRK family | 2 | 367 | 60% | 93% |
| RUTBC family | 2 | 1,302 | 60% | 91% |
| NK-1 homeobox family | 2 | 458 | 60% | 88% |
| BZW family | 2 | 506 | 60% | 94% |
| PDS5 family | 2 | 1,678 | 60% | 91% |
| Inorganic phosphate transporter (PiT) (TC 2,A,20) family | 2 | 800 | 60% | 92% |
| PHYHIP family | 2 | 424 | 60% | 88% |
| NECAP family | 2 | 323 | 60% | 96% |
| Phosphoenolpyruvate carboxykinase [GTP] family | 2 | 757 | 60% | 90% |
| Glutaminase family | 2 | 762 | 60% | 93% |
| Elbow/Noc family | 2 | 741 | 60% | 94% |
| KCTD3 family | 2 | 909 | 60% | 90% |
| Receptor class 4 subfamily | 2 | 895 | 60% | 93% |
| DDX21/DDX50 subfamily | 2 | 905 | 60% | 91% |
| Glycosyltransferase 3 family | 2 | 854 | 59% | 90% |
| YY transcription factor family | 2 | 466 | 59% | 92% |
| SEC15 family | 2 | 954 | 59% | 92% |
| Lin-28 family | 2 | 271 | 59% | 94% |
| NAD-dependent glycerol-3-phosphate dehydrogenase family | 2 | 411 | 59% | 92% |
| Even-skipped homeobox family | 2 | 518 | 59% | 88% |
| TIKI family | 2 | 599 | 59% | 90% |
| CAMK2N family | 2 | 92 | 59% | 91% |
| CTP synthase family | 2 | 689 | 59% | 89% |
| SMAUG family | 2 | 826 | 58% | 89% |
| NIP3 family | 2 | 241 | 58% | 93% |
| HMGA family | 2 | 126 | 58% | 97% |
| RAB6IP1 family | 2 | 1,493 | 58% | 89% |
| NudE family | 2 | 396 | 58% | 88% |

| | | | | |
|---|---|---|---|---|
| UPF0046 family | 2 | 361 | 58% | 94% |
| PMEPA1 family | 2 | 345 | 58% | 86% |
| Cyclin L subfamily | 2 | 606 | 58% | 90% |
| Histone H4 family | 2 | 116 | 58% | 97% |
| RimK family | 2 | 448 | 58% | 89% |
| STRIP family | 2 | 962 | 58% | 89% |
| FAM200 family | 2 | 707 | 57% | 92% |
| ALDH1L subfamily | 2 | 1,046 | 57% | 92% |
| GART family; Aldehyde dehydrogenase family | 2 | 1,046 | 57% | 92% |
| RAD23 family | 2 | 442 | 57% | 91% |
| Engrailed homeobox family | 2 | 414 | 57% | 88% |
| FAM163 family | 2 | 190 | 57% | 81% |
| GCN5 subfamily | 2 | 952 | 57% | 88% |
| SOSS-B family | 2 | 236 | 57% | 92% |
| RRM U1 A/B" family | 2 | 286 | 56% | 89% |
| Proboscipedia subfamily | 2 | 411 | 56% | 83% |
| VAMP-associated protein (VAP) (TC 9,B,17) family | 2 | 275 | 56% | 87% |
| DDI1 family | 2 | 444 | 56% | 89% |
| Msh homeobox family | 2 | 317 | 56% | 84% |
| CHIC family | 2 | 216 | 56% | 93% |
| CLASP family | 2 | 1,572 | 56% | 88% |
| PI transfer class I subfamily | 2 | 300 | 55% | 89% |
| APS kinase family; Sulfate adenylyltransferase family | 2 | 685 | 55% | 85% |
| VIP1 subfamily | 2 | 1,478 | 55% | 83% |
| FBPase class 1 family | 2 | 372 | 55% | 87% |
| ABL subfamily | 2 | 1,265 | 55% | 83% |
| Neutral ceramidase family | 2 | 517 | 55% | 72% |
| TMEM120 family | 2 | 371 | 54% | 86% |
| Peptidase A22A family | 2 | 494 | 54% | 85% |
| Dystrobrevin subfamily | 2 | 739 | 54% | 82% |
| Dystrophin family | 2 | 739 | 54% | 82% |
| 5'-AMP-activated protein kinase beta subunit family | 2 | 292 | 54% | 82% |
| INSIG family | 2 | 270 | 54% | 91% |
| Type II topoisomerase family | 2 | 1,697 | 54% | 83% |
| RHPN family | 2 | 723 | 53% | 80% |
| Eukaryotic ribosomal protein eL22 family | 2 | 133 | 53% | 85% |
| COPG family | 2 | 924 | 53% | 87% |
| Mo25 family | 2 | 358 | 53% | 85% |
| RANBP9/10 family | 2 | 711 | 53% | 84% |
| NKAP family | 2 | 429 | 53% | 83% |
| CD200R family | 2 | 324 | 52% | 80% |

| | | | | |
|---|---|---|---|---|
| UBALD family | 2 | 178 | 52% | 87% |
| Adenylosuccinate synthetase family | 2 | 476 | 52% | 79% |
| Selenophosphate synthase 1 family | 2 | 437 | 52% | 84% |
| DDX5/DBP2 subfamily | 2 | 698 | 52% | 82% |
| LDB family | 2 | 407 | 52% | 82% |
| Musashi family | 2 | 358 | 52% | 82% |
| CDK2AP family | 2 | 125 | 52% | 84% |
| FAM91 family | 2 | 579 | 52% | 77% |
| V-ATPase E subunit family | 2 | 234 | 52% | 76% |
| CYRI family | 2 | 334 | 52% | 86% |
| Cytochrome c oxidase VIIb family | 2 | 83 | 52% | 84% |
| EZ subfamily | 2 | 767 | 51% | 81% |
| Semenogelin family | 2 | 536 | 51% | 88% |
| FAM76 family | 2 | 331 | 51% | 86% |
| CDS family | 2 | 463 | 51% | 78% |
| Arylamine N-acetyltransferase family | 2 | 295 | 51% | 87% |
| Vasopressin/oxytocin family | 2 | 146 | 51% | 78% |
| DNA photolyase class-1 family | 2 | 591 | 50% | 83% |
| ASF1 family | 2 | 203 | 50% | 80% |
| MIC subfamily | 2 | 382 | 50% | 86% |
| LEM family | 2 | 572 | 50% | 68% |
| ST7 family | 2 | 576 | 50% | 83% |
| GuaC type 1 subfamily | 2 | 344 | 50% | 82% |
| B (Shab) (TC 1,A,1,2) subfamily | 2 | 862 | 49% | 73% |
| Odd C2H2-type zinc-finger protein family | 2 | 281 | 49% | 75% |
| FAM133 family | 2 | 240 | 48% | 83% |
| GMF subfamily | 2 | 137 | 48% | 80% |
| NMT family | 2 | 477 | 48% | 79% |
| FAM50 family | 2 | 317 | 48% | 77% |
| Ribonucleoside diphosphate reductase small chain family | 2 | 351 | 47% | 81% |
| Eukaryotic ribosomal protein eL43 family | 2 | 87 | 47% | 82% |
| Endosulfine family | 2 | 108 | 46% | 79% |
| Clathrin heavy chain family | 2 | 1,534 | 46% | 76% |
| Amelogenin family | 2 | 183 | 46% | 84% |
| Fibrillarin family | 2 | 301 | 46% | 82% |
| CT47 family | 2 | 284 | 46% | 79% |
| CFAP144 family | 2 | 122 | 45% | 75% |
| GSK-3-binding protein family | 2 | 232 | 45% | 76% |
| NAC-beta family | 2 | 163 | 45% | 74% |
| IGBP1/TAP42 family | 2 | 303 | 45% | 79% |
| SEC23 subfamily | 2 | 681 | 44% | 78% |
| ZCCHC12 family | 2 | 357 | 44% | 73% |

| | | | | |
|---|---|---|---|---|
| AdoMet synthase family | 2 | 344 | 44% | 72% |
| ISWI subfamily | 2 | 914 | 43% | 78% |
| Tdpoz family | 2 | 331 | 43% | 73% |
| Importin beta-2 subfamily | 2 | 751 | 42% | 71% |
| Phosphoglycerate kinase family | 2 | 347 | 42% | 76% |
| WD repeat WDR5/wds family | 2 | 272 | 41% | 74% |
| V-ATPase e1/e2 subunit family | 2 | 65 | 40% | 69% |
| DPH3 family | 2 | 64 | 40% | 69% |
| TMEM185 family | 2 | 280 | 40% | 69% |
| GSK-3 subfamily | 2 | 361 | 40% | 66% |
| ATPase g subunit family | 2 | 81 | 40% | 76% |
| Phosphatase 2A regulatory subunit A family | 2 | 472 | 40% | 64% |
| CKS family | 2 | 62 | 39% | 82% |
| RRP7 family | 2 | 148 | 39% | 56% |
| Glycophorin-A family | 2 | 65 | 38% | 79% |
| UTP14 family | 2 | 585 | 38% | 69% |
| HPF1 family | 2 | 258 | 37% | 64% |
| CYFIP family | 2 | 922 | 36% | 63% |
| FAM209 family | 2 | 120 | 35% | 59% |
| RsmH family | 2 | 221 | 34% | 52% |
| SAR1 family | 2 | 135 | 34% | 63% |
| HERV class-I H env subfamily | 2 | 381 | 33% | 65% |
| USP12/USP46 subfamily | 2 | 246 | 33% | 60% |
| ARP1 subfamily | 2 | 248 | 33% | 58% |
| S100-fused protein family; S-100 family | 2 | 1,722 | 33% | 99% |
| TMA7 family | 2 | 42 | 33% | 73% |
| WD repeat RBAP46/RBAP48/MSI1 family | 2 | 278 | 33% | 60% |
| Rod/cone cGMP-PDE gamma subunit family | 2 | 55 | 32% | 64% |
| DECD subfamily | 2 | 251 | 29% | 54% |
| RAMP4 family | 2 | 38 | 29% | 46% |
| Eukaryotic ATPase epsilon family | 2 | 29 | 28% | 56% |
| ERF3 subfamily | 2 | 320 | 28% | 49% |
| Dihydrofolate reductase family | 2 | 106 | 28% | 64% |
| ARP3 subfamily | 2 | 221 | 26% | 61% |
| Chromokinesin subfamily | 2 | 640 | 26% | 50% |
| Eukaryotic ribosomal protein eS10 family | 2 | 84 | 25% | 58% |
| SnRNP SmB/SmN family | 2 | 116 | 24% | 50% |
| ZFX/ZFY subfamily | 2 | 384 | 24% | 47% |
| DDX3/DED1 subfamily | 2 | 314 | 24% | 42% |
| TAF1 family | 2 | 864 | 23% | 44% |
| Eukaryotic ribosomal protein eS26 family | 2 | 53 | 23% | 46% |
| SecY/SEC61-alpha family | 2 | 214 | 22% | 42% |

| | | | | |
|---|---|---|---|---|
| PDPK1 subfamily | 2 | 200 | 21% | 30% |
| MCTS1 family | 2 | 71 | 20% | 38% |
| SUI1 family | 2 | 44 | 19% | 38% |
| Glu/Leu/Phe/Val dehydrogenases family | 2 | 209 | 19% | 37% |
| SPAG11 family | 2 | 41 | 18% | 32% |
| TTC30/dfy-1/fleer family | 2 | 238 | 18% | 36% |
| Ribulose-phosphate 3-epimerase family | 2 | 81 | 18% | 39% |
| TYW1 family | 2 | 229 | 16% | 32% |
| FAM153 family | 2 | 111 | 16% | 35% |
| MOZART2 family | 2 | 50 | 16% | 33% |
| ALG10 glucosyltransferase family | 2 | 149 | 16% | 30% |
| Eukaryotic ribosomal protein eS27 family | 2 | 26 | 15% | 27% |
| FCGR1 family | 2 | 100 | 15% | 27% |
| NHER family | 2 | 132 | 14% | 24% |
| SFTPA family | 2 | 67 | 14% | 30% |
| UQCRH/QCR6 family | 2 | 23 | 13% | 29% |
| Gonadal family | 2 | 54 | 12% | 28% |
| Rieske iron-sulfur protein family | 2 | 47 | 8% | 19% |
| DDX19/DBP5 subfamily | 2 | 80 | 8% | 16% |
| Complex I NDUFC2 subunit family | 2 | 18 | 8% | 15% |
| EIF2G subfamily | 2 | 69 | 7% | 17% |
| TMEM183 family | 2 | 54 | 7% | 16% |
| FAM21 family | 2 | 173 | 6% | 15% |
| KCNJ12 subfamily | 2 | 55 | 6% | 15% |
| EOLA family | 2 | 19 | 6% | 14% |
| DENND10 family | 2 | 40 | 6% | 12% |
| FAM106 family | 2 | 18 | 5% | 14% |
| Intercrine gamma family | 2 | 11 | 5% | 11% |
| Eukaryotic ribosomal protein eL42 family | 2 | 10 | 5% | 10% |
| RAM family | 2 | 11 | 5% | 11% |
| GTF2H2 family | 2 | 34 | 4% | 10% |
| EIF-1A family | 2 | 11 | 4% | 8% |
| FAM223 family | 2 | 9 | 4% | 9% |
| Mago nashi family | 2 | 6 | 2% | 5% |
| GATD3 family | 2 | 8 | 1% | 4% |
| AMPK subfamily | 2 | 11 | 1% | 2% |
| EIF-3 subunit C family | 2 | 5 | 0% | 1% |
| CCZ1 family | 2 | 0 | 0% | 0% |
| Golgi pH regulator (TC 1,A,38) family | 2 | 0 | 0% | 0% |

**Table S7.** Analysis of human protein families, with more than 10 members, by CrUP profiling

| Family name | Proteins per Family (number) | CrUPs per Family (number) | Density of CrUPs per Family (%) | Family Coverage by CrUPs (%) |
|---|---|---|---|---|
| Type 2 subfamily | 14 | 7,741 | 75% | 100% |
| Type II cytokine receptor family | 11 | 3,732 | 74% | 100% |
| Type I cytokine receptor family | 32 | 14,355 | 74% | 100% |
| Tumor necrosis factor family | 18 | 3,301 | 73% | 100% |
| BPI/LBP/Plunc superfamily | 14 | 4,383 | 75% | 100% |
| SLC26A/SulP transporter (TC 2,A,53) family | 11 | 5,872 | 73% | 100% |
| IL-1 family | 11 | 1,615 | 74% | 100% |
| Class-I aminoacyl-tRNA synthetase family | 19 | 11,320 | 74% | 100% |
| Acyl-CoA dehydrogenase family | 11 | 4,459 | 73% | 100% |
| XK family | 10 | 3,436 | 71% | 100% |
| Interleukin-1 receptor family | 10 | 4,039 | 72% | 100% |
| Complex I LYR family | 11 | 856 | 72% | 100% |
| Phospholipase A2 family | 12 | 1,822 | 71% | 100% |
| Ankyrin SOCS box (ASB) family | 18 | 5,211 | 71% | 100% |
| Cation diffusion facilitator (CDF) transporter (TC 2,A,4) family | 10 | 3,438 | 73% | 100% |
| SLC30A subfamily | 10 | 3,438 | 73% | 100% |
| Small leucine-rich proteoglycan (SLRP) family | 17 | 5,173 | 70% | 100% |
| Metallo-beta-lactamase superfamily | 11 | 3,532 | 74% | 100% |
| Peptidase T1B family | 11 | 1,949 | 71% | 100% |
| Protein disulfide isomerase family | 11 | 4,204 | 72% | 100% |
| Glycosyltransferase 29 family | 20 | 5,375 | 71% | 99% |
| Glycosyltransferase 31 family | 23 | 6,115 | 71% | 99% |
| Nectin family | 10 | 3,274 | 71% | 99% |
| Tubulin--tyrosine ligase family | 10 | 5,743 | 73% | 99% |
| Monocarboxylate porter (TC 2,A,1,13) family | 14 | 4,824 | 69% | 99% |
| Bcl-2 family | 16 | 2,799 | 72% | 99% |
| Non-receptor class subfamily | 11 | 7,198 | 71% | 99% |
| Alpha-carbonic anhydrase family | 16 | 3,320 | 69% | 99% |
| Membrane-bound acyltransferase family | 11 | 3,943 | 73% | 99% |
| Peptidase S8 family | 10 | 6,827 | 72% | 99% |
| Peptidase C2 family | 15 | 8,374 | 70% | 99% |
| Shisa family | 11 | 2,379 | 70% | 99% |
| Helicase family | 23 | 20,260 | 72% | 99% |
| Peptidase M14 family | 23 | 10,659 | 69% | 99% |

| | | | | |
|---|---|---|---|---|
| MS4A family | 16 | 2,988 | 71% | 99% |
| Lipase family | 22 | 7,182 | 69% | 99% |
| CRISP family | 12 | 2,503 | 67% | 99% |
| Integrin alpha chain family | 18 | 14,203 | 70% | 98% |
| ZIP transporter (TC 2,A,5) family | 14 | 4,819 | 70% | 98% |
| Tetraspanin (TM4SF) family | 31 | 5,524 | 70% | 98% |
| Peptidase C1 family | 15 | 3,900 | 70% | 98% |
| PA-phosphatase related phosphoesterase family | 12 | 3,044 | 68% | 98% |
| Beta/gamma-crystallin family | 15 | 6,168 | 71% | 98% |
| Class-I pyridoxal-phosphate-dependent aminotransferase family | 11 | 3,595 | 70% | 98% |
| Class V-like SAM-binding methyltransferase superfamily | 46 | 40,268 | 69% | 98% |
| Kallikrein subfamily | 15 | 2,549 | 65% | 98% |
| LDLR family | 15 | 16,397 | 67% | 98% |
| Semaphorin family | 20 | 11,747 | 69% | 98% |
| NLRP family | 17 | 13,083 | 71% | 98% |
| Pancreatic ribonuclease family | 13 | 1,467 | 68% | 98% |
| PP2C family | 17 | 5,744 | 70% | 98% |
| 1-acyl-sn-glycerol-3-phosphate acyltransferase family | 12 | 3,434 | 69% | 98% |
| Histone-lysine methyltransferase family | 21 | 24,426 | 69% | 98% |
| Arrestin family | 10 | 2,696 | 67% | 98% |
| Adhesion G-protein coupled receptor (ADGR) subfamily | 26 | 23,105 | 70% | 98% |
| RBR family | 12 | 5,191 | 70% | 97% |
| Dynein heavy chain family | 16 | 46,996 | 69% | 97% |
| Sodium:solute symporter (SSF) (TC 2,A,21) family | 12 | 5,058 | 65% | 97% |
| Peptidase M10A family | 23 | 7,912 | 67% | 97% |
| ABCC family | 12 | 12,023 | 68% | 97% |
| Non-receptor class myotubularin subfamily | 15 | 8,999 | 67% | 97% |
| NEK Ser/Thr protein kinase family | 11 | 5,459 | 69% | 97% |
| NIMA subfamily | 11 | 5,459 | 69% | 97% |
| Sorting nexin family | 31 | 10,107 | 69% | 97% |
| G-protein coupled receptor 2 family | 49 | 37,118 | 69% | 97% |
| Anoctamin family | 10 | 6,363 | 68% | 97% |
| Conjugate transporter (TC 3,A,1,208) subfamily | 11 | 10,914 | 67% | 97% |
| Major facilitator (TC 2,A,1) superfamily | 23 | 8,659 | 68% | 97% |
| Organic cation transporter (TC 2,A,1,19) family | 23 | 8,659 | 68% | 97% |
| ATF subfamily | 13 | 3,516 | 67% | 97% |
| BZIP family | 55 | 13,492 | 66% | 97% |

| | | | | |
|---|---|---|---|---|
| Sulfatase family | 17 | 6,630 | 66% | 97% |
| Class-II aminoacyl-tRNA synthetase family | 19 | 8,081 | 70% | 97% |
| ABC transporter superfamily | 48 | 42,703 | 67% | 97% |
| Non-receptor class dual specificity subfamily | 27 | 6,115 | 67% | 97% |
| ABCA family | 11 | 14,144 | 66% | 97% |
| PMP-22/EMP/MP20 family | 19 | 2,968 | 67% | 97% |
| DOCK family | 11 | 14,620 | 66% | 96% |
| Short-chain dehydrogenases/reductases (SDR) family | 54 | 11,644 | 68% | 96% |
| AB hydrolase superfamily | 51 | 13,908 | 68% | 96% |
| Amino acid/polyamine transporter 2 family | 16 | 5,628 | 67% | 96% |
| Insulin receptor subfamily | 12 | 9,530 | 67% | 96% |
| Two pore domain potassium channel (TC 1,A,1,8) family | 15 | 3,782 | 66% | 96% |
| TKL Ser/Thr protein kinase family | 34 | 16,565 | 67% | 96% |
| Serpin family | 38 | 10,406 | 68% | 96% |
| Peptidase C14A family | 13 | 3,448 | 69% | 96% |
| PI3/PI4-kinase family | 20 | 24,378 | 71% | 96% |
| MAP kinase kinase kinase subfamily | 18 | 11,816 | 67% | 96% |
| Toll-like receptor family | 11 | 6,615 | 69% | 96% |
| Formin homology family | 13 | 10,335 | 65% | 96% |
| Plexin family | 11 | 11,706 | 65% | 96% |
| Syntaxin family | 16 | 3,111 | 67% | 96% |
| MAGUK family | 17 | 10,649 | 66% | 96% |
| Fibrillar collagen family | 11 | 9,840 | 55% | 96% |
| Sodium:neurotransmitter symporter (SNF) (TC 2,A,22) family | 19 | 7,926 | 64% | 96% |
| Aldehyde dehydrogenase family | 16 | 5,335 | 64% | 96% |
| Tyr protein kinase family | 90 | 55,715 | 65% | 96% |
| S-100 family | 23 | 2,104 | 68% | 96% |
| Splicing factor SR family | 26 | 7,161 | 62% | 96% |
| G-protein coupled receptor 3 family | 22 | 13,279 | 66% | 96% |
| ARTD/PARP family | 17 | 10,307 | 68% | 96% |
| Metallo-dependent hydrolases superfamily | 24 | 9,132 | 66% | 96% |
| Protein-tyrosine phosphatase family | 93 | 47,978 | 66% | 96% |
| Major facilitator superfamily | 81 | 28,347 | 67% | 96% |
| Connexin family | 21 | 4,528 | 64% | 96% |
| Cyclin family | 32 | 9,706 | 67% | 96% |
| Heparin-binding growth factors family | 22 | 3,120 | 64% | 96% |
| Glycosyltransferase 2 family | 26 | 9,448 | 65% | 95% |
| OSBP family | 12 | 6,320 | 64% | 95% |

| | | | | |
|---|---|---|---|---|
| Ser/Thr protein kinase family | 67 | 34,552 | 67% | 95% |
| Class I-like SAM-binding methyltransferase superfamily | 62 | 19,805 | 69% | 95% |
| ADIPOR family | 11 | 2,387 | 66% | 95% |
| SnRNP Sm proteins family | 15 | 1,205 | 68% | 95% |
| Sp1 C2H2-type zinc-finger protein family | 19 | 5,524 | 63% | 95% |
| SNF7 family | 12 | 1,831 | 66% | 95% |
| AAA ATPase family | 35 | 19,409 | 67% | 95% |
| GalNAc-T subfamily | 20 | 7,675 | 63% | 95% |
| Fatty-acid binding protein (FABP) family | 17 | 1,395 | 62% | 95% |
| Acetyltransferase family | 19 | 4,753 | 66% | 94% |
| ABCB family | 11 | 6,779 | 65% | 94% |
| Monovalent cation:proton antiporter 1 (CPA1) transporter (TC 2,A,36) family | 14 | 6,785 | 65% | 94% |
| Kinesin family | 45 | 36,421 | 65% | 94% |
| Cullin family | 10 | 6,753 | 67% | 94% |
| Synaptobrevin family | 11 | 1,263 | 65% | 94% |
| HAD-like hydrolase superfamily | 18 | 3,824 | 65% | 94% |
| Amino acid-polyamine-organocation (APC) superfamily | 15 | 4,739 | 61% | 94% |
| CAMK Ser/Thr protein kinase family | 75 | 59,158 | 63% | 94% |
| Adenylyl cyclase class-4/guanylyl cyclase family | 19 | 13,172 | 65% | 94% |
| Krueppel C2H2-type zinc-finger protein family | 543 | 170,767 | 50% | 94% |
| ATP-dependent AMP-binding enzyme family | 27 | 11,581 | 65% | 93% |
| E2F/DP family | 11 | 3,454 | 64% | 93% |
| Peptidase S1 family | 119 | 33,097 | 63% | 93% |
| Type IV subfamily | 14 | 10,641 | 61% | 93% |
| Ephrin receptor subfamily | 14 | 7,941 | 57% | 93% |
| DEAH subfamily | 22 | 16,111 | 65% | 93% |
| CD225/Dispanin family | 16 | 1,876 | 62% | 93% |
| Protein kinase superfamily | 491 | 265,955 | 63% | 93% |
| Cyclic nucleotide phosphodiesterase family | 21 | 10,101 | 61% | 93% |
| Wnt family | 19 | 4,101 | 59% | 93% |
| Importin beta family | 11 | 6,978 | 63% | 93% |
| TRAFAC class TrmE-Era-EngA-EngB-Septin-like GTPase superfamily | 24 | 5,749 | 60% | 93% |
| Cystatin family | 15 | 1,301 | 63% | 92% |
| Peptidase M1 family | 13 | 7,382 | 67% | 92% |
| DHHC palmitoyltransferase family | 24 | 6,281 | 64% | 92% |
| Transient receptor (TC 1,A,4) family | 22 | 13,331 | 62% | 92% |
| Histone deacetylase family | 11 | 5,060 | 62% | 92% |
| Eukaryotic diacylglycerol kinase family | 10 | 5,955 | 62% | 92% |

| | | | | |
|---|---|---|---|---|
| Methyltransferase superfamily | 43 | 10,351 | 66% | 92% |
| Abd-B homeobox family | 16 | 3,080 | 61% | 92% |
| SNF2/RAD54 helicase family | 32 | 33,819 | 62% | 92% |
| ETS family | 29 | 7,535 | 62% | 92% |
| Claudin family | 24 | 3,411 | 63% | 92% |
| GLI C2H2-type zinc-finger protein family | 12 | 5,566 | 60% | 92% |
| SRC subfamily | 10 | 2,761 | 53% | 91% |
| Mitochondrial carrier (TC 2,A,29) family | 53 | 11,063 | 61% | 91% |
| Sugar transporter (TC 2,A,1,1) family | 14 | 4,676 | 63% | 91% |
| STE Ser/Thr protein kinase family | 56 | 27,858 | 60% | 91% |
| Anion exchanger (TC 2,A,31) family | 10 | 6,330 | 58% | 91% |
| DEAD box helicase family | 60 | 31,097 | 63% | 91% |
| TGFB receptor subfamily | 12 | 4,003 | 59% | 91% |
| PKC subfamily | 12 | 5,076 | 57% | 90% |
| Bicoid subfamily | 12 | 2,268 | 58% | 90% |
| Ov-serpin subfamily | 13 | 3,019 | 60% | 90% |
| Glucose transporter subfamily | 13 | 4,186 | 62% | 90% |
| TRAFAC class myosin-kinesin ATPase superfamily | 85 | 74,902 | 59% | 90% |
| Calycin superfamily | 38 | 3,858 | 62% | 90% |
| Immunoglobulin superfamily | 130 | 44,970 | 59% | 90% |
| G protein gamma family | 14 | 532 | 54% | 90% |
| S6 kinase subfamily | 10 | 3,927 | 55% | 90% |
| Intercrine beta (chemokine CC) family | 26 | 1,654 | 61% | 90% |
| TRAFAC class dynamin-like GTPase superfamily | 30 | 12,299 | 59% | 90% |
| PMG family | 10 | 1,078 | 62% | 89% |
| Ubiquitin-conjugating enzyme family | 43 | 9,251 | 63% | 89% |
| G-protein coupled receptor Fz/Smo family | 11 | 3,896 | 56% | 89% |
| Insulin family | 10 | 991 | 61% | 89% |
| Antp homeobox family | 26 | 4,316 | 57% | 89% |
| BTN/MOG family | 19 | 4,634 | 56% | 89% |
| Nudix hydrolase family | 24 | 3,943 | 65% | 89% |
| TPT transporter family | 10 | 2,307 | 64% | 89% |
| Intercrine alpha (chemokine CxC) family | 17 | 1,140 | 57% | 89% |
| Beta-defensin family | 35 | 1,860 | 63% | 89% |
| Zinc-containing alcohol dehydrogenase family | 16 | 3,582 | 60% | 89% |
| TGF-beta family | 37 | 8,819 | 61% | 89% |
| NDK family | 10 | 1,543 | 62% | 89% |
| NR1 subfamily | 19 | 5,222 | 58% | 89% |
| G-protein coupled receptor T2R family | 27 | 4,615 | 55% | 89% |
| Nuclear hormone receptor family | 48 | 14,346 | 58% | 88% |
| Histone H1/H5 family | 11 | 1,397 | 55% | 88% |

| | | | | |
|---|---|---|---|---|
| G-protein coupled receptor 1 family | 724 | 137,161 | 54% | 88% |
| Acetylcholine receptor (TC 1,A,9,1) subfamily | 16 | 4,609 | 58% | 88% |
| Arf family | 28 | 3,509 | 58% | 88% |
| Septin GTPase family | 13 | 2,941 | 53% | 88% |
| Lipocalin family | 21 | 2,463 | 61% | 87% |
| STE20 subfamily | 29 | 13,008 | 56% | 87% |
| Cation transport ATPase (P-type) (TC 3,A,3) family | 36 | 23,404 | 55% | 87% |
| Glutamate-gated ion channel (TC 1,A,10,1) family | 18 | 10,840 | 57% | 87% |
| AGC Ser/Thr protein kinase family | 58 | 26,169 | 56% | 87% |
| TRIM/RBCC family | 65 | 20,811 | 60% | 87% |
| TCP-1 chaperonin family | 14 | 4,978 | 62% | 87% |
| Synaptotagmin family | 19 | 4,974 | 58% | 87% |
| Dynamin/Fzo/YdjA family | 14 | 5,488 | 56% | 87% |
| DNA mismatch repair MutL/HexB family | 10 | 3,273 | 59% | 87% |
| Potassium channel family | 44 | 18,153 | 55% | 87% |
| GB1 subfamily | 11 | 3,384 | 51% | 86% |
| GB1/RHD3 GTPase family | 11 | 3,384 | 51% | 86% |
| Adaptor complexes large subunit family | 11 | 6,193 | 58% | 86% |
| Myosin family | 37 | 35,246 | 54% | 86% |
| SNF1 subfamily | 14 | 6,006 | 57% | 86% |
| Paired homeobox family | 44 | 7,803 | 57% | 86% |
| CMGC Ser/Thr protein kinase family | 62 | 19,831 | 56% | 86% |
| MHC class II family | 14 | 1,885 | 52% | 85% |
| MAP kinase subfamily | 14 | 3,610 | 54% | 85% |
| Calcium channel alpha-1 subunit (TC 1,A,1,11) family | 12 | 12,775 | 53% | 85% |
| SIGLEC (sialic acid binding Ig-like lectin) family | 15 | 4,943 | 54% | 85% |
| Type-B carboxylesterase/lipase family | 15 | 6,665 | 55% | 84% |
| TRAFAC class translation factor GTPase superfamily | 22 | 9,502 | 61% | 84% |
| CDC2/CDKX subfamily | 29 | 8,521 | 54% | 84% |
| CaMK subfamily | 13 | 3,473 | 50% | 84% |
| Ligand-gated ion channel (TC 1,A,9) family | 46 | 11,787 | 53% | 84% |
| Cytochrome P450 family | 60 | 16,195 | 54% | 84% |
| Intermediate filament family | 75 | 19,566 | 48% | 83% |
| Cytidine and deoxycytidylate deaminase family | 16 | 2,568 | 57% | 83% |
| MHC class I family | 16 | 2,508 | 49% | 83% |
| Inward rectifier-type potassium channel (TC 1,A,2,1) family | 16 | 3,502 | 53% | 83% |
| Organo anion transporter (TC 2,A,60) family | 13 | 5,377 | 59% | 83% |

| | | | | |
|---|---|---|---|---|
| TRAPP small subunits family | 10 | 991 | 60% | 83% |
| Gamma type-C retroviral envelope protein family | 12 | 3,821 | 58% | 82% |
| Gamma-aminobutyric acid receptor (TC 1,A,9,5) subfamily | 19 | 4,647 | 51% | 82% |
| Secretoglobin family | 11 | 573 | 56% | 82% |
| Sulfotransferase 1 family | 32 | 7,292 | 53% | 82% |
| Small GTPase superfamily | 162 | 19,838 | 52% | 82% |
| Classic translation factor GTPase family | 20 | 8,002 | 59% | 82% |
| Cyclophilin-type PPIase family | 21 | 2,989 | 54% | 82% |
| Rab family | 74 | 9,262 | 51% | 81% |
| POU transcription factor family | 16 | 3,537 | 51% | 81% |
| Phosphoglycerate mutase family | 10 | 1,784 | 50% | 81% |
| Sodium channel (TC 1,A,1,10) family | 10 | 8,226 | 43% | 81% |
| TFS-II family | 12 | 1,194 | 48% | 80% |
| NR2 subfamily | 12 | 2,759 | 49% | 80% |
| Peptidase C19 family | 75 | 37,070 | 55% | 80% |
| Recoverin family | 11 | 1,101 | 47% | 80% |
| CK1 Ser/Thr protein kinase family | 12 | 3,458 | 51% | 80% |
| Rho family | 20 | 2,529 | 49% | 79% |
| CEA family | 22 | 3,425 | 40% | 79% |
| MIP/aquaporin (TC 1,A,8) family | 16 | 2,351 | 50% | 78% |
| Annexin family | 14 | 2,809 | 53% | 77% |
| GST superfamily | 23 | 2,818 | 50% | 77% |
| Heat shock protein 70 family | 17 | 5,572 | 49% | 76% |
| Ras family | 26 | 2,524 | 47% | 76% |
| Globin family | 12 | 880 | 49% | 75% |
| Peptidase A1 family | 10 | 2,164 | 51% | 73% |
| KRTAP type 5 family | 11 | 347 | 16% | 72% |
| Alpha/beta interferon family | 16 | 1,072 | 35% | 72% |
| PPP phosphatase family | 13 | 2,470 | 44% | 71% |
| Heat shock protein 90 family | 11 | 2,187 | 37% | 71% |
| G-alpha family | 17 | 2,742 | 40% | 71% |
| KRTAP type 10 family | 13 | 912 | 24% | 71% |
| KRTAP type 4 family | 12 | 409 | 18% | 70% |
| Aldo/keto reductase family | 15 | 1,822 | 39% | 70% |
| Metallothionein superfamily | 14 | 219 | 26% | 70% |
| Type 1 family | 14 | 219 | 26% | 70% |
| Humanin family | 14 | 93 | 27% | 69% |
| Nucleosome assembly protein (NAP) family | 21 | 3,271 | 44% | 68% |
| Actin family | 28 | 4,909 | 44% | 68% |
| Opsin subfamily | 11 | 1,865 | 46% | 66% |
| Neurexin family | 12 | 5,862 | 40% | 62% |

| | | | | |
|---|---|---|---|---|
| LCE family | 19 | 482 | 24% | 61% |
| Cornifin (SPRR) family | 11 | 273 | 29% | 61% |
| KRTAP type 28 family | 10 | 105 | 11% | 60% |
| UDP-glycosyltransferase family | 22 | 3,333 | 29% | 56% |
| Histone H2A family | 19 | 434 | 16% | 50% |
| GAGE family | 27 | 605 | 20% | 43% |
| Tubulin family | 24 | 1,850 | 17% | 42% |
| Histone H2B family | 21 | 475 | 17% | 41% |
| PRAME family | 26 | 2,302 | 19% | 41% |
| SPATA31 family | 14 | 3,114 | 19% | 37% |
| Beta type-B retroviral envelope protein family | 13 | 753 | 10% | 36% |
| NBPF family | 16 | 1,149 | 5% | 35% |
| GOLGA8 family | 15 | 390 | 4% | 33% |
| TAF11 family | 14 | 351 | 13% | 32% |
| HERV class-II K(HML-2) env subfamily | 11 | 297 | 4% | 30% |
| GOLGA6 family | 17 | 933 | 8% | 30% |
| Speedy/Ringo family | 23 | 745 | 10% | 28% |
| HERV class-II K(HML-2) subfamily | 12 | 106 | 6% | 27% |
| Peptidase A2 family | 12 | 106 | 6% | 27% |
| FAM90 family | 21 | 761 | 8% | 25% |
| Beta type-B retroviral Gag protein family | 10 | 481 | 7% | 22% |
| HERV class-II K(HML-2) gag subfamily | 10 | 481 | 7% | 22% |
| USP17 subfamily | 21 | 365 | 3% | 17% |
| NPIP family | 19 | 348 | 3% | 16% |
| Beta type-B retroviral polymerase family | 10 | 429 | 4% | 15% |
| HERV class-II K(HML-2) pol subfamily | 10 | 429 | 4% | 15% |

**Table S8.** Proteins without Tryptic-digest Unique Peptides (TUPs)

| **Accession Number** | **Protein Name** | **Amino Acids number** |
|---|---|---|
| P22103 | Pneumadin | 10 |
| P01358 | Gastric juice peptide 1 | 10 |
| A0A075B706 | T cell receptor delta joining 1 | 16 |
| A0A075B700 | T cell receptor alpha joining 31 | 18 |
| Q9NRI7 | Putative pancreatic polypeptide 2 | 21 |
| P0CJ68 | Humanin-like 1 | 24 |
| P62945 | 60S ribosomal protein L41 | 25 |
| P0CJ69 | Humanin-like 2 | 28 |
| A0A0A0MT76 | Immunoglobulin lambda joining 1 | 42 |
| Q13072 | B melanoma antigen 1 | 43 |
| Q86Y27 | B melanoma antigen 5 | 43 |
| P62891 | 60S ribosomal protein L39 | 51 |
| Q59GN2 | Putative 60S ribosomal protein L39-like 5 | 51 |
| P04553 | Sperm protamine P1 | 51 |
| P35325 | Small proline-rich protein 2B | 72 |
| P22531 | Small proline-rich protein 2E | 72 |
| P61583 | Endogenous retrovirus group K member 5 Np9 protein | 75 |
| P61581 | Endogenous retrovirus group K member 24 Np9 protein | 75 |
| P0DMR2 | Secretoglobin family 1C member 2 | 95 |
| Q9NY87 | Sperm protein associated with the nucleus on the X chromosome C | 97 |
| Q9BXN6 | Sperm protein associated with the nucleus on the X chromosome D | 97 |
| P61571 | Endogenous retrovirus group K member 21 Rec protein | 104 |
| P0DOY3 | Immunoglobulin lambda constant 3 | 106 |
| P0DOY2 | Immunoglobulin lambda constant 2 | 106 |
| P01834 | Immunoglobulin kappa constant | 107 |
| Q9GZM3 | DNA-directed RNA polymerase II subunit RPB11-b1 | 115 |
| Q13066 | G antigen 2B/2C | 116 |
| Q9UEU5 | G antigen 2D | 116 |
| P0DP02 | Immunoglobulin heavy variable 3-30-3 | 117 |
| A0A0C4DH73 | Immunoglobulin kappa variable 1-12 | 117 |
| P01599 | Immunoglobulin kappa variable 1-17 | 117 |
| P0DSO3 | G antigen 4 | 117 |
| Q13069 | G antigen 5 | 117 |
| Q4V321 | G antigen 13 | 117 |
| P0DP04 | Immunoglobulin heavy variable 3-43D | 118 |
| Q5T751 | Late cornified envelope protein 1C | 118 |
| Q5T7P3 | Late cornified envelope protein 1B | 118 |
| P06310 | Immunoglobulin kappa variable 2-30 | 120 |
| P33778 | Histone H2B type 1-B | 126 |

**Table S9.** Profiling and quantification of CrUPs in specialized databases – datasets

| | Cancer Antigenic Peptides (number) | Immune Epitope Peptides (number) | Bioactive Peptides (number) |
|---|---|---|---|
| **Peptides** | 400 | 3,709 | 1,206 |
| **Peptides containing CrUPs** | 355 | 3,230 | 1,174 |
| **Percentage (%)** | 89% | 87% | 97% |

**Table S10.** Analysis of CrUP length in 20 organisms, including model systems

| Organism | Reviewed Proteins (number) | CrUPs (number) | Tetra-peptides (%) | Penta-peptides (%) | Hexa-peptides (%) | Hepta-peptides (%) |
|---|---|---|---|---|---|---|
| **Homo Sapiens (Human)** | 20,422 | 7,271,607 | 0% | 9% | 69% | 21% |
| **Mus musculus (Mouse)** | 17,178 | 6,536,956 | 0% | 10% | 70% | 18% |
| **Arabidopsis Thaliana (A. thaliana)** | 16,351 | 4,295,898 | 0% | 15% | 71% | 13% |
| **Rattus norvegicus (Rat)** | 8,182 | 2,825,568 | 0% | 26% | 66% | 7% |
| **Saccharomyces cerevisiae (Baker's Yeast)** | 6,727 | 2,002,523 | 0% | 33% | 62% | 4% |
| **Bos taurus (Bovine)** | 6,040 | 1,649,223 | 1% | 42% | 54% | 3% |
| **Caenorhabditis elegans (C. elegans)** | 4,456 | 1,641,926 | 1% | 44% | 53% | 2% |
| **Escherichia coli (E. coli)** | 4,357 | 952,021 | 2% | 56% | 41% | 2% |
| **Drosophila melanogaster (Fruit Fly)** | 3,723 | 1,623,848 | 1% | 43% | 53% | 3% |
| **Danio rerio (Zebrafish)** | 3,306 | 1,093,006 | 1% | 55% | 42% | 2% |
| **Gallus gallus (Chicken)** | 2,307 | 766,884 | 2% | 65% | 31% | 1% |
| **Sus scrofa (Pig)** | 1,458 | 390,683 | 8% | 77% | 15% | 0% |
| **Oryctolagus cuniculus (Rabbit)** | 901 | 261,990 | 14% | 76% | 10% | 0% |
| **Zea mays (Maize)** | 851 | 149,362 | 23% | 69% | 7% | 0% |
| **Nicotiana tabacum (Common Tobacco)** | 505 | 80,560 | 40% | 57% | 2% | 0% |
| **Ovis aries (Sheep)** | 466 | 93,208 | 43% | 54% | 2% | 0% |
| **Glycine max (Soybean)** | 431 | 75,995 | 43% | 55% | 2% | 0% |
| **Pisum sativum (Garden Pea)** | 402 | 76,518 | 44% | 53% | 2% | 0% |
| **Triticum aestivum (Wheat)** | 381 | 49,819 | 57% | 40% | 2% | 0% |
| **Equus caballus (Horse)** | 292 | 64,920 | 56% | 42% | 1% | 0% |

**Table S11.** Common proteins are being recognized among Vertebrata or Magnoliopsida (Flowering Plants) group members

| Common Proteins | | | |
| --- | --- | --- | --- |
| **Vertebrata** | | **Magnoliopsida – Flowering Plants** | |
| **Entry Name** | **Protein Name** | **Entry Name** | **Protein Name** |
| ATP6 | ATP synthase subunit a | ATP9 | ATP synthase subunit 9, mitochondrial |
| ATP8 | ATP synthase protein 8 | ATPA | ATP synthase subunit alpha |
| B2MG | Beta-2-microglobulin | ATPB | ATP synthase subunit beta |
| COX1 | Cytochrome c oxidase subunit 1 | ATPE | ATP synthase epsilon chain |
| COX2 | Cytochrome c oxidase subunit 2 | ATPF | ATP synthase subunit b |
| COX3 | Cytochrome c oxidase subunit 3 | ATPH | ATP synthase subunit c, chloroplastic |
| CYB | Cytochrome b | ATPI | Actinia tenebrosa protease inhibitors |
| CYC | Cytochrome c | CB21 | Chlorophyll a-b binding protein 2.1, chloroplastic |
| HBA | Hemoglobin subunit alpha | CYB6 | Cytochrome b6 |
| MYG | Myoglobin | CYF | Cytochrome f |
| NU1M | NADH-ubiquinone oxidoreductase chain 1 | IF4E1 | Eukaryotic translation initiation factor 4E-1 |
| NU2M | NADH-ubiquinone oxidoreductase chain 2 | NDHK | NAD(P)H-quinone oxidoreductase subunit K, chloroplastic |
| NU3M | NADH-ubiquinone oxidoreductase chain 3 | NU5C | NAD(P)H-quinone oxidoreductase subunit 5, chloroplastic |
| NU4LM | NADH-ubiquinone oxidoreductase chain 4L | PETD | Cytochrome b6-f complex subunit 4 |
| NU4M | NADH-ubiquinone oxidoreductase chain 4 | PSAA | Photosystem I P700 chlorophyll a apoprotein A1 |
| NU5M | NADH-ubiquinone oxidoreductase chain 5 | PSAB | Photosystem I P700 chlorophyll a apoprotein A2 |
| NU6M | NADH-ubiquinone oxidoreductase chain 6 | PSAC | Photosystem I iron-sulfur center |
| P53 | Cellular tumor antigen p53 | PSAI | Photosystem I reaction center subunit VIII |
| SODC | Superoxide dismutase [Cu-Zn] | PSAJ | Photosystem I reaction center subunit IX |
| ST7 | Suppressor of tumorigenicity 7 protein | PSBA | Photosystem II protein D1 |
| | | PSBC | Photosystem II CP43 reaction center protein |
| | | PSBD | Photosystem II D2 protein |
| | | PSBE | Cytochrome b559 subunit alpha |
| | | PSBF | Cytochrome b559 subunit beta |
| | | PSBH | Photosystem II reaction center protein H |
| | | PSBJ | Photosystem II reaction center protein J |
| | | PSBK | Photosystem II reaction center protein K |
| | | PSBL | Photosystem II reaction center protein L |
| | | PSBM | Photosystem II reaction center protein M |
| | | PSBN | Photosystem biogenesis factor 1 |
| | | PSBT | Photosystem II reaction center protein T |
| | | PSBZ | Photosystem II reaction center protein Z |
| | | RBL | Ribulose bisphosphate carboxylase large chain |
| | | RPOA | DNA-directed RNA polymerase subunit alpha |
| | | RPOC2 | DNA-directed RNA polymerase subunit beta' |
| | | RR19 | Small ribosomal subunit protein uS19c |
| | | RR7 | Small ribosomal subunit protein uS7cz/uS7cy |

20 reviewed proteins are commonly recognized in the 10 representative Vertebrates included in this study, while 37 proteins are identified in the 6 Flowering Plants, with no reviewed protein in common being detected in between the Vertebrate and Flowering Plant groups

**Table S12**. ATP6 protein identification across Vertebrates, by the UUP entity profiling

| Vertebrata (number) | Vertabrata Reviewed Proteins (number) | Vertabrata ATP6 Protein (number) |
|---|---|---|
| 3,313 | 86,649 | 68 |

| Universal Unique Peptide (UUP) | Proteins Containing the Peptide | ATP6 Proteins Containing the UUP | Other Proteins Containing the UUP | Classification Rate | Identification Rate |
|---|---|---|---|---|---|
| FTPTTQLS | 61 | 61 | 0 | 100% | 89.7% |
| VRLTAN | 65 | 65 | 0 | 100% | 95.5% |
| IQAYVF | 64 | 64 | 0 | 100% | 94.1% |
| FTPTTQLS and VRLTAN and IQAYVF | 56 | 56 | 0 | 100% | 82.3% |
| FTPTTQLS or VRLTAN or IQAYVF | 68 | 68 | 0 | 100% | 100% |

In the 86,648 reviewed Vertebrate proteins, 68 ATP6 proteins are being recognized. Each UUP alone identifies more that 90% (up to 96%) of these proteins. The 3 UUPs simultaneously identify 82% of the ATP6 proteins, while the “FTPTTQLS” or “VRLTAN” or “IQAYVF” individually can identify all the ATP6 proteins of Vertebrate. The “Classification Rate” is the “%” of the proteins that contain the UUP(s) and they belong to the ATP6 family, to the total number of proteins being identified by the UUP(s) in Verterbrates. The “Identification Rate” is the “%” of ATP6 proteins identified by the UUP(s) to the total number of ATP6 proteins in Vertebrates.

**Table S13.** CYB protein identification across Vertebrates by the UUP profiling

| **Eukaryota (number)** | | **Eukaryota Reviewed Proteins (number)** | | **Eukaryota CYB Proteins (number)** | |
|---|---|---|---|---|---|
| 8,286 | | 197,016 | | 1,764 | |
| **Universal Unique Peptide (UUP)** | **Proteins Containing the Peptide (number)** | **CYB Proteins Containing the Peptide (number)** | **Other Proteins Containing the Peptide (number)** | **Classification Rate (%)** | **Identification Rate (%)** |
| HANGAS | 1,597 | 1,591 | 6 | 99.6% | 90.2% |
| PEWYFL | 1,626 | 1,625 | 1 | 99.9% | 92.1% |
| HANGAS and PEWYFL | 1,512 | 1,512 | 0 | 100% | 85.7% |
| HANGAS or PEWYFL | 1,719 | 1,712 | 7 | 99.6% | 97.0% |

In the 197,016 reviewed Vertebrate proteins, 1,764 CYB proteins are being recognized. Each UUP entity alone identifies more that 90% of the CYB proteins. The 2 UUPs simultaneously identify ~86% of the CYB proteins, while the “HANGAS” or “PEWYFL” peptide alone can identify almost all the CYB proteins of Vertebrates. The “Classification Rate” is the “%” of the proteins that contain the UUP(s) and they belong to the CYB family, to the total number of proteins that are being identified by the UUP(s) in Verterbrates. The “Identification Rate” is the “%” of CYB proteins identified by the UUP(s) to the total number of CYB proteins in Vertebrates.

**Table S14.** Searching for CYB proteins by the UUP profiling into the unreviewed proteomes of *Nicotiana tabacum* and *Glycine max*

| ***Nicotiana tabacum*** **(Common Tobacco)** | | | |
|---|---|---|---|
| Total unreviewed proteins | 75,689 | | |
| **Proteins containing the UUP** | | | |
| **"HANGAS"** | | **"PEWYFL"** | |
| **Protein name** | **Entry Code** | **Protein name** | **Entry Code** |
| Cytochrome b | O03194 | Cytochrome b | O03194 |
| Cytochrome b | Q5MA49 | Cytochrome b | Q5MA49 |
| Cytochrome b | Q95871 | Cytochrome b | Q95871 |
| | | Cytochrome b | A0A1S4C7L2 |
| ***Glycine max*** **(Soybean)** | | | |
| Total unreviewed proteins | 84,712 | | |
| **Proteins containing the UUP** | | | |
| **"HANGAS"** | | **"PEWYFL"** | |
| **Protein name** | **Entry Code** | **Protein name** | **Entry Code** |
| Cytochrome b | H8Y6Q3 | Cytochrome b | H8Y6Q3 |
| Cytochrome b | K7KP66 | Cytochrome b | K7KP66 |
| C3H1-type domain-containing protein | A0A0R0FST1 | | |
| C3H1-type domain-containing protein | I1MQ33 | | |

The UUPs are being used for the identification of CYB proteins into the unreviewed proteomes of *Nicotiana tabacum* and *Glycine max*. "HANGAS" UUP mediated for the identification of 3 unreviewed proteins and "PEWYFL" UUP facilitated the detection of 4 proteins, in Tobacco. In Soybean, "HANGAS" UUP led to the recognition of 4 unreviewed proteins, while "PEWYFL" UUP resulted in the characterization of 2 CYB proteins.
